# Structural basis for reprogramming group II intron target specificity

**DOI:** 10.64898/2026.11.13.765053

**Authors:** Nicholas Dayrell-Armes, Frank Bowler, Simona Cugusi, Ian Cook, Nikola Dzhindzhev, Jessica Leigh, Chanakan Lorvoralak, Jonathan Ward, Jamie Elwell, Sara Toral-Perez, Maria Luisa Miranda Vera, Dariia Babych, Steven W. Hardwick, Lee Cooper, Dimitri Y. Chirgadze, Niall A. Armes

**Affiliations:** Gene Weaver Ltd, Building 1, Granta Centre, Granta Park, Cambridge CB21 6AL, UK; Cryo-EM Facility, Department of Biochemistry, University of Cambridge, Tennis Court Road, Cambridge CB2 1GA, UK

## Abstract

Group II introns are mobile ribozymes that, with their intron-encoded protein (IEP), insert into duplex DNA without a double-strand break—a route to programmed gene insertion so far confined to bacteria. How class IIA complexes recognize and remodel duplex DNA was unknown. We identify group IIA intron–IEP pairs that retrohome efficiently into linear, relaxed duplex DNA, and report cryo-EM structures of one, Voyager-1, at successive stages of insertion. Major- and minor-groove peptides clamp the 5′ target and prime melting; intron nucleotides capture one strand; and a peptide linker threads between the re-annealing strands, reading 3′ sequence and trapping the HNH nuclease in a nicking-competent position. Guided by these structures, we reprogram protein-encoded specificity, yielding combined RNA- and protein-encoded recognition sufficient, by sequence, for single-site targeting in complex plant and animal genomes. Voyager-1 functions when formed in mammalian cells without Mg²⁺ supplementation, overcoming a long-standing barrier to class IIA introns in eukaryotes.

**Highlights:**

- Cryo-EM captures a group IIA intron–IEP complex invading linear duplex DNA
- Peptides clamp target DNA and a linker positions the HNH nuclease
- The protein-encoded specificity is reprogrammable
- Targeting complexes are highly active in mammalian cells

## Introduction

Group II (GII) introns are structured ribozymes that encode a mobilisation protein, the intron-encoded protein (IEP), within an internal open reading frame^1^ (Figure 1). They are the ancestors of spliceosomal introns, and the IEP is the ancestor of the catalytic core of the spliceosomal protein PRP8^2–4^. GII intron RNAs position the nucleotides and catalytic residues for splicing by spontaneous folding of RNA helices and tertiary interactions, both Watson-Crick and non-canonical^5–7^. Remarkably, the excised intron in complex with its IEP can recognise a specific duplex DNA target and integrate into it by reverse splicing coupled to template-primed reverse transcription (TPRT), a process termed retrohoming^8–10^ (Figure 1). Orchestrating this DNA insertion appears to be the principal function of the IEP, since many intron RNAs execute forward splicing independently^11^.

**Figure 1.**
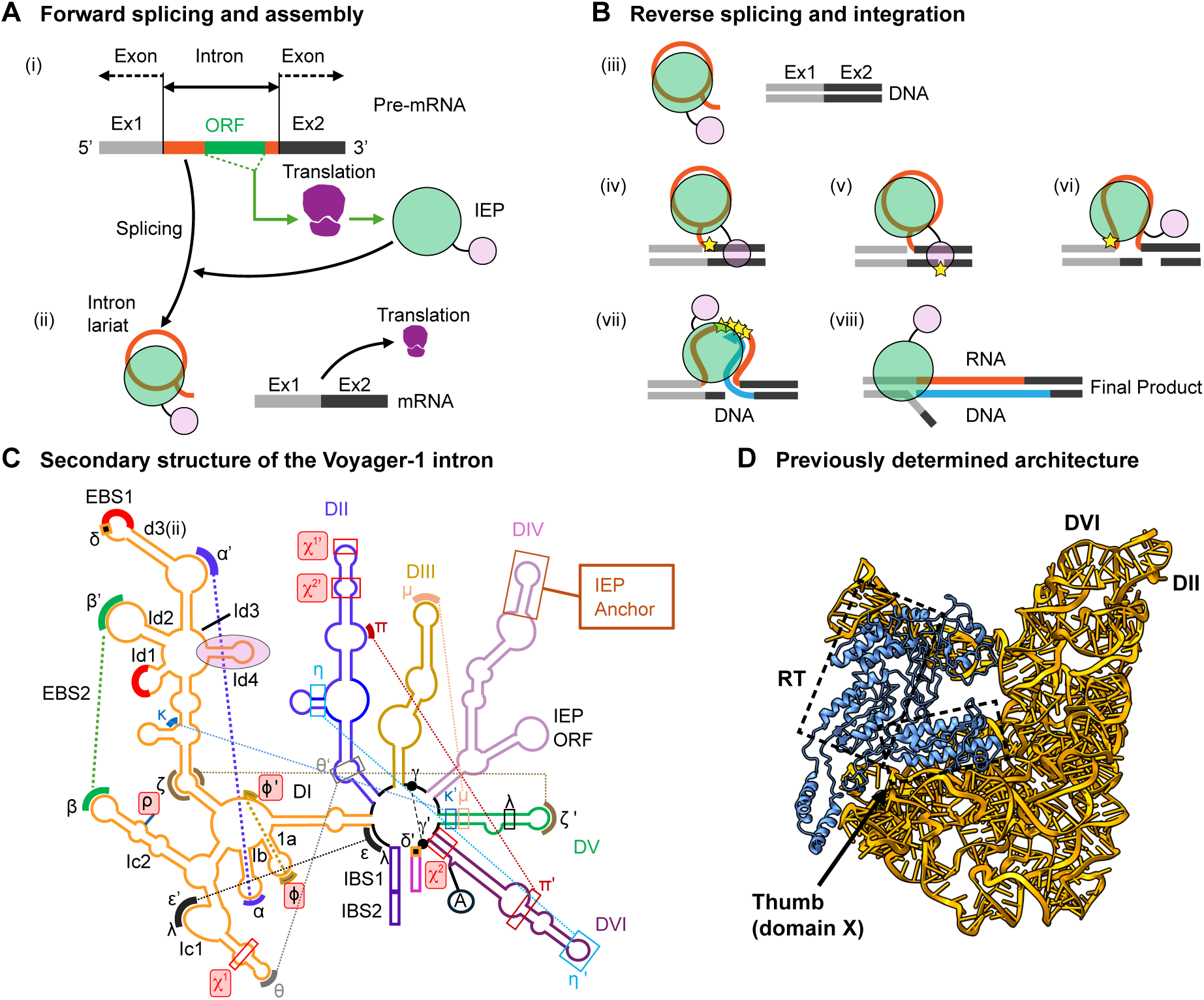
The group II intron retrohoming cycle and the Voyager­1 intron. (A) Forward splicing and assembly. The IEP is translated from an open reading frame within domain IV of the pre­mRNA (i); the intron excises as a lariat and binds the IEP to form the mobile ribonucleoprotein (ii). (B) Reverse splicing and integration. The ribonucleoprotein binds a duplex target (iii) and reverse­splices the intron into the top strand (iv); HNH nicks the opposite strand (v) and reverse splicing completes (vi), freeing a 3′ end that primes reverse transcription (vii) to give the integrated product (viii). (C) Secondary structure of the Voyager­1 intron, domains I–VI coloured separately, with EBS1/EBS2, IBS1/IBS2, δ, the branch adenosine (A) and the IEP anchor marked. Greek letters denote tertiary interactions; those highlighted in red (φ, χ1, χ2 and ρ) are proposed here (Supplementary Note 2). (D) Previously determined group II intron architecture (PDB:8H2H), shown for orientation.

GII introns were recognised as a gene-targeting platform well before CRISPR and the current generation of break-free insertion technologies ^12–14^. Heterologous cargo can be carried within the body of a retargeted intron^15^, and target recognition draws on both RNA–DNA and protein–DNA contacts across a combined footprint of up to ∼35 bp^16,17^. Yet successful application has remained largely confined to microbial systems^13,15^, with eukaryotic use proving more difficult^12,14,18,19^.

Two gaps are conspicuous. First, while the programmable RNA–DNA component of target recognition has been defined and exploited to retarget introns by RNA engineering^16,20,21^, the protein-encoded component is ill-defined outside the IIC class^3,22^. Bacterial GII introns fall into classes IIA, IIB and IIC^1,23,24^; IIA systems being the most elaborate, carrying a C-terminal HNH endonuclease on the IEP to nick the target for TPRT^9,25^, an additional intron RNA loop (DId4), and expanded RT-domain insertions RT4a and RT7a^22,26^. No structure has captured a IIA intron during retrohoming, and in the one reported structure of a IIB intron reverse splicing into DNA the target duplex was largely disordered^27^, leaving duplex recognition, melting and nuclease positioning unresolved for both classes.

Second, functional understanding of class IIA rests almost entirely on a single system, LtrA^10,16,17,28,29^. LtrA protein is difficult to purify alone in soluble form, has a short half-life at 37°C^28^, and has often been used with high complex-to-target ratios — 50:1 in the only quantified in vitro retrohoming reaction^30^, and 48:1 and 1000:1 in other mechanistic studies in vitro^28,31^. Studies exploring LtrA activity in eukaryotic cells showed limited success, and authors proposed inadequate free Mg²⁺ concentrations as one possible explanation^18,19^. A concern is that general properties of group II introns, and this class in particular, may not have been readily separated from the characteristics of a single member. Other examples should consequently be explored to establish a generality to this proposal.

We therefore screened broadly for intron–IEP pairs that retrohome efficiently into linear, relaxed duplex DNA in vitro — deliberately selecting against any dependence on supercoiling or replication-derived single-stranded substrates. Here we define cryo-EM structures of one such class IIA system, Voyager-1, bound to target DNA in successive configurations during integration, and use them to identify and reprogram the protein–DNA interfaces that set target specificity. Testing their activity in mammalian cells shows that the earlier claimed requirements for supplemented Mg²⁺ may be confined to the LtrA system, and not a fundamental constraint for IIA class systems.

## Results

### Highly active class IIA complexes retrohome into linear duplex DNA

GII IEPs have been reported to be difficult to produce as soluble recombinant proteins without their cognate RNA or a heterologous solubility partner, as well-documented for LtrA^10,28,32^. We cloned and expressed in *E. coli* a panel of class IIA and IIB IEPs, applying no in silico pre-selection. A subset purified with or without an N-terminal His tag, retained polymerase activity, and were screened for IEP-dependent forward splicing and retrohoming. Voyager-1 and Voyager-2, two particularly active IIA candidates, along with several other IEPs we identified, are the first full-length, HNH-containing, retrohoming-competent class IIA IEPs reported as soluble, stable and active without a larger fusion or partner RNA (Supplementary Note 1).

Voyager-1 is a class IIA intron–IEP pair from a *Bacillus mycoides* plasmid (CP092835.1; intron 57,694–60,210; IEP ORF 58,276–60,099); Voyager-2, a related system from a *Bacillus cereus* plasmid (CP053963; intron 1,153–3,681; IEP 1,738–3,576), corroborates key structural and reprogramming findings. Introns were derivatives of the natural sequences with partial or complete deletions of the domain IV ORF and, in some cases, recoded EBS1 or EBS2 (Figure S2 and Table S1; STAR Methods).

Linear duplex targets for retrohoming spanned −1 to −25 and +1 to +13 relative to the splice site, with all proposed RNA–DNA interactions adjusted to Watson–Crick pairing^16,20^ (Figure S10). Integration was quantified by four independent routes — gel shift, endpoint qPCR, a real-time fluorescence assay, and nanopore sequencing, which reads integrations directly and so reports target-sequence preference (Figure S3; STAR Methods). Retrohoming was efficient at low complex-to-target ratios (Figure S16) — e.g. 4:1 at 100 nM target (Figure S3C) — with full-length retrohomed products appearing within 10 min of initiation. Retrohoming was also detectable at relatively low Mg²⁺ concentrations suggesting that free ion concentration in a cellular context may not be a gating limitation (Figure S17). Overall this system appeared well suited to structural work.

### Cryo-EM structures of Voyager-1 during retrohoming

Intron RNA, IEP and target duplex were assembled under modified retrohoming conditions and vitrified, with dNTPs omitted to prevent polymerase activity (Figure S1; STAR Methods). Three classes of complex were resolved. Named by configuration and abundance: the TPRT-primed complex (state 1), in which the primed template occupies the polymerase cleft; the nicking-competent complex (state 2); and the domain VI-down complex (state 3). Detailed models are presented for states 1 and 2; state 3 is described at lower resolution. Target positions are numbered relative to the splice site, −1 and +1 denoting the nucleotides immediately preceding and following it.

#### The TPRT-primed complex (state 1)

Duplex target DNA enters on the 5′ side of the splice site and exits on the 3′ side at roughly 90° to the entry path, emerging from the RT catalytic cleft (Figure 2A). The expected EBS1/IBS1 and EBS2/IBS2 RNA–DNA pairings are present ^5,16,33^, and IEP contacts to the duplex lie upstream. Two distinguishing class IIA IEP features — the 4a and 7a insertions — engage opposite faces of the DNA: 4a is a helix-turn-helix (HTH) motif appended to the RT body by peptides we term hinges (HG) and binds the major groove, while 7a is a short loop we term the beta finger (BF) and binds the minor groove. The DNA is B-form duplex from −14 outward, whereas the −13 sense base is unpaired, projecting into the pocket adjacent to the EBS2:IBS2 interaction.

**Figure 2.**
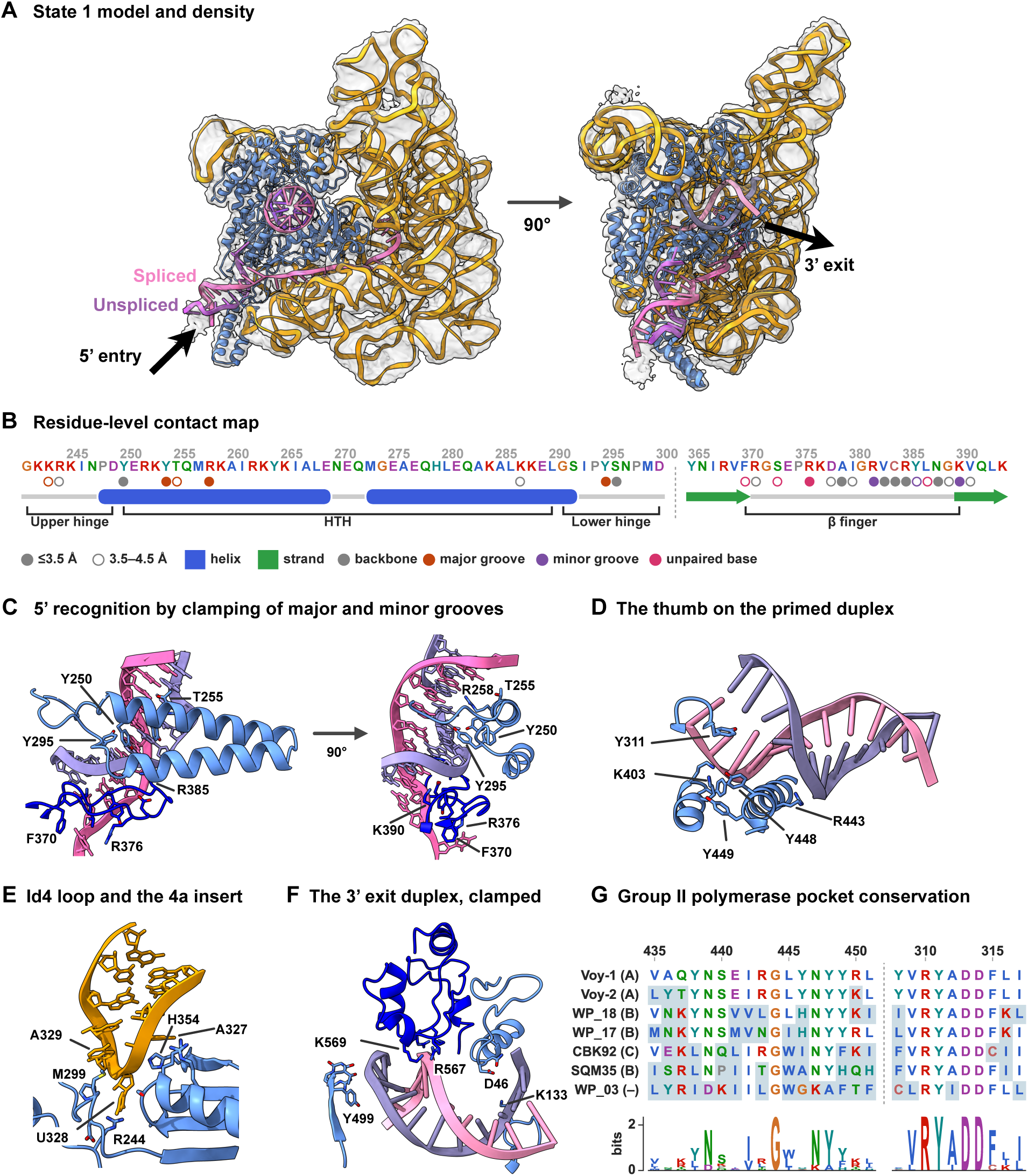
Cryo­EM structure of Voyager­1 engaging a linear duplex target in the TPRT­primed state (state 1). (A) Map and model in two orthogonal views; protein blue, intron RNA gold, target DNA pink (spliced strand) and purple (unspliced strand). The duplex enters on the 5′ side of the splice site and exits at approximately 90° from the reverse transcriptase cleft. (B) Residue­level contact map for the upper hinge, HTH, lower hinge and β finger. Filled circles mark contacts ≤3.5 Å, open circles 3.5–4.5 Å, coloured by contact type; secondary structure is drawn beneath. (C) 5′ recognition from opposite faces of the duplex: the HTH and hinges (Y250, T255, R258, Y295) read the non­spliced­strand backbone and major groove, while the β finger (F370, R376, R385, K390) binds the minor groove opposite. (D) The thumb on the primed duplex, contacting the sugar–phosphate backbone only (K403, R443, Y448, Y449), with Y311 of the YADD motif positioning the primer 3′­OH. (E) The domain I d4 loop (A327–A329) and the 4a insert, engaged by R244, M299 and H354. (F) The 3′ exit duplex enclosed by the RT0 insert, the DNA­binding prong and HNH (D46, K133, Y499, R567, K569), which form a clamp over roughly 10 bp rather than a catalytic arrangement. (G) Conservation of the polymerase pocket, the thumb region and the YADD motif, across seven group II IEPs, as alignment and frequency logo; parentheses give intron class, or – for the ORF­only element G2L4.

The class IIA-specific domain I d4 loop (residues 322–334) forms an integral structure with the IEP 4a insert (Figure 2E). Three nucleotides at the tip — A327, U328 and A329 — are contacted by converging protein segments: R244 of the upper hinge, P298–K304 of the lower hinge, K152 and the highly conserved W153, K323 and R327, and H354. The d4 loop co-ordinates the HTH and hinge segments; we infer that this RNA–protein co-innovation underpins IIA target engagement.

Elsewhere in the intron, IEP residue W418 π-stacks with intron base A226 immediately preceding the five EBS2 residues; an aromatic is conserved at this position in most class IIA IEPs including LtrA. The branchpoint adenosine 2′-hydroxyl lies adjacent to the 5′ phosphate of intron residue G1, geometry consistent with an intact 2′–5′ branch^6,7^, and the branch site is entirely RNA-encapsulated, the nearest protein atom (R479) lying 13 Å away. Other intron tertiary and intron–protein interactions not previously reported are described in Supplementary Note 2.

In state 1 the intron 3′ end and the DNA splice site lie in proximity, blurring whether the first reverse splicing reaction has occurred or exists in equilibrium with its precursor. The second reverse splice, however, is not represented: the first two nucleotides of the intron 5′ end bend upward toward the branchpoint, indicating that the covalent bridge between G1 and the domain VI branchpoint adenosine remains intact. This is consistent with gel-based data showing no detectable fully integrated intron RNA when dNTPs were omitted (Figure S3Cviii), suggesting such intermediates are rare at steady-state under these conditions.

#### The HTH and beta finger clamp the 5′ duplex

We define G241–D249 as the upper hinge (UHG), G291–D300 as the lower hinge (LHG) and Y250–L290 as the HTH (Figure 2C; Figures S8, S9, S11). These segments contact the non-spliced-strand backbone and the adjacent major groove from approximately −17 to −22. Y250, referred to as the reference tyrosine (RefY), is well conserved and contacts the backbone at −18; Y254, T255 and R258 contact the major groove between −19 and −22; and Y295 of the lower hinge reaches both backbone and major groove at −17/−18 (Figure 2B). The BF (F370–K390) binds the minor groove on the opposite face (Figure 2B). Its distal segment (R376–C384) projects into the minor groove between −16 and −18, where R382 makes the only base contacts — to C−17 and G−18 and their partners — while the remaining distal residues contact sugar and phosphate oxygens only. R385 and N388 extend this backbone scaffold across −16 to −18; the proximal surface formed by Y386, L387 and K390 engages the final duplex base pairs at −14/−15. The highly conserved R385 side chain crosses the backbone to contact the hinges near G291–G293 (Figure S12), suggesting a common clamping role.

These peptides grip the DNA from opposing sides, the 4a helices projecting into the major groove while the BF clamps from the minor-groove face, registering the duplex to present the adjacent EBS2 bases for sampling. Modelling an extended duplex further 3′ of −14 suggests a possible collision/interference with the RT 2a insert, potentially providing a mechanism to drive initial helix distortions in this region. At −13 the sense-strand base is flipped toward the IBS2:EBS2 cavity, stacked against the EBS2 loop and coordinated by F370 and R376.

Non-spliced-strand nucleotides 5′ of the fork are disordered in state 1, supporting release of the nicked single-stranded segment from −13 to +8. We confirmed that this strand is indeed accessible for hybridisation and ligation using an adapter followed by nanopore sequencing (Figure S7). Additional contacts to this strand are identified in state 2 below. Variation in hinge, HTH and BF sequences is a prime candidate for protein-encoded 5′ recognition, which we test below.

#### EBS/IBS pairings and the −7 base stack

Intron residues 227–231 pair with DNA −8 to −12, and residues 286–291 pair with −1 to −6 (Figures S4 and S10). The delta base, intron G285, pairs with DNA +1; +2 to +8 of the spliced strand are disordered in state 1, with no clear interaction between +2 and intron A284. Between the two EBS:IBS pairings the −7 base projects in opposition to its neighbours with a pronounced backbone kink and stacks with intron G153. The RNA twists to present this guanine, which likely accounts for the conservation of an unpaired CG dinucleotide in the domain 1C2 stem.

#### The 3′ duplex occupies the RT pocket and is clamped by RT0, the thumb and HNH

On the 3′ side, state 1 contains a duplex comprising both the sense and the non-spliced strand, nicked between +8 and +9 and thereby presenting a free 3′ end in the RT pocket (Figures 2A, 2G). Complexes with duplex positioned in an IEP polymerase pocket have not previously been visualised for IIA and IIB complexes, nor during the act of retrohoming for any IEP. A synthetic RNA:primer bound to RT catalytic cleft has previously determined for a thermophilic class C IEP^22^ (PDB:6AR1). The conserved YADD motif (Y311–A312–D313–D314) sits at the expected incorporation site^22,26^, with the primer 3′-OH less than 3 Å from D313 and Y311. D313, D314 and D158 form the carboxylate cluster expected to coordinate the catalytic metals; neither metal nor nucleotide is resolved, as expected without dNTPs. The thumb contacts the duplex exclusively through the sugar–phosphate backbone, with a bias toward the priming strand.

Many residues are highly conserved among GII IEPs^22,26,34^. In Voyager-1 the single-stranded template bridging the duplex to the splice site (+2 to +8) is poorly resolved, but the corresponding Voyager-2 state resolves it passing through a channel formed by the RT0 finger on one side and the RT1 finger and the turn preceding helix 9 on the other, an arrangement likely general given the conservation of these regions (Figure S4), and largely consistent with the earlier structure with synthetic template (PDB:6AR1).

Three elements thus enclose the duplex. The RT0 insert projects into the major groove between N45 and M51 (Figure 2G; cf. Stamos et al.^22^). The DNA-binding (DBD) prong contacts the templating strand through Y499–R503^25^. The HNH domain, well resolved for residues 540–606, contacts the N-terminal RT region and the non-spliced-strand backbone, and bridges to intron domain VI through N576 and E606. Critically, its active-site residues lie far from the DNA backbone, so HNH is not positioned for catalysis in state 1; with RT0 and the DBD prong it instead forms a clamp covering roughly 10 bp of duplex, thus proposing a second and previously unanticipated role.

#### State 1 is a reversible intermediate off the synthesis pathway

Substantial reverse transcription requires both reverse-splicing reactions, or synthesis would collide with the branchpoint; omitting dNTPs might therefore logically trap the doubly reverse-spliced state. We did not, however, detect it in the TPRT-primed state 1 configuration. We propose instead that state 1, which cannot itself support processive synthesis, exists in equilibrium with a state 3-like conformation in which domain VI and possibly HNH are repositioned, and that this doubly reverse-spliced state is transient and reversible, while the domain VI-in state 1 is stabilised by multiple RNA–RNA contacts and so readily accumulates. Consistent with this, limited RT walking in a gel-based assay indicated that synthesis of only a few nucleotides shifts the equilibrium toward the doubly reverse-spliced state (Figure S3Dii), in agreement with earlier proposals^8,28^.

### The nicking-competent complex – state 2

State 1 revealed little about pre-catalytic engagement of the 3′ target or the position of the non-spliced strand in the melted segment. Particles were collected with an alternative conformation with a different DNA exit path, incompatible with polymerase-competent geometry (Figure 3A). We term this the nicking-competent complex (state 2). Contacts on the 5′ side mirror state 1, with additional contacts visible from the RT2a insert to the non-spliced strand, and the path of the non-spliced strand is now visible, although could not be modelled in parts due to the resolution limits and the convoluted, tightly coiled conformation of the strand. The HNH domain is resolved in a new position, no longer contacting the N terminus or the exiting duplex but instead at the expected catalytic location.

**Figure 3.**
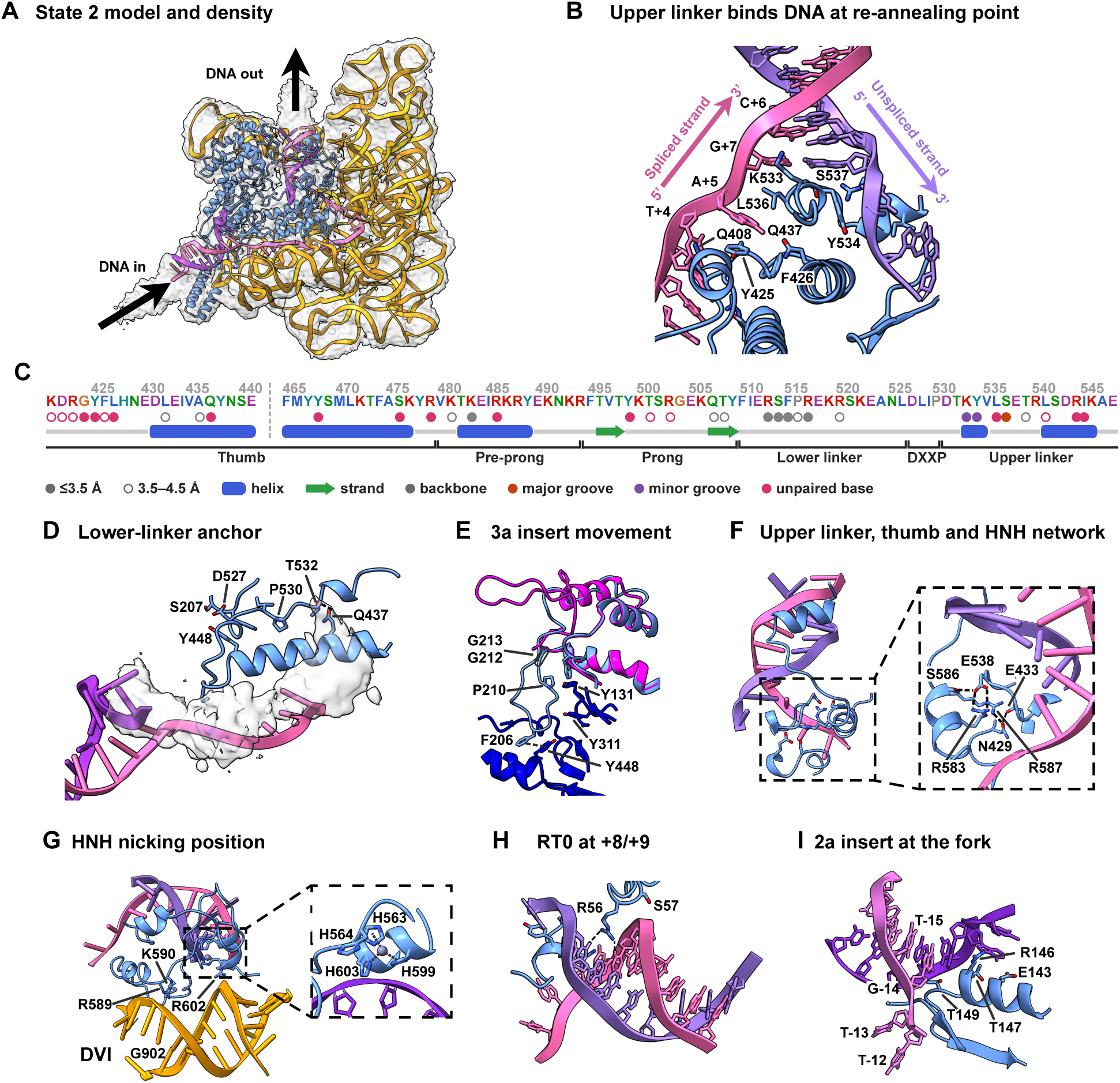
The nicking­competent complex (state 2) resolves both strand paths and the upper linker threaded between them where they re­anneal. (A) Overall map and model; arrows mark DNA entry and exit. (B) The upper linker at the re­annealing point: K533, Y534, L536 and S537 against the unpaired bases T+4 and A+5 and the re­formed duplex at C+6 and G+7, with the thumb residues Q408, F426 and Q437. (C) Contact map for the thumb, pre­prong, prong, lower linker, DXXP motif and upper linker; symbols as in Figure 2B. (D) The lower­linker anchor: D527 contacts S207, with Y448, P530, T532 and Q437 completing the interface. The non­spliced strand is shown as a density cloud, not modelled in detail. (E) Movement of the 3a insert between states, with state 1 (Y131, F206, P210, G212, G213, Y311, Y448) overlaid in pink. (F) The network linking upper linker, thumb and HNH: E538 with R583 and S586, and R587 reaching N429 and E433. (G) HNH in the nicking position between the 3′ exit duplex and intron domain VI; R589, K590 and R602 face the DNA, with G902 of domain VI adjacent. Inset, the histidine cluster (H563, H599, H603) around the DNA­proximal of two zincs (grey sphere). (H) RT0 contacts at +8/+9 through R56 and S57. (I) The distal 2a insert at the fork, with E143, R146, T147 and T149 at the flipped T−13 and the −15 to −12 backbone.

#### DNA strand paths

The spliced strand mirrors its state 1 path to the splice site, then turns sharply and passes over the top of the thumb (Figures 3A, 3D). The +1 base pairs the delta base as expected and +2 may interact weakly with intron delta−1. Bases +3, +4 and +5 are unpaired and project toward the protein: +3 toward S476 and R479, while +4 and +5 are sandwiched between thumb residues G424, Y425 and F426 on one face and HNH residues K577, S578 and F579 on the other. From +6 the strand re-forms duplex with the non-spliced strand. The melted segment therefore extends from −13 to +5.

The non-spliced strand, completely delocalised in state 1, can now be traced. Following duplex separation at −13 it takes a path along the thumb before re-forming duplex at +6. The arrangement of its first unpaired bases was unexpected: the −13 base — necessarily a purine in Voyager-1 — is flipped out toward the backbone at Q142/E143 and π-stacks with −11, while −12 twists back to stack with sense-strand −14 and contacts R146. Beyond this a portion of single-stranded DNA is coiled and contacts K519 and R520 before extending toward the DBD to contact Y499 (Figure 3B). Shortly before duplex reformation the backbone at +4 and +5 is guided through a channel formed by R544 and I545 on one side and Y534 on the other (Figure 3D). R544/I545 are conserved in class IIA IEPs, suggesting a scaffold that organises this region.

#### A ratchet precedes a peptide linker threading between the re-annealing strands

Between the last thumb helix and the HNH domain we define V480–R494 as the pre-prong, F495–F510 as the DBD prong, I511–L526 as the lower linker (LL) and P530–R549 as the upper linker (UL); the three residues preceding P530 form the DXXP motif, conserved in many class IIA IEPs (Figure S8). Together these elements were earlier described collectively as the DNA-binding, or D, domain^17,25^.

In state 2 the thumb has wedged between the separated strands and, strikingly, the HNH domain has penetrated the duplex. The LL and UL first run parallel to the thumb and beneath the non-spliced strand, then turn and cross the top of the thumb (LL) before passing between the two DNA strands (UL) just below where they re-zipper. Proximally, P517 and R518 contact the sense-strand backbone at −11 and −12 before the chain turns upward to engage the disordered strand through K519, R520 and K522 (Figure 3B).

N525–D527, just preceding the DXXP motif, forms a multipartite anchor: N525 contacts the RT0 backbone between A48 and L49, L526 contacts H138, and D527 of the DXXP motif binds tightly to T205, F206 and S207 of the 3a insert, whose F206 ring π-stacks with the conserved thumb residue Y448. In state 1 the 3a insert is displaced far from this position (Fig 3F). We infer one role of the 3a insert is to stabilise the linker path and occlude the polymerase DNA-binding pocket during early remodelling, establishing a ratchet against DNA re-association.

Distally, upper-linker residues K533, Y534, L536, S537 and E538 contact or guide the reforming helix at +5, +6 and +7 (Figure 3D). These interactions have no counterpart in earlier structural work or in AlphaFold predictions of the isolated protein^35^.

This confirms that the previously defined DNA-binding domain contacts target DNA^17,25^, but localises specific contacts to DNA 3′ of the splice site, where they serve in part to position HNH for backbone attack.

#### HNH is trapped in a nicking-competent position

We define HNH as C550 to the C terminus. In state 2 it is sandwiched between the 3′ exit duplex and intron domain VI. Class IIA HNH domains carry a longer peptide between the di-histidine motif (H563, H564) and the CXXCH motif (C595, C598, H599) than their IIB counterparts, which we term the HNH insert (Figure S13). H563, H599 and H603 coordinate a zinc 4.3 Å from the DNA backbone; H564 lies close to the backbone between +8 and +9, as expected for incision there (Fig 3).

Positioning appears to combine gross steric trapping — the domain having passed through the helix with the thumb segments — with an array of specific contacts (Table 1). The C-terminal residues R602–E606 contact domain VI RNA near G943–G945 and the insert region M584–K590 contacts the RNA backbone at 900–902. The insert and core pack densely against the upper linker: T539 stabilises the conformation of H564, R583 forms a bidentate salt bridge with E538, F579 forms a hydrophobic interface with L536, and R587 packs along V535–L541. The network extends into the thumb, where R583 and M584 pack against the imidazole of H428 and R587 acts as a hub bridging HNH, linker and thumb through N429 and E433.

HNH is subject to strict spatial and conformational constraint: tethered near the target by a short linker spanning the re-forming duplex, and positioned by concerted contacts with thumb, linker and intron. This four-way coupling supports that the catalytic domain has little autonomous DNA affinity, limiting uncontrolled incision, and predicts that disrupting these networks may affect nicking efficiency or specificity.

### A domain VI-down state, state 3

A lower-abundance class was distinguished by altered morphology of intron domains VI and II (state 3, the domain VI-down state). Particle numbers restricted resolution and we present no atomic model, but combining particles across experiments gave a 5.11 Å map from 7,031 DNA-bound particles (state 3C) in which domain VI is rotated by at least 90° relative to the other states (Figure S5); we present this as preliminary, and subject to appropriate cautious interpretation. The domain II extension is unresolved, consistent with increased mobility, and the resolved region shifts from A401 onward, supporting that identified domain II–domain VI contacts stabilise the VI-up state. The new position of domain VI brings the branchpoint, intron 5′ end and sense-strand DNA 3′ end into close proximity.

Docking state 1 or state 2 models suggests most other regions are reproduced, the HNH domain mobile, and a large proportion of particles with the LL contacting the 3a insert through D527, density further C-terminal being absent. We did not observe the state 2 non-spliced-strand density, nor density for a second strand passing through the upper-linker region, consistent with a transient reverse-splice-2 intermediate with upper linker and HNH released from their constrained positions and the 21-nt non-spliced segment freed. Presence of the 3a insert–lower-linker assembly, with no HNH docked in the synthesis pocket, suggests this state may be incompatible with the HNH-clamped geometry of state 1.

Introducing EBS1:IBS1 mismatches to impede melting while preserving early DNA contacts shifted the population almost entirely to state 3-like classes, raising the possibility that a similar configuration describes earlier forms (Figure S6; Supplementary Note 3). We use this to build a stepwise model for future confirmation.

### A model for retrohoming

We present a stepwise model (Figure 4) in which several transitions are inferred and require further confirmation. We propose DNA scanning occurs in state 3 to minimise steric conflict between B-form DNA and intron domains II and VI. The low-resolution state 3 complexes imply initial DNA contacts are made by the tip of the DBD prong and possibly the end of the beta finger. The HNH domain appears to be initially docked to the N terminus (Figure S5).

**Figure 4.**
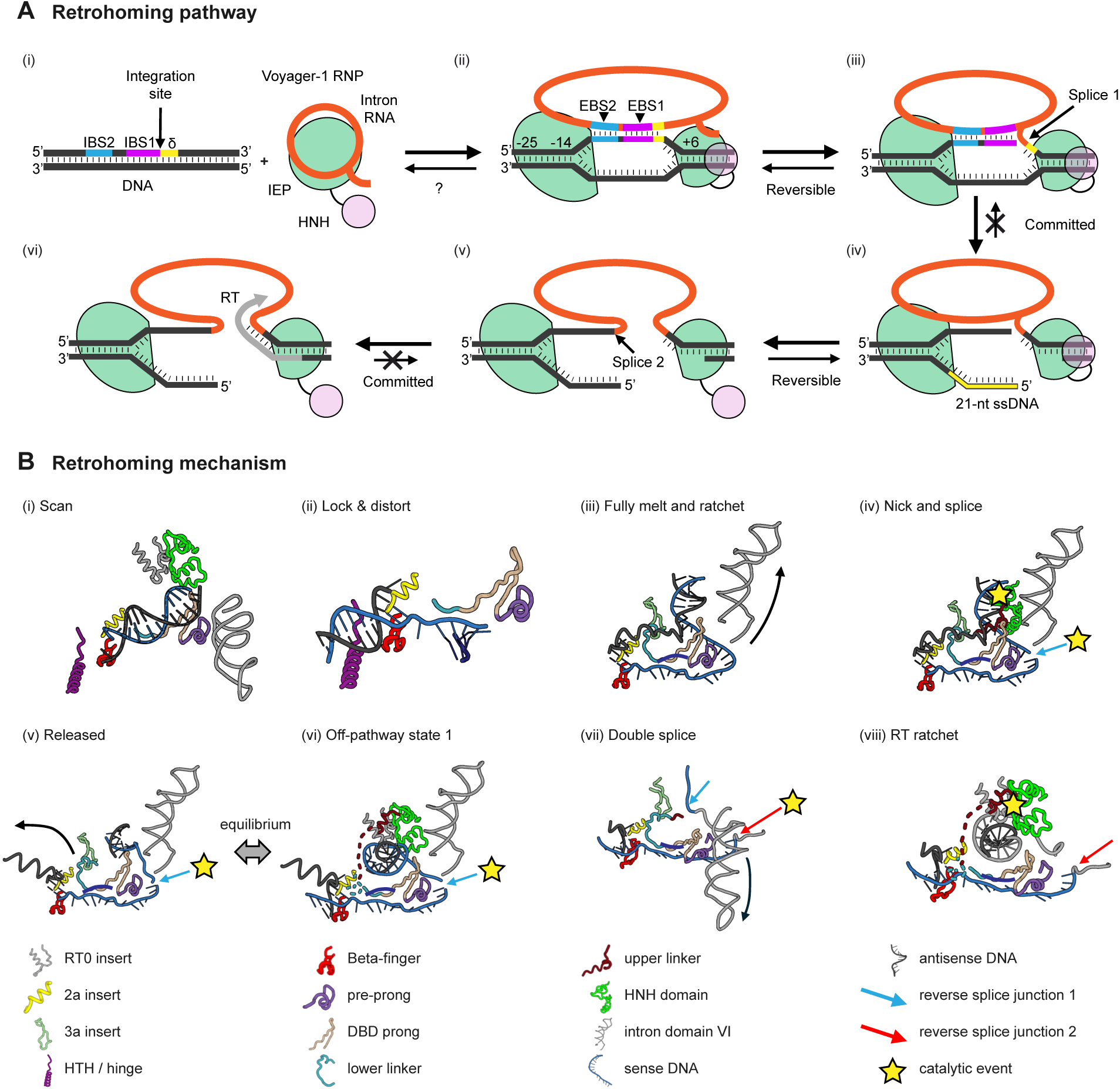
A model for retrohoming. (A) Reaction pathway. The ribonucleoprotein engages the duplex target through IBS2, IBS1 and the δ base (i) and clamps it from −25 to +6, melted from −14 (ii); whether this first step is reversible is not established (?). Reverse splicing at junction 1 commits the complex (iii). Nicking releases a 21­nt single­stranded segment whose 5′ flap shares 8 nt with the cDNA subsequently synthesised (iv); reverse splicing at junction 2 completes intron integration (v), and the freed 3′ end is then extended by the reverse transcriptase (vi). Reversible, committed and equilibrium steps are marked as shown. (B) The same reaction mapped onto the structures, in eight steps from scanning to reverse transcription. State 1 is placed off­pathway, in equilibrium with the released state (steps v and vi). Keys to elements, reverse­splice junctions and catalytic events are given on the figure.

Next, sequence-specific engagement of the HTH, hinge, beta-finger and distal 2a elements arrests scanning and redirects the trajectory of the adjacent duplex: when repositioned into the 5′ protein clamp, B-form duplex DNA likely clashes with the proximal beta finger and 2a insert helix, and this strained interaction likely destabilises the duplex near −13, where EBS2 captures the flipped bases; the DBD prong inserts between the separating strands and the lower linker threads around with stabilising contacts to the antisense ssDNA to anchor the 3a insert through its DXXP motif to form a ratchet opposing reannealing. Melting propagates from −7 to +5 as the sense strand slides along the basic surface preceding the thumb prong, with intron G153 stacking the −7 base and EBS1 capturing further nucleotides.

The HNH and the UL pass through the gap between the separated strands to form the nicking-competent complex. Detachment of the HNH domain from the N terminus has likely occurred in an earlier state 3 form, and DNA binding to the antisense strand may aid the final arrangement. Nicking is the committing event: melting of +6 to +8 releases the non-spliced strand from −13 to +8, allowing the adjacent duplex to rotate into the polymerase pocket and freeing the upper linker and HNH to rearrange. The complex can then either reversibly adopt a quasi-stable state 1 clamp with domain VI up or proceed through a transient domain VI-down conformation to reverse splice 2; DNA synthesis acts as a ratchet, each added nucleotide making reversal of either splicing reaction less accessible; once eight nucleotides have been translocated, the enzyme switches to processive reverse transcription and eventually unwinds the intron.

### Protein-encoded target specificity is extensively reprogrammable

Integration sequencing read target preferences directly. These mirrored the natural residence sequences of Voyager-1 relatives, grouped by conserved proximal beta-finger residues and EBS2 sequence (Figure S14). On the 5′ side, positional base counts gave reproducible preferences and consensus sequences — consistent with the structures, where 5′ DNA is held in a single register from the duplex-binding surfaces through the fork at −13 to the splice site. On the 3′ side the preferred +4 YAYGY motif was frequently recovered shifted by one, occasionally two, nucleotides, confounding conventional logo analysis (Figures S3F and S19); we therefore grouped sequences by pairwise correlation before generating logos (STAR Methods), resolving distinct patterns despite register shifts.

The natural diversity of class IIA sequences at these interfaces should imply a corresponding breadth of programming potential, so we curated class IIA intron and IEP sequences into homology groups, assigned host target sites where possible, and used these to select peptide-exchange points for hybrids on the Voyager-1 backbone.

### 5′ recognition is set by the HTH, beta finger and 2a insert

Replacing G241–M299 (both hinges and the HTH) with blocks from diverse class IIA clades yielded hybrids whose base preference across −17/−18 to −24 changed substantially, often extending the programmed footprint further 5′ than Voyager-1 (Figure 5D). Several hybrids selected beyond −21, the most 5′ position under substantial selection in Voyager-1, consistent with some earlier reports of broader footprints^16,17^ and showing that both strength and extent of selection are tuneable. The libraries used in this study only extended as far as −24 from the splice site, further 5’ to the selectivity observed for Voyager-1 as well as the contacts observed in the structure for wild type protein. We cannot however exclude that some hybrids or native proteins could extend contacts to −25, or even beyond, which should be tested in future. Nonetheless HTH and hinge exchanges left −15/−14/−13 unaltered, still dominated by the Voyager-1 TGY motif, confirming the structural inference that fork-proximal positions are encoded elsewhere (Figure S18).

**Figure 5.**
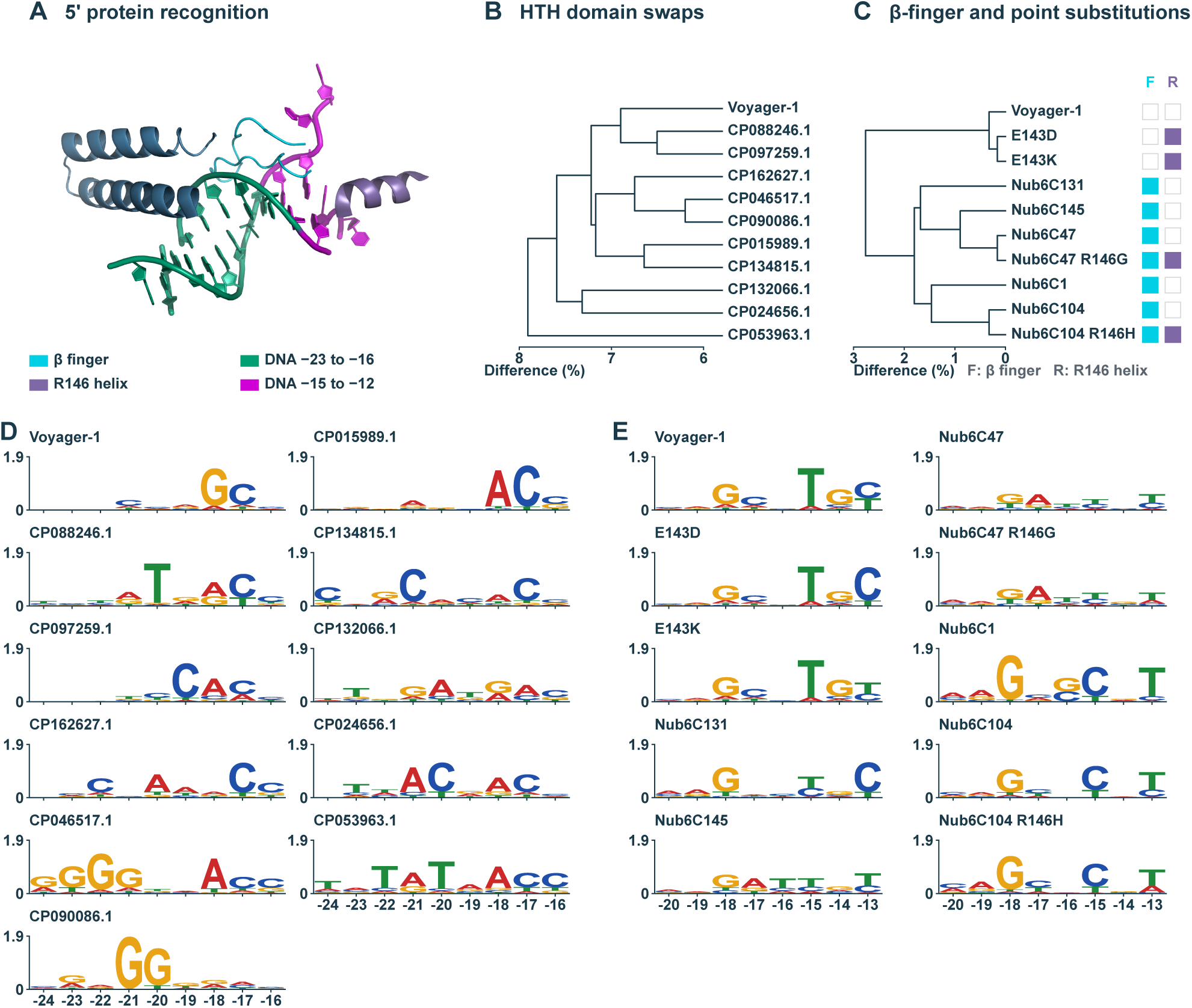
Reprogramming 5′ specificity. (A) The 5′ recognition elements on the target duplex: the β finger and the R146 helix of the 2a insert, contacting DNA at −23 to −16 and −15 to −12 respectively (key on figure). (B and C) Dendrograms of sequence divergence from Voyager­1, as percentage difference, for the HTH­swap series (B) and for the β­finger and point­substitution series (C). Swatches in C mark which of the β finger (F) and the R146 helix (R) was substituted in each construct. (D) Integration­sequencing logos from −24 to −16 for Voyager­1 and ten HTH donors, labelled by accession. Base preference changes substantially between hybrids. (E) Logos from −20 to −13 for β­finger hybrids and for point substitutions at E143 and R146. Selection across this window is altered by both.

Beta-finger replacement (F370–K390) with counterparts from natural class IIA IEPs abolished activity in some cases, consistent with incompatibility with adjacent elements including the hinges; others retained activity with altered selection across −14 to −18 and new selection at −16 (Figure 5E). Recognition at −19 and −20 was unaffected, so this region is largely separable and HTH-programmed. Position −13 remained strongly pyrimidine-biased.

The structures show the −13 sense base engaged by F370, R376 and L387, and the corresponding non-spliced base coordinated by E143, R146 and T147 of the distal 2a insert — so we tested whether the residual constraint at −13 is imposed through the opposite strand. Substitutions at E143 in wild-type Voyager-1, and at R146 and T147 in beta-finger variants, altered selection at −13 with more modest effects at −14 (Figure 5E): E143D increased the preference for C over T; E143K weakened the pyrimidine preference and permitted G; R146H/T147Q in one beta-finger background strongly increased selection for A. Preference at −13 therefore arises from the beta finger and the distal 2a insert acting on opposite sides of the DNA fork, and can be modulated from either.

### 3′ recognition is set by HNH, the upper linker, the thumb and RT0

We expected 3′ reprogramming to be harder, and the register shifts indicate greater conformational freedom of the DNA here. The selected +4 to +8 segment spans both single-stranded and duplex DNA in state 2, contacted by the thumb, upper linker and HNH, while RT0 contacts the sense strand at +8/+9. Replacing HNH from the first CXXC motif (C550) to the C terminus with counterparts from other class IIA IEPs gave 9 of 17 hybrids retaining >95% of parental activity. The most active retained the canonical YAYGY motif with subtle shifts: the first position moved frequently toward A, and some hybrids tolerated substitution at the central pyrimidine or the fourth-position G (Figure S19). HNH exchange therefore largely loosens rather than reprograms specificity.

Exchanging the upper linker with HNH produced substantially larger shifts (Figure 6), consistent with the linker contacting both strands at the re-zipper point. One hybrid yielded two distinct consensus sequences, each of two submotifs either contiguous or separated by one nucleotide. Extending the exchange to thumb segments involved in DNA binding and quaternary assembly gave active proteins with markedly altered 3′ specificity; combining thumb, upper linker and HNH substitutions on the Voyager-1 scaffold in one instance gave a preferred sequence closely matching the known target site of the donor IEP (Figure S14).

**Figure 6.**
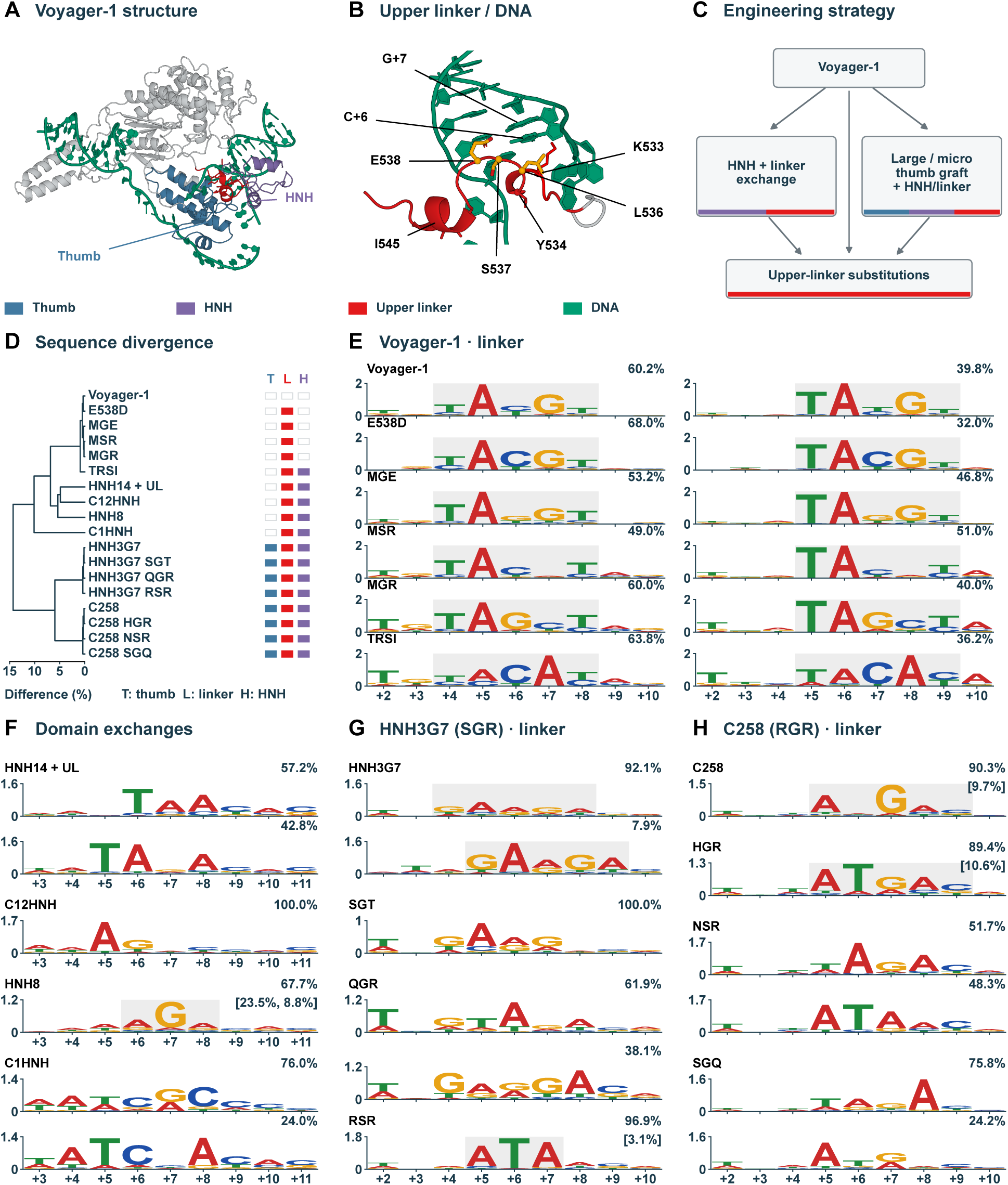
Reprogramming 3′ specificity. (A) Voyager­1 state 2, coloured to show the exchanged segments: thumb, upper linker and HNH. (B) The upper linker threaded between the re­annealing strands, with the pass­through residues K533, Y534, L536, S537, E538 and I545 against C+6 and G+7. (C) Engineering strategy: HNH­plus­linker exchange, large or micro thumb grafts combined with HNH/linker exchange, and point substitution across the pass­through motif, singly and in combination. (D) Dendrogram of sequence divergence of the resulting proteins from Voyager­1, with the matrix at right marking which of thumb (T), linker (L) and HNH (H) was substituted in each. (E) Integration­sequencing logos from +2 to +10 for Voyager­1 and upper­linker substitutions. (F) Logos for domain exchanges. HNH exchange alone loosens rather than reprograms specificity; adding the upper linker produces substantially larger shifts. (G and H) Logos for pass­through triplet substitutions on two reprogrammed backgrounds, HNH3G7 (SGR) and C258 (RGR). Throughout, grey boxes mark the recovered consensus motif and follow its register; where two consensus classes were recovered both are shown with the percentage of reads in each; bracketed percentages denote minor classes. The bits axis differs between rows, so letter heights are not comparable between logos. Logos were generated after grouping sequences by pairwise correlation to accommodate register shifts (STAR Methods).

Point substitution across the upper-linker pass-through motif K533–E538 confirmed its predicted role at +5 to +7. Y534, tolerated other aromatics. K533 substitutions remained active, varying in the strength of selection at +6, +7 and +8, and K533R — common among Voyager-1 relatives — was significantly more active. S537G altered selection at +6 toward G; E538N, E538Q, E538S and E538D all retained good activity and specificity but relaxed notably at +7, which now permitted a purine. Combined substitutions here, in both Voyager-1 and fully reprogrammed backbones, showed substantial capacity for recoding +6/+7 (Figure 6). Finally, substituting the RT0 residues contacting the backbone and minor groove at +8/+9 had a subtle effect: G residues became substantially more frequent, but A remained excluded (Figure S15F).

3′ recognition is therefore driven jointly by upper-linker residues — principally K533, Y534, L536, S537 and E538 — with contributions from the thumb, HNH insert and RT0. This supports a two-stage reprogramming strategy: large exchanges spanning upper linker, HNH and thumb generate substantially recoded clones, while focused substitution can then refine.

#### Retrohoming occurs efficiently in mammalian cells without Mg²⁺ supplementation

Earlier studies reported that group IIA introns were unable to undergo retrohoming in mammalian cells without substantial supplementation of the culture medium with Mg²⁺^19^. We therefore investigated the activity of Voyager-1 in HEK293 cells. Cells were co-transfected by lipofection with plasmids encoding the Voyager-1 IEP, T7 RNA polymerase, a T7-promoter-driven intron and the cognate retrohoming target, so that all components were supplied as DNA and the complex assembled *in situ* (Figure 7A). Active retrohoming complexes were readily reconstituted, as demonstrated by highly efficient integration into co-delivered plasmids. Integration was assessed by gel-based detection following junction PCR and by qPCR, with the expected products confirmed by sequencing (Figure 7B). Estimates based on 3′ integration junctions indicated yields of tens of millions of successful integration events per well of a 24-well plate. qPCR analysis of 5′ junctions indicated a comparable frequency, consistent with a high proportion of full-length integration events.

**Figure 7.**
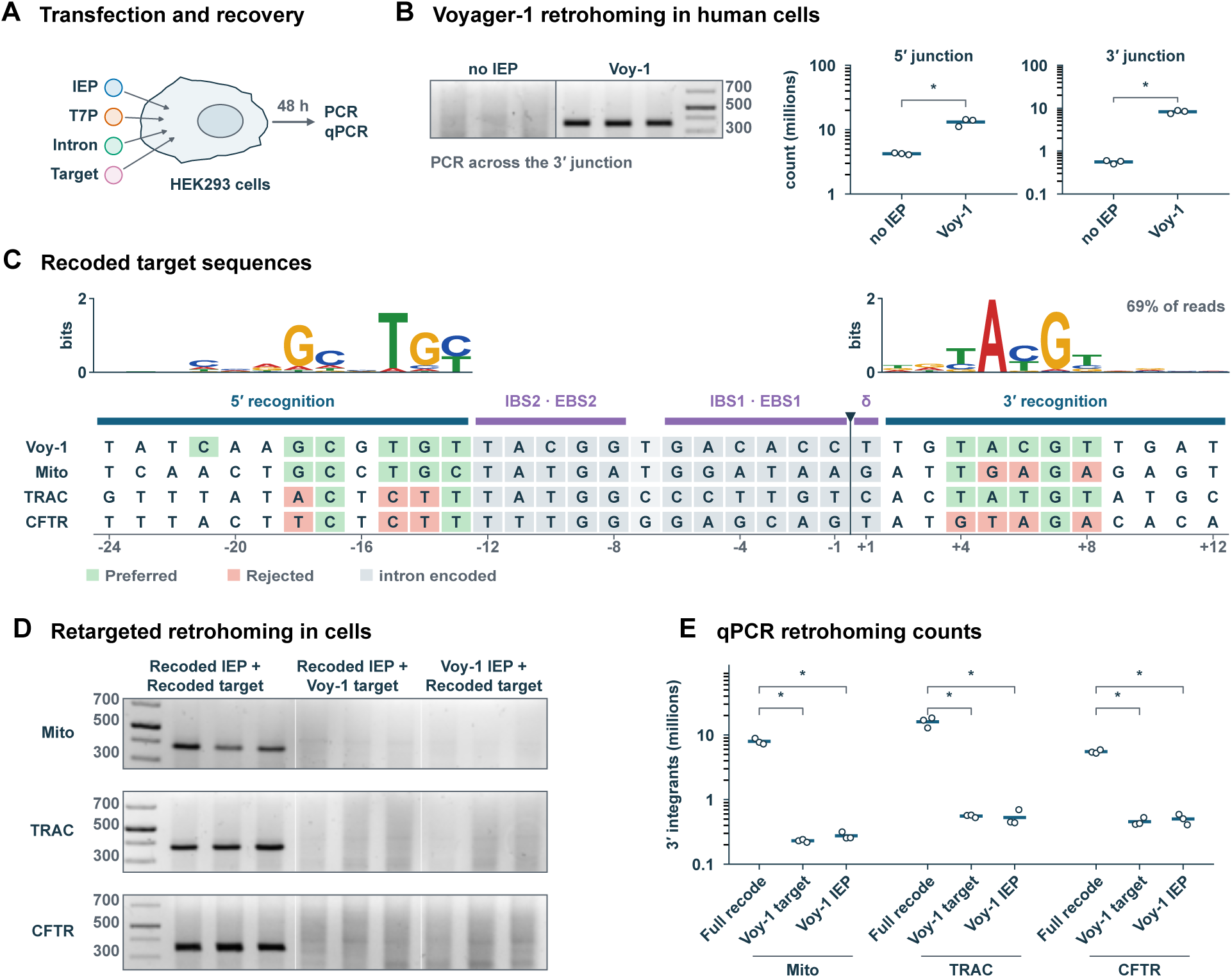
Retrohoming of recoded Voyager­1 to three distinct targets in human cells. (A) HEK293 cells receive the intron­encoded protein (IEP), T7 polymerase, the intron and the target together. After 48 h the DNA is recovered and quantified by PCR and qPCR. (B) Gel shows PCR quantification across the 3′ junction. The graphs show qPCR counts for reactions at the 3′ and 5′ junctions. These show molecules per well from n=3 replicates per condition. Brackets marked by * represent P<0.01 for paired conditions under Welch two­sided t­tests. (C) Voyager­1 base preferences and target sequences. (D) Gel showing PCR products from the 3′ integration junction for recoded target/IEP pairings along with negative controls that replace the recoded target or IEP with the Voyager­1 parent. All conditions contain an intron with recoded EBS1/2 sequences. (E) qPCR quantification of conditions described in D.

As an initial exploration of the potential for combined RNA and protein reprogramming in a cellular context, we tested jointly reprogrammed IEPs and introns directed at sequences from intron 1 of the TRAC locus, from CFTR, and from human mitochondrial DNA, each carried on a co-delivered target plasmid (Figure 7C). Comparable levels of integration at these reprogrammed target sites were observed only when the corresponding reprogrammed IEP was supplied in place of the wild-type Voyager-1 protein (Figures 7D and 7E). These results extend our findings on native protein–intron integration to mammalian cells and demonstrate the feasibility of joint IEP and intron reprogramming in this context, highlighting its potential as a platform for genome engineering and, ultimately, therapeutic applications.

## Discussion

How class IIA intron–IEP complexes bind targets, melt duplex DNA and process it was unknown, and what was known came predominantly from one system, LtrA^29^. Screening without in silico pre-selection identified class IIA IEPs that overexpress and purify simply, with high in vitro activity on linear, relaxed duplex DNA. We show that complex target sites are read through extensive protein–DNA networks on both sides of the splice site, in addition to established RNA–DNA pairings.

On the 5′ side, coordinated major- and minor-groove peptides clamp the duplex and prime local melting at the handoff between DNA:DNA and RNA:DNA pairing; EBS2 nucleotides then capture the exposed sense strand, and a flexible linker peptide stabilises it, driving and holding the target open. On the 3′ side, an unusual trapping mechanism exploits precise positioning of DNA and peptide backbones to constrain HNH activity while enforcing specificity through peptide–DNA contacts at the re-annealing strands. This likely matters functionally: HNH in its catalytically competent configuration likely has little autonomous DNA affinity, limiting uncontrolled incision — valuable in an insertion tool.

Our systems retrohome efficiently into linear, relaxed duplex targets at low complex-to-target ratios and in mammalian cells without Mg²⁺ supplementation. Neither supercoiling nor elevated Mg²⁺ is therefore a general requirement of class IIA retrohoming, and the earlier poor performance of GII introns in eukaryotic cells is unlikely to reflect a single biochemical constraint of this kind. This finding re-invigorates the potential of GII introns for programmable gene insertion in animal cells.

Protein-based reprogramming proved facile, and the very large number of distinct IIA introns and native residence sites in public databases^24,36^ suggests the accessible space is correspondingly large. Because the coding elements are physically separated into discrete protein and RNA segments that recombine productively, targeting complexes can be assembled combinatorially. Combined RNA- and protein-encoded recognition interrogates up to 24 residues 5′ and at least 9 residues 3′ of the splice site — a footprint of up to 35 bp, not all of it read. Protein-driven specificity can readily be engineered to select approximately 4–5 nucleotides on the 5′ side and three or more on the 3′ side; with the 12–13 residues of RNA-encoded recognition^16,20^, this brings typical specificity to the equivalent of around 20 bp, and more in some cases — sufficient to design complexes for single targets in mammalian genomes. Coupling the approaches used here with predictive protein design should make such retargeting increasingly routine.

Beyond insertion by programmable target-primed synthesis, the intermediates themselves may be useful. Sequence-dependent strand separation and nicking generate an accessible single-stranded segment of 21 nucleotides displaced from the helix, which could be exploited in targeted sequencing, DNA assembly, or genome modification strategies that use the exposed flap for hybridisation and ligation. Our current focus is the delivery and assembly of complexes required for programmed insertion in eukaryotic nuclei and mitochondria.

## Acknowledgments

This publication is based on research funded in part by the Gates Foundation. The findings and conclusions contained within are those of the authors and do not necessarily reflect positions or policies of the Gates Foundation.

## Author Contributions

N.D.-A. — Developed the methodology, conceived, planned and conducted experiments, managed and maintained the research data for analysis, performed and developed computational techniques and algorithms, and provided project oversight. Reviewed and edited the original draft.

F.B. — Conceived, planned and conducted experiments, developed the integration sequencing methodology, managed and maintained the research data for analysis, and provided project oversight. Reviewed and edited the original draft.

S.C. — Conceived, planned and conducted experiments, developed splicing detection assays, managed and maintained the research data for analysis and provided project oversight. Reviewed and edited the original draft.

I.C. — Conceived, planned and conducted experiments, managed and maintained the research data for analysis.

N.D. — Conceived, planned and conducted experiments, managed and maintained the research data for analysis.

J.L. — Conceived, planned and conducted experiments, managed and maintained the research data for analysis.

C.L. — Conceived, planned and conducted experiments to produce soluble recombinant proteins.

J.W. — Managed research data for analysis, applied statistical and computational techniques to analyse the data, developed computer algorithms. Reviewed and edited the original draft.

J.E. — Conducted integration-sequencing experiments and subsequent data analysis.

S.T.-P. — Conceived, planned and conducted experiments, managed and maintained the research data for analysis.

M.L.M.V. — Conducted experiments, managed and maintained the research data for analysis.

D.B. — Conducted experiments, managed and maintained the research data for analysis.

S.W.H. — Conducted Cryo-EM data collection and analysis, structural model building and prepared figures.

L.C. — Conducted Cryo-EM sample and grid preparation.

D.Y.C. — Provided project administration of the cryo-EM works.

N.A.A. — Formulated the overarching research goals and aims and provided oversight and leadership for the research activities and planning, prepared the original draft. Secured the financial resources for the project.

## STAR Methods

### 1. EXPERIMENTAL MODEL AND STUDY PARTICIPANT DETAILS

#### Bacterial strains

Intron-encoded proteins (IEPs) were expressed in *Escherichia coli* BL21(DE3).

#### Cell lines

HEK293 cells were cultured in DMEM (high glucose) supplemented with 10% (v/v) heat-inactivated fetal bovine serum, GlutaMAX and 100 U/mL penicillin / 100 µg/mL streptomycin, at 37 °C in a humidified atmosphere of 5% CO₂. Cells were passaged at 80–90% confluency by dissociation with TrypLE and used between passages 1 and 30. Cultures were confirmed free of mycoplasma.

### 2. METHOD DETAILS

#### General plasmid cloning

All components were cloned into a vector (see Table S1, Plasmids used in this study) containing an ampicillin (*bla*) resistance gene via Gibson assembly using NEBuilder HiFi DNA Assembly Master Mix. Fragments for Gibson assembly were either amplified by PCR using Q5 High-Fidelity 2X Master Mix or obtained as synthetic linear DNA fragments (Twist Bioscience). Vectors were linearised by PCR also using Q5 High-Fidelity 2X Master Mix. Assemblies were transformed into NEB 5-alpha competent cells and selected on LB agar containing carbenicillin (100 µg/mL). Plasmid DNA was isolated with EasyPure Plasmid MiniPrep Kit and verified by either Sanger sequencing or whole plasmid nanopore sequencing by GENEWIZ (Azenta Life Sciences).

#### Production of intron RNA

##### In vitro transcription and self-splicing

Intron RNAs flanked by 5′ and 3′ exons were generated by in vitro transcription of PCR amplified templates containing a T7 promoter (primers listed in Table S2). Transcription reactions were carried out in transcription buffer (40 mM Tris-HCl pH 7.5, 30 mM MgCl_2_, 10 mM DTT, 2 mM spermidine) supplemented with 0.15 µg/µL T7-RNA polymerase (T7-P, in-house), 0.014 µg/µL inorganic pyrophosphatase (YIPP, in-house) and 0.05 µg/µL MBP-RNase inhibitor (MBP-RI). The reactions were incubated at 37 °C for 2–4.5 h. Template DNase digestion was carried out for 0.5–1 h at 37 °C by adding CaCl_2_ to a final concentration of 0.75–1.4 mM and 0.05 U/µL TURBO DNase (AM2238 ThermoFisher Scientific). The RNA was then either purified using GeneJet RNA cleanup kit (K0731 ThermoFisher Scientific) or RNAClean XP-beads (A63987 Beckman Coulter), eluting into RNase-free water or RNA Storage Solution (ThermoFisher Scientific, 1 mM sodium citrate pH 6.5). For self-splicing reactions, the purified RNA was diluted 10X in 40 mM Tris-HCl pH 7.5, subjected to a pre-folding cycle (90 °C 1 min followed by a stepwise reduction to 47 °C) and then incubated in self-splicing buffer (500 mM NH_4_Cl, 40 mM Tris-HCl pH 7.5, 50 mM MgCl_2_) for 1–2 hours at 47 °C. Self-spliced RNA was purified using a GeneJet RNA cleanup kit (K0731 ThermoFisher Scientific), eluting into RNase-free water.

#### General pre-qPCR retrohoming protocol

Unless otherwise stated, in vitro retrohoming reactions prior to qPCR analysis were carried out in 40 µL retrohoming buffer (500 mM KOAc, 40 mM Tris-OAc pH 8.2, 5 mM DTT and 5 mM Mg(OAc)_2_) in the presence of MBP-RNase inhibitor and 10 to 25 µM of each dNTP (NEB N0447) for 1 h at 37 °C. Reactions in Figure S16 also included 2 mM spermidine and 3.75% (v/v) DMSO. Retrohoming was performed using 0.67 pmol (4.03 × 10¹¹ molecules) of a 243-bp PCR generated target together with 2.68 pmol Voyager-1 intron RNA and IEP (4:1 ratio RNP:target).

#### Production of intron-encoded proteins (IEPs)

##### Expression

The initial protein expression protocol was identical to that used for cryo-electron microscopy (cryo-EM, vide infra) and structural studies. However, the protocol was progressively adapted to facilitate higher-throughput protein production.

##### Initial modification of the expression protocol

Following overnight incubation on LB agar plates, five colonies were randomly selected and inoculated into 14 mL of LB medium supplemented with 100 µg/mL carbenicillin in 50 mL Falcon tubes covered with breathable sealing membrane. Cultures were grown at 37 °C with shaking at 220 rpm. Upon reaching an OD₆₀₀ of 0.6, the culture volume was reduced to 10 mL, and the cultures were transferred to 20 °C with shaking at 220 rpm for 15 min. Protein expression was induced by the addition of IPTG and ZnCl₂ to final concentrations of 0.5 mM and 0.1 mM, respectively, and cells were incubated overnight. Post-induction cells were harvested by centrifugation at 3,200 × g for 5 min. Pellets were stored frozen at −60 °C until further purification.

##### Final modification: high-throughput expression protocol (autoinduction)

To further streamline the expression procedure, auto-induction cultures were grown based on the method of Studier^1^. Recovered transformants (15 µL) were directly inoculated into 3 mL of LB medium supplemented with 0.5% (v/v) glycerol, 0.05% (w/v) glucose, 0.1 mM ZnCl₂ and 100 µg/mL carbenicillin. The cultures were grown overnight at 37 °C with shaking at 220 rpm.

The following day, 150 µL of each overnight starter culture was inoculated into 15 mL of fresh LB medium supplemented with 0.5% (v/v) glycerol, 0.05% (w/v) glucose, 0.2% (w/v) lactose, 0.1 mM ZnCl₂ and 100 µg/mL carbenicillin. Cultures were incubated at 37 °C with shaking at 220 rpm for 2.5 h before being transferred to 20 °C and incubated overnight with continued shaking at 220 rpm. Cell pellets were subsequently harvested and stored frozen as above.

##### IEP Purification

Cell pellets were thawed at room temperature and resuspended in either 0.5 mL (10 mL non-autoinduction preparations) or 0.75 mL (15 mL autoinduction preparations) of lysis buffer (50 mM Tris-HCl pH 7.5, 1 M NaCl, 30 mM imidazole, 1 mg/mL lysozyme, and Pierce™ EDTA-free protease inhibitor cocktail at the manufacturer’s recommended concentration). Samples were incubated for 30 min at room temperature on an end-over-end rotator to facilitate lysozyme-mediated bacterial lysis. Partially lysozyme-lysed samples were sonicated on ice at 50% amplitude for 15 s (10 mL non-autoinduction preparations) or 30 s (15 mL autoinduction preparations). An aliquot of the total lysate was collected for subsequent analysis of protein expression. The lysate was then clarified by centrifugation at 20,000 × g for 5 min, and an aliquot of the resulting supernatant (soluble fraction) was collected to assess protein solubility.

The soluble fraction was incubated with 100 µL of washed ABT Ni-Agarose EXTRACHEL resin slurry (50 µL bed volume) for 60 min at room temperature with end-over-end rotation to facilitate protein binding. The resin–lysate mixture was subsequently transferred to 2 mL Pierce™ disposable columns. An aliquot of the flow-through fraction may be collected to assess binding efficiency. The resin was then washed with at least 20 bed volumes of wash buffer (50 mM Tris-HCl pH 7.5, 1 M NaCl, and 30 mM imidazole). Bound proteins were eluted using two bed volumes of elution buffer (50 mM Tris-HCl pH 7.5, 1 M NaCl, and 300 mM imidazole), collected as four sequential fractions. A post-elution resin sample may be collected to assess elution efficiency. Each elution fraction (3 µL) was tested for the presence of protein with 100 µL of Pierce™ Bradford Plus Protein Assay Reagent and the most positive fractions were pooled and transferred to a Slide-A-Lyzer MINI dialysis device (10 kDa MWCO). Dialysis was performed overnight at 4 °C with gentle stirring against at least a 300-fold excess volume of dialysis buffer (50% v/v glycerol, 20 mM Tris-HCl, pH 7.5, 400 mM NaCl, and either 2 mM DTT or 1 mM TCEP), followed by an additional 2 h dialysis the next day against fresh dialysis buffer.

Quantification and qualification of purified IEPs were performed as described below for cryo-electron microscopy (cryo-EM) and structural studies

For Voyager-1 with Cytiva™ Protein™ Select Tag, the pellet from a 200 mL culture was resuspended in 20 mL of lysis buffer (50 mM Tris-HCl pH 7.5, 1 M NaCl, 10% glycerol, 1 mg/mL lysozyme, and Pierce™ EDTA-free protease inhibitor cocktail at the manufacturer’s recommended concentration), then sonicated and cleared as above, and bound with 1 mL of the resin (500 µL bed volume). Bound resin was washed with wash buffer, which was identical to lysis buffer minus lysozyme and protease inhibitor cocktail. A 4 h hold step was performed to remove the tag as per the manufacturer’s recommendation.

Purified proteins were stored at −20 °C.

##### Production of MBP-RNase Inhibitor

Expression and purification were performed using the same protocol as that described for cryo-electron microscopy (cryo-EM) and structural studies, with the following modifications. Kanamycin was used at a final concentration of 50 µg/mL. Cells were grown in Terrific Broth (TB) medium to an OD₆₀₀ of 1.0, followed by induction with 0.5 mM IPTG and 5 mM DTT.

For purification, cells from 400 mL of culture were resuspended in 50 mL of lysis buffer containing 50 mM Tris-HCl (pH 7.5), 1 M NaCl, 10% glycerol, 30 mM imidazole, 1 mg/mL lysozyme, and Pierce™ EDTA-free protease inhibitor cocktail at the manufacturer’s recommended concentration. The lysate was applied to 300 µL of resin bed volume and washed with wash buffer containing the same components as the lysis buffer, excluding lysozyme and protease inhibitor cocktail. Bound protein was eluted using elution buffer (50 mM Tris-HCl (pH 7.5), 1 M NaCl, 10% glycerol, 300 mM imidazole). The eluted protein was subsequently dialysed against dialysis buffer containing 50% v/v Glycerol, 20 mM Tris-HCl (pH 7.5), 400 mM NaCl, and 2 mM DTT.

##### Production of YIPP

Expression and purification were performed using the same protocol as that described for cryo-electron microscopy (cryo-EM) and structural studies, with the following modifications. Cells were grown in 3x Luria Broth (3xLB) medium to an OD₆₀₀ of 1.0, followed by induction with 0.5 mM IPTG and 1 mM DTT for overnight before harvesting by centrifugation at 2,800 × g for 10 min (SLA-3000 rotor). Pellets were stored frozen at −60 °C until used

For purification, cells from 400 mL of culture were resuspended in 20 mL of lysis buffer containing 50 mM Tris-HCl (pH 7.5), 1 M NaCl, 10% glycerol, 5 mM imidazole, 1 mg/mL lysozyme, and Pierce™ EDTA-free protease inhibitor cocktail at the manufacturer’s recommended concentration. The lysate was applied to 500 µL of resin bed volume and washed with wash buffer containing the same components as the lysis buffer, excluding lysozyme and protease inhibitor cocktail. Bound protein was eluted using elution buffer (50 mM Tris-HCl (pH 7.5), 1 M NaCl, 10% glycerol, 300 mM imidazole). The eluted protein was subsequently dialysed against dialysis buffer containing 50% v/v Glycerol, 20 mM Tris-HCl (pH 7.5), 400 mM NaCl, and 2 mM DTT.

##### Production of T7 Pol

Expression and purification were performed using the same protocol as that described for cryo-electron microscopy (cryo-EM) and structural studies, with the following modifications. Carbenicillin was used at a final concentration of 100 µg/mL. Cells were grown in Terrific Broth (TB) medium to an OD₆₀₀ of 1.0, followed by induction with 0.5 mM IPTG for 4 h before harvesting by centrifugation at 2,800 × g for 10 min (SLA-3000 rotor). Pellets were stored frozen at −60 °C until used.

For purification, cells from 400 mL of culture were resuspended in 20 mL of lysis buffer containing 50 mM sodium phosphate monobasic anhydrous (pH 8), 0.5 M NaCl, 10mM EDTA, 10 mM imidazole, 1 mg/mL lysozyme, and Pierce™ EDTA-free protease inhibitor cocktail at the manufacturer’s recommended concentration). The lysate was applied to 800 µL of resin bed volume and washed with wash buffer containing the same components as the lysis buffer, excluding lysozyme and protease inhibitor cocktail. Bound protein was eluted using elution buffer (50 mM sodium phosphate monobasic anhydrous (pH 8), 0.5 M NaCl, 10mM EDTA, 300 mM imidazole The eluted protein was subsequently dialysed against dialysis buffer containing 50% v/v Glycerol, 20 mM Tris-HCl (pH 7.5), 0.5 M NaCl, and 2 mM DTT.

##### Reverse transcriptase (RT) activity of IEPs

An isothermal fluorescence intensity (FI) assay was developed as an initial assessment of IEP RT activity (see Figure S3A(ii) for a schematic representation). Reaction mixtures contained 50 mM Tris-HCl pH 8.0, 300 mM KCl, 10 mM MgCl_2_, 2 mM DTT, 0.1 µM FAM-labelled RNA RT_probe oligo, 0.5 µM RT_quencher oligo, 0.5 µM RT_primer oligo, 0.5 mM dNTPs and 100 ng/µL IEP in a final volume of 40 µL (see Table S2 for oligo sequences). All buffer and reaction components except the IEP were pre-assembled and heated at 96 °C for 5 min in a PCR thermocycler before cooling at room temperature for 5 min. IEP addition was made by spinning in from reaction tube lids (microfuge), then reactions were mixed by flicking, spun down, and transferred to a T8-ISO instrument (Axxin) for a minimum of 10 min at 37 °C. Reaction tubes were placed into the T8-ISO device within 5 s of the final spin step. Displacement of the quencher oligo by IEP RT activity is proportional to the detected FAM fluorescence (see Figure S3A(iii)).

##### IEP sequences and nomenclature

Intron-encoded protein (IEP) sequences used for the sequence comparisons in this study were retrieved from the NCBI Protein database by text-based searches (for example “group II intron” and “LtrA”) followed by manual inspection of the returned entries for the reverse-transcriptase, maturase and endonuclease features described in the Results. Sequences retained for comparison were assigned in-house identifiers, and these in-house names are the labels used on the alignment panels in the supplementary figures. Table S3 cross-references every in-house name appearing in the figures to its database protein accession.

Two points of nomenclature should be noted. First, the Eco prefix records the search set in which a sequence was recovered and not the genus of the source organism: Eco1–Eco10 were recovered from a search of Escherichia and related enterobacteria and include Klebsiella and Salmonella entries, while Eco12–Eco14 are Listeria, Eco15–Eco17 Salmonella and Eco20 a Streptococcus entry. Second, the designations BacF and Firmus refer to the same Bacillus firmus protein, which appears under both names across the supplementary alignment panels. Accessions are given at the protein level; where a genome assembly rather than a protein entry is the primary record this is stated in the table.

#### Cryo-EM and structural biology

##### IEP expression

Constructs were transformed into chemically competent BL21(DE3) and plated on LB agar containing 100 µg/mL carbenicillin, then grown overnight at 37 °C. Colonies were scraped from the transformation plate into 400 mL of either 1x or 3x LB medium containing 100 µg/mL carbenicillin. Cultures were grown at 37 °C with shaking at 220 rpm to an OD_600_ of 0.6 (for 1x LB) or 1.0 (for 3x LB), then transferred to 20 °C with shaking at 220 rpm for 15 min. Expression was induced by addition of IPTG, ZnCl₂ and DTT to final concentrations of 0.5 mM, 0.1 mM and 1.0 mM respectively. Cultures were grown overnight at 20 °C and harvested post-overnight by centrifugation at 2,800 × g for 10 min (SLA-3000 rotor). Pellets were stored frozen at −60 °C until used.

##### IEP purification

For each target, the pellet from 100 mL of culture was thawed at room temperature, resuspended in 10 mL of lysis buffer (50 mM Tris-HCl pH 7.5, 1 M NaCl, 30 mM imidazole, 1 mg/mL lysozyme and Pierce™ EDTA-free protease inhibitor cocktail at the manufacturer’s recommended concentration) and incubated for 30 min at room temperature with mixing. Lysates were sonicated on ice using a small probe, with six cycles of 15 s sonication and 45 s recovery (50% amplitude), then clarified by centrifugation at 26,500 × g for 20 min (SS-34 rotor) and the soluble fraction collected. To each soluble fraction, 100 µL of washed ABT Ni-Agarose EXTRACHEL resin slurry (50 µL bed volume) was added and the samples rotated end over end for 60 min at room temperature. The resin was pelleted by centrifugation at 500 × g for 1 min, the supernatant discarded, and the resin washed twice with 10 mL of wash buffer (50 mM Tris-HCl, 1 M NaCl and 30 mM imidazole) with gentle centrifugation between washes. The resin was then transferred to Pierce™ disposable columns and washed with a further 2 x 5 mL of wash buffer through the column.

Bound protein was eluted into a 96 deep-well block in eight sequential fractions of 50 µL elution buffer (50 mM Tris-HCl pH 7.5, 1 M NaCl, and 300 mM imidazole), each equal to one resin bed volume. Each fraction (3 µL) was assayed with 100 µL of Pierce™ Bradford Plus Protein Assay Reagent, and the most positive fractions were pooled.

Pooled fractions were dialysed in a cassette (10 kDa MWCO) against 500 mL of dialysis buffer (20 mM Tris-HCl, 400 mM NaCl, and 2 mM DTT) in a 1 L beaker, overnight at 4 °C with gentle stirring. Dialysed material was harvested, precipitate was removed by centrifugation (17,000 × g for 5 min), and the samples were quantified and qualified (vide supra).

Protein expression and purification were assessed by SDS-PAGE. Samples (0.5 µL each) of the total lysate, soluble fraction, flow-through (unbound material), post-elution resin (residual bound material), and post-dialysis protein sample were analysed on 14% Tris-glycine gels (Invitrogen) using Tris-glycine SDS running buffer. Gels were stained with Quick Coomassie Stain (New Biotech) to visualise total protein.

Purified proteins were aliquoted in 20 µL volumes, snap frozen and stored at −60 °C.

##### DNA target preparation for Cryo-EM

The dsDNA targets used for complex assembly were PCR amplified from synthetic single-stranded 62-nt DNA templates, purchased with RP-HPLC purification (IDT), using the primer pair matched to each target: Cryo-Voy1_F1 and Cryo-Voy1_R1 for Voyager_1_Cryo-EM_target and EBS1:IBS1_mismatch_Cryo-EM_target, and Cryo-Voy2_F1 and Cryo-Voy2_R1 for Voyager_2_Cryo-EM_target (Table S2).

Amplification of each target was performed as twelve reactions of 50 µL, each containing Q5 High-Fidelity 2X Master Mix (NEB), 160 pmol of each primer and 0.1 pmol of the ssDNA template, using the following program: 16 cycles of (98 °C 15 s, 64 °C 15 s, 72 °C 15 s). Products were resolved on a 2% agarose gel to separate the amplicon from residual primers; bands were excised and the DNA recovered by electro-elution and precipitation, resuspended in 20–25 µL of water and pooled. The pooled DNA was desalted on a Cytiva desalting column, eluted in 1.5 mL of water, frozen at −80 °C and dried overnight under vacuum. The pellet was resuspended in water to approximately 500 ng/µL, quantified by NanoDrop, and its size and purity confirmed on a 4–12% TBE gel.

##### Complex assembly for Cryo-EM

Voyager-1 preparation for Cryo-EM: 85 pmol of in vitro transcribed intron and 85 pmol of IEP in dialysis buffer (20 mM Tris-HCl pH 7.5, 400 mM NaCl, 2 mM DTT) were incubated in a total volume of 20 µL assembly buffer (300 mM KOAc, 40 mM Tris-OAc pH 8.2, 5 mM Mg(OAc)_2_) in the presence of MBP-RNase inhibitor, for either 20 min (Voyager-1 State 1), 35 min (Voyager-1 State 2) or 45 min (EBS1:IBS1 mismatch) at 30 °C. The reactions were then placed on ice, 85 pmol of DNA target and extra RNase inhibitor were added. The buffer was adjusted to obtain a 40 µL final volume in retrohoming buffer (150 mM KOAc, 40 mM Tris-OAc pH 8.2, 5 mM Mg(OAc)_2_, 5 mM DTT) and incubated either for 40 min (Voyager-1 State 1), 7 min (Voyager-1 State 2) or 10 min (EBS1:IBS1 mismatch) at 37 °C and then stored in ice before loading on Cryo-EM grids.

Voyager-2 preparation for Cryo-EM: 111 pmol of in vitro transcribed and self-spliced intron, 111 pmol of IEP in dialysis buffer (1.79 µL of 20 mM Tris-HCl pH 7.5, 400 mM NaCl, 2 mM DTT) and 111 pmol of DNA target were assembled together in a 40 µL final volume of retrohoming buffer (150 mM KOAc, 40 mM Tris-OAc pH 8.2, 5 mM Mg(OAc)_2_, 5 mM DTT) in the presence of MBP-RNase inhibitor, and incubated for 35 min at 37 °C (Voyager-2 State 1). The reaction was then stored in ice before loading on Cryo-EM grids.

##### Cryo-EM Grid Vitrification

A 3 µL aliquot of Voyager-1 sample was applied onto an UltraFoil R1.2/1.3 Au 300-mesh grid. Prior to sample application grids were glow-discharged on both sides at 25 mA for 60 s per side. Grids were vitrified using a Vitrobot (ThermoFisher Scientific) operated at 4 °C and 95% relative humidity, with a blotting time of 3 s and a blot force of −5.

##### Cryo-EM Data Collection

Cryo-EM data were acquired on a Titan Krios transmission electron microscope (ThermoFisher Scientific) operated at 300 kV and equipped with a Selectris X energy filter and a Falcon 4i direct electron detector operated in electron counting mode. Images were recorded at a nominal magnification of 165000x, corresponding to a calibrated pixel size of 0.729 Å pixel^−1^. A 10 eV energy-filter slit width was used. A total of 38246 movies were acquired using aberration-free image shift (AFIS) in EPU (ThermoFisher Scientific). Data were collected over a nominal defocus range of −1.8 to −0.6 µm in 0.2 µm increments.

##### Cryo-EM image processing

Movies were pre-processed in real time using WARP^2^. Gain correction, motion correction and contrast transfer function (CTF) estimation were performed prior to automated particle picking using the BoxNet2 masked model within WARP. A total of 2659506 particles were picked, extracted in 512-pixel boxes, and imported into cryoSPARC (v5.0.2)^3^ for subsequent steps of data processing as summarised in Figure S1. In brief, 2D classification was used to distinguish between particles representing Voyager-1 complexes and erroneous picks. Several ab-initio volumes were generated representing Voyager-1 assemblies and “junk” particles. Through several iterative rounds of heterogenous refinement and ab-initio volume generation junk particles and particles from over-represented orientations were discarded, leaving particles representing three major categories of Voyager-1 assemblies (States 1–3). These volumes were further classified using focussed 3D classification on the DNA and/or HNH regions of the volumes. Final volumes were subjected to non-uniform refinement, producing final maps for States 1–3 at resolutions of 3.02, 3.04 and 5.11 Å respectively.

The PDB file 8H2H was docked into the map for state 1 and used as a guide to manually re-model the entire protein, RNA and DNA chains in Coot 0.9.8.95^4^, and subsequently refined in Phenix 1.21^5^. The state 1 model was then docked into maps for states 2 and 3 and subsequently manually adjusted and refined using Coot and Phenix.

#### qPCR

TaqMan qPCR assays were carried out with 1 µL of 1:100 dilution for in vitro retrohoming reactions or 2 µL of DNA extract for in vivo retrohoming quantification, in a total of 20 µL. Luna universal probe qPCR master Mix (NEB M3004) was used, following manufacturer instructions. To detect the integration junction, a primer specific for the target and a primer specific for the intron sequence were used to amplify either the 5′ or the 3′ region of the integration site. The Taqman probe binds to a sequence in the target. The reactions were assembled in triplicates in an AriaMx non-skirted 96 well optical plate (Agilent Cat 401494), sealed with AriaMx Adhesive seal (Agilent Cat. 401492) and run on an AriaDx qPCR machine (Agilent) with the following program: 95 °C 60 s, 40 cycles (95 °C 15 s; 60 °C 30 s). The data were analysed with AriaDx 2.0 Software. When used, the standard curve included 4 to 6 points of 10X serial dilutions ranging from 200 pg to 0.002 pg. The DNA fragments used for the standard curve were purchased from Twist Bioscience. Primers, probes and Standard DNA used are listed in Table S2.

#### Retrohoming in mammalian cells

##### Plasmids

Group II intron retrohoming was reconstituted in HEK293 cells from four co-delivered plasmids: (i) a T7 RNA polymerase expression plasmid, codon-optimised in house for mammalian expression; (ii) an intron donor plasmid carrying the Voyager-1 GII intron under T7 transcriptional control; (iii) a target plasmid carrying the cognate intron target sequence; and (iv) an expression plasmid encoding the Voyager-1 IEP, codon-optimised in house for mammalian expression. For each set, the negative control was the identical transfection with the IEP plasmid omitted.

The native Voyager-1 system was compared side by side with three recoded sets, in each of which the intron, its target and the IEP were matched. Each set is named for the locus from which its target sequence is taken: the mitochondrial set (target and intron Mito_L11598r, IEP Mito_Voyager-1_8141_6270), the TRAC set (TRAC_I1_3049, TRAC_Voyager-1_9031-5943) and the CFTR set (CFTR_I1_1857r, CFTR_Voyager-1_9034-6270). All plasmids used in this study are listed in Table S1.

##### Transfection

HEK293 cells were seeded at 2.4 × 10⁵ cells per well in 500 µL of complete medium in 24-well plates and transfected the following day with Lipofectamine 3000 (Thermo Fisher Scientific). Each condition was transfected in triplicate.

Two mixes were prepared per condition: MM1, 25 µL Opti-MEM with 1 µL Lipofectamine 3000 reagent per well; MM2, 25 µL Opti-MEM with 1 µL P3000 reagent per well plus the plasmid DNA. All plasmids were diluted to 100 ng/µL and 1 µL of each added per reaction, except the intron donor plasmid which was added at 10 ng/transfection for Voyager-1 or 20 ng/transfection for the recoded IEPs. MM1 was added to MM2, incubated for 10 min at room temperature, and 50 µL of the combined mix added dropwise per well. The medium was replaced with fresh complete medium 4 h after transfection. After 24 h the cells were trypsinised from the well and replated in a fresh 24-well plate and harvested a further 24 h later, totalling 48 h post-transfection. In the no-IEP control the IEP plasmid was omitted. Recoded sets were also tested against Voyager-1 IEP or the intron donor plasmid carrying the Voyager-1 GII intron as controls.

##### Plasmid DNA recovery

At 48 h post-transfection, cells were washed with PBS, collected, and pelleted by centrifugation at 1,000 × g for 3 min. Cell pellets were resuspended in 50 µL PBS, 50 µL of QuickExtract DNA Extraction Solution (LGC) was added and lysis was carried out following manufacturer’s protocol.

##### PCR detection of retrohoming

Integration of the intron into the target plasmid was detected by endpoint PCR with DreamTaq Hot Start Green PCR Master Mix (2X; Thermo Fisher Scientific) under the manufacturer’s standard conditions. Each 50 µL reaction contained 2 µL of extracted DNA as template and 0.5 µM of each primer. The forward primer Voy1_3-for-7 anneals within the intron sequence and the reverse primer Voy1_target_R2 within the target plasmid (Table S1), so that a 378-bp product is generated only when the intron has integrated into the target plasmid.

Cycling conditions were initial denaturation at 95 °C for 2 min; 31 cycles of 95 °C for 30 s, 62 °C for 30 s and 72 °C for 30 s; final extension at 72 °C for 10 min.

Products were resolved on a 2% (w/v) agarose gel in 1× TBE buffer alongside 5 µL of GeneRuler 1 kb Plus DNA Ladder (Thermo Fisher Scientific), stained with SafeView, and imaged on a Bio-Rad ChemiDoc. The same primer pair, amplicon size and cycling conditions were used for the native Voyager-1 system and for all three recoded sets.

#### dsDNA fluorophore-quencher retrohoming probes

dsDNA fluorophore-quencher retrohoming probes were prepared by either:

i. Annealing two 60-nt synthetic oligonucleotides in reaction buffer, in a 1:1 ratio of F:Q;
ii. Annealing two 60-nt synthetic oligonucleotides in reaction buffer, in a 1:1.5 ratio of F:Q;
iii. PCR to generate a 243-bp amplicon from a plasmid, using a 5′-FAM-labelled forward primer and unlabelled reverse primer, followed by labelling of 3′-ends using a terminal deoxynucleotidyl transferase enzyme and a quencher-labelled dideoxy nucleotide – note that both ends of the dsDNA probe will be labelled with quencher, as opposed to a single 3′-end as obtained by annealing two synthetic oligonucleotides.

Methods (i)–(iii) are detailed below:

i. Dried single stranded oligonucleotides were purchased (IDT) with RP-HPLC purification, then reconstituted in water; see Table S2 for the nucleotide sequences of 60-nt oligos with a 5′-FAM (Voy1_retro_probe) and 3′-quencher (Voy1_retro_quench). An equimolar mix of Voy1_retro_probe (10 µL at 20 µM) and Voy1_retro_quench (10 µL at 20 µM) in water (5 µL) plus 2X retrohoming buffer (1 M KOAc, 80 mM Tris-OAc pH 8.5, 10 mM Mg(OAc)_2_) was pre-annealed in a PCR thermocycler (MJ Research PTC-225, default temperature ramp rate) using the following program: 95 ^°^C 3 min; 70 °C 5 min; 55 °C 5 min; 30 °C hold. The annealed dsDNA fluorophore-quencher retrohoming probe was used in subsequent assays (4 pmol, i.e. 1 µL of the 4 µM dsDNA probe thus generated, in a 40 µL reaction).
ii. The annealed dsDNA fluorophore-quencher retrohoming probe was prepared as for method (i), but using a mix of Voy1_retro_probe (10 µL at 20 µM) and Voy1_retro_quench (15 µL at 20 µM) plus 2X retrohoming buffer (1 M KOAc, 80 mM Tris-OAc pH 8.5, 10 mM Mg(OAc)_2_) for pre-annealing.
iii. A dsDNA retrohoming target, Voyager_1_target, was PCR amplified from pVoyager_1_target using primers Voy1_probe_F (i.e. a 5′-FAM-labelled forward primer) and Voy1_target_R (Tables S1 and S2). Amplification was performed as 25 reactions of 60 µL, each containing Q5 High-Fidelity 2X Master Mix (NEB), 50 pmol of each of the 5′-FAM-labelled forward and unlabelled reverse primers, and 2.5 ng of the plasmid template, using the following program: 40 cycles of (98 °C 15 s, 72 °C 30 s), then 75 °C 5 min, then 4 °C hold. Reactions were combined into eighteen equal aliquots and each was purified separately on EasyPure PCR Purification Kit columns (TransGen Biotech), eluting each column into 30 µL of warmed water (from an aliquot in a heat block set to 55 °C), to obtain a combined 0.5 mL of 5′-FAM-labelled dsDNA at 0.75 µM.

The 3′-ends of the 5′-FAM-labelled amplicon were then labelled with ATTO-540Q-ddATP using terminal deoxynucleotidyl transferase (TdT, NEB). Eighteen labelling reactions of 50 µL each containing the 5′-FAM PCR product (10 µL at 0.75 µM), 10X TdT reaction buffer (5 µL, NEB), ATTO-540Q-ddATP aq. (2 µL at 50 µM), CoCl_2_ (5 µL at 2.5 mM, NEB), and TdT enzyme stock (4 µL at 20 U/µL, NEB) were incubated in a PCR thermocycler (MJ Research PTC-225) using the following program: 37 °C 90 min; 70 °C 10 min; 10 °C hold. Reactions were combined into nine aliquots of 100 µL and purified on EasyPure PCR Purification Kit columns (TransGen Biotech). The columns were eluted in series, in three sets of three: the first column was eluted with 2 x 35 µL of warmed water (from an aliquot in a heat block set to 55 °C), the eluate was then applied as 2 x approximately 35 µL aliquots to elute the second column, and the eluate from the second column was applied as 2 x approximately 35 µL aliquots to elute the third column; the final eluates were combined, to obtain a combined 0.2 mL of 5′-FAM/3′-quencher dsDNA at 0.55 µM.

#### Retrohoming for isothermal FI and gel assays

For isothermal FI monitoring of retrohoming (see Figure S3C(vii)), a ‘reaction master mix’ was prepared fresh in excess, comprising 2X retrohoming buffer (200 µL of 1 M KOAc, 80 mM Tris-OAc pH 8.5, 10 mM Mg(OAc)_2_), RNase inhibitor (5 µL at 27.5 µM, in dialysis buffer comprising 20 mM Tris-HCl pH 7.5, 400 mM NaCl, 50% Glycerol, 2 mM DTT), fresh aqueous 0.2 M DTT (5 µL), DMSO (15 µL) and 80 mM spermidine (10 µL). To 4 pmol of a 243-bp dsDNA fluorophore-quencher retrohoming probe ‘Voyager_1_target’ plus water (to give a final retrohoming reaction volume of 40 µL) was added ‘reaction master mix’ (23.5 µL) followed by 16 pmol of Voyager_1_precursor_RNA or Voyager_1_DVdel_RNA (in water or Invitrogen RNA Storage Solution, 1 mM sodium citrate pH 6.5), followed by 16 pmol of Voyager_1_IEP (2 µL at 8 µM in dialysis buffer 20 mM Tris-HCl pH 7.5, 400 mM NaCl, 50% Glycerol, 1 mM TCEP), with pipette mixing for each addition. Where dNTPs were included, 1 µL of dNTPs mix (1 mM in each dNTP) was added prior to the reaction master mix, replacing an equal volume of water. The reaction was then monitored for 60 min in a T8-ISO instrument at 37 °C.

To assess the effect of Mg²⁺ concentration on retrohoming (Figure S17), isothermal FI monitoring was performed as above, except that the 2X retrohoming buffer contained 2, 3 or 4 mM Mg(OAc)_2_ in place of 10 mM. This gave final Mg²⁺ concentrations of 1, 1.5 and 2 mM, respectively. Also, the ‘Voyager_1_target’ probe for this experiment was generated using oligos ‘Voy1_retro_probe_-7T’ and ‘Voy1_retro_quench_-7T’.

For gel analysis (see Figure S3C(viii)), reaction mixtures were each brought to 50 µL with water then purified into 30 µL of warmed water (from an aliquot in a heat block set to 55 °C) using EasyPure PCR Purification Kit columns (TransGen Biotech). 4 µL of the purified reaction mixtures, or of unreacted 243-bp dsDNA fluorophore-quencher retrohoming probe ‘Voyager_1_target’ at 0.14 µM, were mixed with 8 µL of glyoxal solution (prepared from 45 µL fresh glyoxal + 125 µL DMSO + 25 µL 0.1 M aqueous sodium phosphate pH 7) and 2 µL of 6X MassRuler DNA Loading Dye (Thermo Fisher Scientific), then heated to 50 °C for 30 min before loading onto a pre-run Novex TBE-Urea 6% PAGE gel (Thermo Fisher Scientific) and the gel run at 180 V 40 min before imaging (ChemiDoc, Bio-Rad).

For the nucleotide walking experiment (see Figure S3D), retrohoming reactions were performed in a similar way (*vide supra*), except that the 243-bp dsDNA retrohoming probe ‘Voyager_1_target’ possessed a sense-strand 5′-FAM label but no quencher, i.e. the amplicon generated by method (iii) above, taken prior to the TdT labelling step. A ‘reaction master mix’ was again prepared fresh in excess, comprising 2X retrohoming buffer (190 µL of 1 M KOAc, 80 mM Tris-OAc pH 8.5, 10 mM Mg(OAc)_2_), RNase inhibitor (5 µL at 27.5 µM, in dialysis buffer comprising 20 mM Tris-HCl pH 7.5, 400 mM NaCl, 50% Glycerol, 2 mM DTT), fresh aqueous 0.2 M DTT (5 µL), DMSO (15 µL) and 80 mM spermidine (10 µL). To 4 pmol of a 60-nt F/Q probe prepared by method (ii) to serve as a fluorescent positive control, or 243-bp dsDNA fluorophore-only retrohoming probe ‘Voyager_1_target’, plus water (to give a final retrohoming reaction volume of 40 µL in all cases) was added ‘reaction master mix’ (22.5 µL). For reactions containing the 243-bp dsDNA target, an additional 1 µL of 2X retrohoming buffer was also added (as the annealed 60-nt probe already carried buffer). 16 pmol of Voyager_1_precursor_RNA or Voyager_1_DVdel_RNA (in water or Invitrogen RNA Storage Solution, 1 mM sodium citrate pH 6.5) was then added, followed by 16 pmol of Voyager_1_IEP (2 µL at 8 µM in dialysis buffer 20 mM Tris-HCl pH 7.5, 400 mM NaCl, 50% Glycerol, 1 mM TCEP), with pipette mixing for each addition. Where dNTPs were included, 1 µL of a dNTPs mix (1 mM in each of the dNTPs stated in Figure S3D, derived from the NEB Deoxynucleotide (dNTP) Solution Set) was added prior to the reaction master mix, replacing an equal volume of water. The reactions then proceeded for 60 min in a T8-ISO instrument at 37 °C with FI monitoring to confirm the positive control (data not shown), then snap-frozen and stored at −60 °C overnight. Reaction mixtures containing the 243-bp probe were then processed for gel analysis as described above, except that 2 µL of the purified reaction mixture was mixed with 4 µL of glyoxal solution and 1 µL of 6X MassRuler DNA Loading Dye.

#### IEP Reprogramming

##### dsDNA retrohoming target libraries

Library oligonucleotides, Library_oligo_, 100 nt in length and possessing the RNP target recognition sequence but with a contiguous section of randomized bases, were purchased from Integrated DNA Technologies (IDT; Table S2). To minimize library bias, it was specified that randomized positions were introduced during the solid-phase synthesis of the oligonucleotides using hand-mixed phosphoramidites, according to a proprietary and empirically derived ratio employed by IDT. Oligonucleotides were purified by desalting only, to further minimize any library bias that may have arisen during chromatographic purification methods.

dsDNA libraries were prepared by PCR amplification of library oligonucleotides, using the following program, which employs a low cycle number to minimize the formation of bubble products (i.e. dsDNA with sections of non-complementary sequence in the library segment): 98 °C 1 min; 5 cycles of (98 °C 20 s, 60 °C 20 s, 72 °C 20 s), then a final cycle of (98 °C 20 s, 68 °C 20 s, 72 °C 320 s), then 4 °C hold^6^. PCR primers (selected from Library_F1, _F2, _R1 and _R2 depending upon the Library_oligo_ sequence) were employed with or without 5′-phosphorylation (_5P), depending on the experiment. 5′-phosphorylation was preferred as it enabled subsequent nanopore sequencing to be performed without use of the ‘End-prep’ step during the sequencing protocol (Oxford Nanopore, Ligation sequencing amplicons - Native Barcoding Kit V14 (SQK-NBD114.24)). The End-prep step, which employs a proprietary mix of enzymes in the NEBNext Ultra II End Repair/dA-Tailing Module (NEB), adds an additional step to the workflow and creates some ‘noise’ (i.e. sequences that are not solely derived from the retrohoming reaction), including additional sense strand DNA that can be detected and read during nanopore sequencing (possibly by extending partial reaction products that can serve as primers). If primers are not 5′-phosphorylated, then an ‘End-prep’ (and no separate dA-tailing of the PCR-generated dsDNA retrohoming target, *vide supra*) is required.

Library oligonucleotide stocks were quantified by absorbance at 260 nm – using extinction coefficients calculated by the manufacturer (IDT) from the sequence by the nearest neighbour method – then diluted in two steps to a 10 µM intermediate stock and a 125 nM working solution for use as PCR template. Amplification reactions contained 1x Q5 High-Fidelity 2X Master Mix (NEB) and were run in one of three formats: 0.5 µM of each primer with 2.5 nM of the library oligonucleotide template in 50 µL, 1 µM with 5 nM in 25 µL, or 1 µM with 5 nM in 50 µL. Since the cycle number is deliberately low, sufficient material was obtained by running identical reactions in parallel, and these were pooled before purification.

Pooled reactions were purified on EasyPure PCR Purification Kit columns (TransGen Biotech), applying 200 to 400 µL of pooled reaction to each column in two or four successive loadings, washing once with the kit wash buffer and eluting in 30 to 37.5 µL of water. A second elution was performed to improve recovery, either by applying a single volume of water (40 to 75 µL) successively to all of the columns, or by passing 35 µL of water through two or four columns in series and repeating this three times; all eluates were then combined.

Where 5′-phosphorylated primers had been used, a 3′ dA overhang was added to the purified product so that it could be ligated directly to a native barcode during the sequencing workflow. Reactions of 50 µL contained 42 µL of purified PCR product, 5 µL of NEBNext dA-Tailing Reaction Buffer and 3 µL of Klenow Fragment (3′→5′ exo−) from the NEBNext dA-Tailing Module (NEB), and were incubated at 37 °C for 30 min before being purified on EasyPure columns as above, normally one reaction per column with a 35 µL elution.

Purified libraries were quantified by absorbance at 260 nm on a spectrophotometer (DeNovix or Implen) using the mass extinction coefficient for a 100 bp dsDNA fragment (101 bp after dA addition), and were stored at −20 °C. Working concentrations were typically between 0.3 and 0.7 µM. Each new synthesis lot was checked for bias by barcoding and sequencing a sample of the library alongside retrohoming reactions, as described below.

##### Determining IEP sequence specificity

The sequence specificity of an IEP was determined by reacting it, as a ribonucleoprotein (RNP) with its intron RNA, against a dsDNA retrohoming target library in which the positions of interest were randomized, and then sequencing the products of the reaction, so that the randomized segment recovered for a given IEP could be compared with that of the unreacted library. Retrohoming reactions of 40 µL contained 16 pmol of intron RNA, 16 pmol of IEP and 4 pmol of dsDNA target library (0.4, 0.4 and 0.1 µM respectively) in a buffer containing 500 mM KOAc, 40 mM Tris-OAc pH 8.5, 5 mM Mg(OAc)_2_, 2.5 mM DTT and 25 µM of each dNTP, together with RNase inhibitor (an in-house MBP-RNase inhibitor fusion protein, normally 0.5 µL of a 2.54 mg/mL (27.5 µM) solution per reaction). Components were added in a fixed order, the target library and water first, then the dNTPs, then the intron RNA and finally the IEP, with pipette mixing after each addition.

In some experiments the reactions additionally contained 3.75% (v/v) DMSO and 2 mM spermidine to boost yield. The intron RNA was either the full-length transcript Voyager_1_precursor_RNA, or a variant Voyager_1_precursor_trimmed_RNA for which the region that interacts with the IEP N-terminus had been trimmed, again depending on the experiment, and the 16 pmol input was calculated from the length of the transcript in use (1.2 kb and 1.0 kb respectively).

Reactions were incubated at 37 °C for 60 min, either in a T8-ISO instrument or in a PCR thermocycler, then treated with 1 µL of RNase A (NEB) at 37 °C for a further 60 min to remove the intron RNA, giving a final volume of 41 µL, and stored at −60 °C until clean-up. Every batch of reactions included a control in which the dsDNA target library was replaced by 4 pmol of 60-nt dsDNA fluorophore-quencher retrohoming probe (Table S2, 1:1 F:Q prepared by method (i) *vide supra*); this control was run in the T8-ISO instrument using IEP Voyager_1_IEP_seqref and its endpoint fluorescence used to confirm that the RNP preparation was active in the reaction buffer lot for that experiment, before the batch was taken forward to sequencing. Every batch also included Voyager_1_IEP_seqref as an internal reference, retrohomed against the same target library and at the same 4 pmol input.

Reactions were purified using RNAClean XP beads (Beckman Coulter; AMPure XP beads in RNase-free buffer) at 0.9 volumes (36.9 µL), washed three times with 500 µL of 70% ethanol and eluted in 15 to 16.3 µL of water. Eluates were quantified using the Qubit dsDNA HS assay kit on a Qubit 2.0 Fluorometer device, 1 µL of eluate being diluted into 200 µL of Qubit buffer.

Where the target library had been amplified with unphosphorylated primers, 200 fmol of each eluate was taken into an End-prep reaction in a final volume of 12.5 µL, using the NEBNext Ultra II End Repair/dA-Tailing Module (NEB) as described in the ONT protocol, and the end-prepped material was quantified again; eluates containing less than 200 fmol were taken forward as 12.5 µL of undiluted eluate. Where the library had been amplified with 5′-phosphorylated primers and dA-tailed, the End-prep step was omitted and the bead eluate was taken directly into barcoding.

Each sample was ligated to a single native barcode in a reaction taking 7.5 µL of sample, using the Native Barcoding Kit 24 V14 (SQK-NBD114.24) or the Native Barcoding Kit 96 V14 (SQK-NBD114.96). In some experiments this volume was treated as a maximum rather than a fixed input: the least concentrated sample in a batch was taken at the full 7.5 µL and every other sample was reduced to the volume containing the same mass of DNA and made up to 7.5 µL with water, so that the barcodes would be represented in comparable numbers in the sequencing data. Otherwise, all samples were taken at 7.5 µL without adjustment, so that read numbers also reflect the amount of material recovered from each reaction.

A sample of the dsDNA target library itself, diluted so that it would not dominate the pool (typically 0.5 µL of a one in ten dilution, made up to 7.5 µL with water), was barcoded and sequenced alongside the retrohoming reactions in every run, and provided the unreacted sequence distribution against which the recovered sequences were normalized.

Barcoded samples were pooled, cleaned up and ligated to the native adapter as described in the ONT protocol for the kit in use, and the adapter-ligated pool was quantified with the Qubit dsDNA HS assay kit. Between 1 and 12 µL of the pool was loaded onto a MinION/GridION Flow Cell (DNA) (FLO-MIN114) on a MinION Mk1B or Mk1D device. Sequencing was performed with real time super-accurate base calling in the MinKNOW software. Flow cells were washed and re-used with unique barcodes where the available pore count allowed.

#### Targeted sequencing of a retrohoming target

A dsDNA retrohoming target, Voyager_1_target, was PCR amplified from pVoyager_1_target using primers Voy1_target_F and Voy1_target_R (Tables S1 and S2). In the absence of dNTPs, Voyager_1_RNP is expected to hydrolyze the P-O bond on the corresponding antisense strand, located between nucleotides +8 and +9 from the splice site (immediately upstream of a dT residue indicated in bold underline in Table S2), thus generating a 5′-phosphorylated strand. Cryo-EM data suggest that this strand is single stranded for up to 21 nucleotides from its 5′-end, thus presenting a sequence for ligation to a modified adaptor for targeted next generation sequencing applications. To this end, ‘native barcode’ (NB) sequence adaptors were designed from sequences published by Oxford Nanopore Technologies, Ltd. (ONT) for their ‘Native Barcoding’ kits in e.g. Chemistry Technical Document CHTD_500_v1_revAV_24July2026, and purchased from IDT, Inc., with ‘standard desalting’ purification; a dA nucleotide was added to the reverse primers to account for the elimination of the end-prep step from the standard ONT kit protocols (indicated in bold in Table S2), and sequences were added to allow one, six or twelve nucleotides of overlap with the putative antisense ssDNA sequence (underlined in Table S2). These oligos were reconstituted in water and combined to a 1 µM concentration of the dsDNA barcodes (e.g. NB01 was prepared by mixing NB1_F and NB1_ReverseT).

To test the targeted sequencing concept, 400 nM of Voyager_1_RNP was reacted with 100 nM of the 243-bp dsDNA Voyager_1_target in the absence of dNTPs, in a buffer containing 500 mM KOAc, 40 mM Tris-OAc pH 8.5, 5 mM Mg(OAc)_2_, 2.5 mM DTT, and 0.34 µM RNase inhibitor. The reaction proceeded for 20 min at 37 °C in a PCR thermocycler, before it was cooled to 10 °C. Two 40 µL reactions were performed in this way and combined, before 3 x 22.5 µL aliquots were combined separately with 7.5 µL of one of the three modified barcodes NB01-NB03 at 1 µM described above, followed by 30 µL of NEB Blunt/TA Ligase Master Mix. Barcode ligation proceeded for 20 min at room temperature, before the three barcoding reactions were each quenched with 12 µL of 0.25 M EDTA aq. Barcoded sample clean-up, native adapter ligation and sample sequencing on an ONT MinION/GridION Flow Cell (DNA) were then performed as per an ONT protocol for the Native Barcoding Kit 24 V14, NBA_9168_v114_revO_15Sep2022, using 12 µL of the final adapter ligated sample at a concentration of 13.32 ng/µL as measured using the Qubit dsDNA HS assay kit on a Qubit 2.0 Fluorometer device.

DNA sequencing was performed for 69 min on a flow cell with 1,288 pores, with real time Super accurate base calling in the MinKNOW software. Interrogation of the FASTQ files thus generated showed that this method allows targeted sequencing of a dsDNA target, and that longer overlaps between the antisense ssDNA strand and barcode reverse strand gave increased numbers of reads, which commenced at the expected position on the antisense strand after stripping of barcode and adapter sequences (see Figure S7 for an illustration of the data).

### 3. QUANTIFICATION AND STATISTICAL ANALYSIS

#### Mammalian cell culture

##### Plasmid retrohoming

Each condition was transfected in triplicate. Retrohoming was scored qualitatively as presence or absence of the 378-bp product on the agarose gel.

##### qPCR data analysis

Quantities were read against a standard curve of a synthetic double-stranded fragment (1114 bp for in vitro retrohoming reactions, 1060 bp for in vivo retrohoming reactions) carrying the same junctions as the integrated molecules. The average of two to three technical replicates was used for every biological replicate. The total number of integrated molecules was estimated by correcting for the dilution factor. Because reverse transcription generates only one DNA strand across the junction, whereas the double-stranded standard templates from both of its strands in the first cycle, for the in vitro reactions every value reported was multiplied by two (Figure S16G). This was not applied to the quantification of in vivo data due to the impossibility of determining how many of these molecules have been converted to dsDNA in the cells. Retrohoming efficiency was calculated by dividing the estimated number of integrated molecules by the total number of target molecules present in the retrohoming reaction.

#### Analysis of DNA-binding Specificity from Nanopore Sequencing Experiments

##### Read processing

Following nanopore base-calling and assignment of reads to barcodes, individual sequences were extracted without any quality thresholding and were aligned against the expected retrohoming product using BLAST^7^. This product is composed of the invariant target sequence along with the intron inserted at its known integration site. A read contributed a valid count when at least 90% of the sequence aligned with the expected product and where the alignment spanned the entire intron, so that only reads crossing both integration junctions were retained. This ensured that only completed retrohoming events were quantified. The target sequence was extracted from the aligned read at the positions that are randomized in the template. Extracted sequences that contained a gap or were incomplete were rejected. Identical target sequences were then tallied, so that each barcode yielded one count for every distinct sequence observed in the randomized window. The matched input library was processed in the same way using the unreacted target sequence and was used to compute background position-specific base frequencies.

##### Sequence selection

High-affinity sequences were identified using a one-sided Poisson enrichment test with a threshold of P < 10^−3^. The base frequencies at each position within each randomized window was estimated using the count frequency of sequences in the matched input library. The background probability of a specific target sequence was then given by the product of its position-specific base frequencies. The expected count in the retrohoming experiments was given by this probability multiplied by the total product count. The P-value under this null model, for the observed count c and expected count λ, is given by the inclusive upper tail Pr[X ≥ c], where X is assumed to be a Poisson distribution with mean λ. The estimate did not include a correction for multiple hypothesis tests.

##### Motif logos

Consensus DNA-interaction specificities for the selected sequences were displayed using motif logos^8^. The logos represent the discrete probability distribution d(i, b) over a randomized window, indexed by i, for the b bases A, C, G and T. The height of the column represents the information content I(i) of bases at that position compared with uniform background probabilities

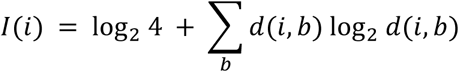

The information content is zero if the four bases are observed with equal frequency and two bits when a single base is observed in all recovered sequences.

The letters within each column are stacked in order of increasing height, so that the most probable base is uppermost, and the height h(i, b) of an individual letter is the share of the information content contributed by that base

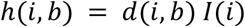

##### Clustering

The sequence clusters were obtained by fitting a mixture model to the selected sequence population. Each sequence was modelled as a draw from a finite mixture of independent categorical distributions^9^. A similar formulation is used for motif discovery from unaligned biological sequences^10^. Models were fitted with several different numbers, K, of cluster components. During fitting, every cluster is associated with a mixing weight and a probability for the four nucleotide bases across the randomized window. Observed read counts were used as weights, so a sequence that was counted many times contributed proportionally more than a sequence that was observed rarely.

Model parameters were estimated using expectation-maximization (EM)^11^ from five random initialization conditions. The fit with highest likelihood was retained for further analyses. The fitting algorithm ran until convergence or 2000 EM iterations.

##### Estimating cluster support with bootstrap resampling

The estimates of cluster stability were determined by bootstrap resampling^12^. In each of 200 draws the read count for each sequence was resampled multinomially in proportion to its observed count. The resampled counts were clustered using an identical procedure to that used to fit the source data. Each cluster in the source fit was compared with its best match in the resampled fit by the Jaccard coefficient, which represents the size of the sequence set intersections divided by the size of the union^13^. The sequences were weighted by their original count, and the reported support was the arithmetic mean of the Jaccard coefficient across draws.

##### Selecting the number of cluster components

The candidate models for each number K of cluster components were scored by the integrated complete likelihood (ICL) score^14^. The ICL score is based on the Bayesian information criterion^15^ but also includes an entropy term that penalizes components whose memberships overlap to reduce overfitting^16^. Bootstrap resampling was used to compare model fit quality as the difference between model ICL scores was frequently small and inconsistent in sign and magnitude across bootstrap resampled draws.

The model selection started with a single cluster and only added further clusters if they improved the model fit. To be preferred, a more complex model had to achieve a better ICL score in more than 50% of paired bootstrap draws. The separated clusters also had to have a Euclidean distance ratio that exceeded a threshold of 0.9. This restriction was included to prevent separation of closely related clusters that have distinct dinucleotide sequence recognition or other local non-linear effects^17^. Additional clusters were also rejected if the component with the lowest cluster weight contained fewer than 1% of the selected reads.

##### Detecting and validating DNA register shifts

Recognition of multiple motifs is a common property of DNA-binding proteins^17^. The sequence clustering algorithm represents an unbiased approach to identifying these effects, which can arise from several different mechanisms^18,19^. An additional method was developed to distinguish register-shifted motifs from other types of non-linearity such as dinucleotide sequence recognition.

Register-shifts represent invariant motifs that exist at fixed frame shifts with respect to each other. The algorithm for detecting register shifts involves three steps. In the first step, a scoring scheme was used for measuring the similarity of two motif logos. This scheme was then used to identify the best-scoring register offset, by sliding the motif logo of two distinct clusters through every whole-base offset up to three in both directions^20^. The test statistic is the distance between the maximal register score and the unshifted score. This provides a baseline that prevents false detection of offsets between motifs that are uninformative or have similar propensities across the window. In the third step a P-value for that improvement was estimated against a null model of motifs that carry the same information content as the observed clusters but are otherwise random.

##### Scoring cluster similarity

Each cluster was represented by its information-content motif logo^8^, which is derived from the base probability distribution *d(i, b)* for the bases *b* in A, C, G and T over the extent of the window, indexed by *i*. This can be used to compute the information *I(i)* for the position, which varies between zero bits for uninformative positions and two bits for full conservation. The agreement, *a(i)*, is the probability mass that is shared by the two positions^21^, summed over the four bases

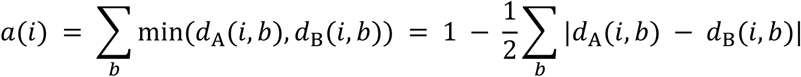

The agreement score is one when the two distributions are identical and zero when they are disjoint. Each position was weighted by the geometric mean of the information at that position

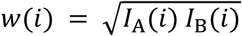

and the positions were combined into a single score as a weighted mean

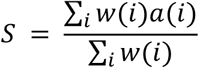

The weighting gives greater emphasis to positions that are strong determinants of the motif’s binding specificity and de-emphasizes uninformative positions.

The score was computed for pairs of logos at every whole-base offset j from minus three to plus three, with the two sums running over only the positions that overlap, of which there are L - |j| for a window of length L. The offset j with the highest score, taken over the non-zero offsets only, is the register shift reported for that pair, and the test statistic is given by

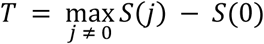

##### A null model for the test statistic

The null model tests whether offset motifs have greater similarity than would be expected by chance. For each assay, we generated a random set of motifs, termed decoys, with the same window length. Each decoy was generated by pooling the per-position information values from all clusters identified in the assay and then sampling with replacement one value per position. The sampled values were scaled to match the mean total information across the observed clusters, with the maximum information at each position capped at 1.999 bits. One of the four bases was chosen randomly and with equal probability as the dominant base at that position. The probability of the dominant base was set to match the specified information content, with the remaining probability divided equally among the remaining three bases.

For assays with two fitted clusters, we calculated T for the observed pair and for 4,000 pairs of independently generated decoy motifs. The P-value is the fraction of decoy pairs whose T is greater than or equal to the observed value.

##### Parameters

The table below lists the fitting, resampling and register-test parameters and threshold parameters. These apply to the complete Cell figure input set, including the all-fine-tuning export. The target-sequence P-value filter was applied before fitting and bootstrap resampling.

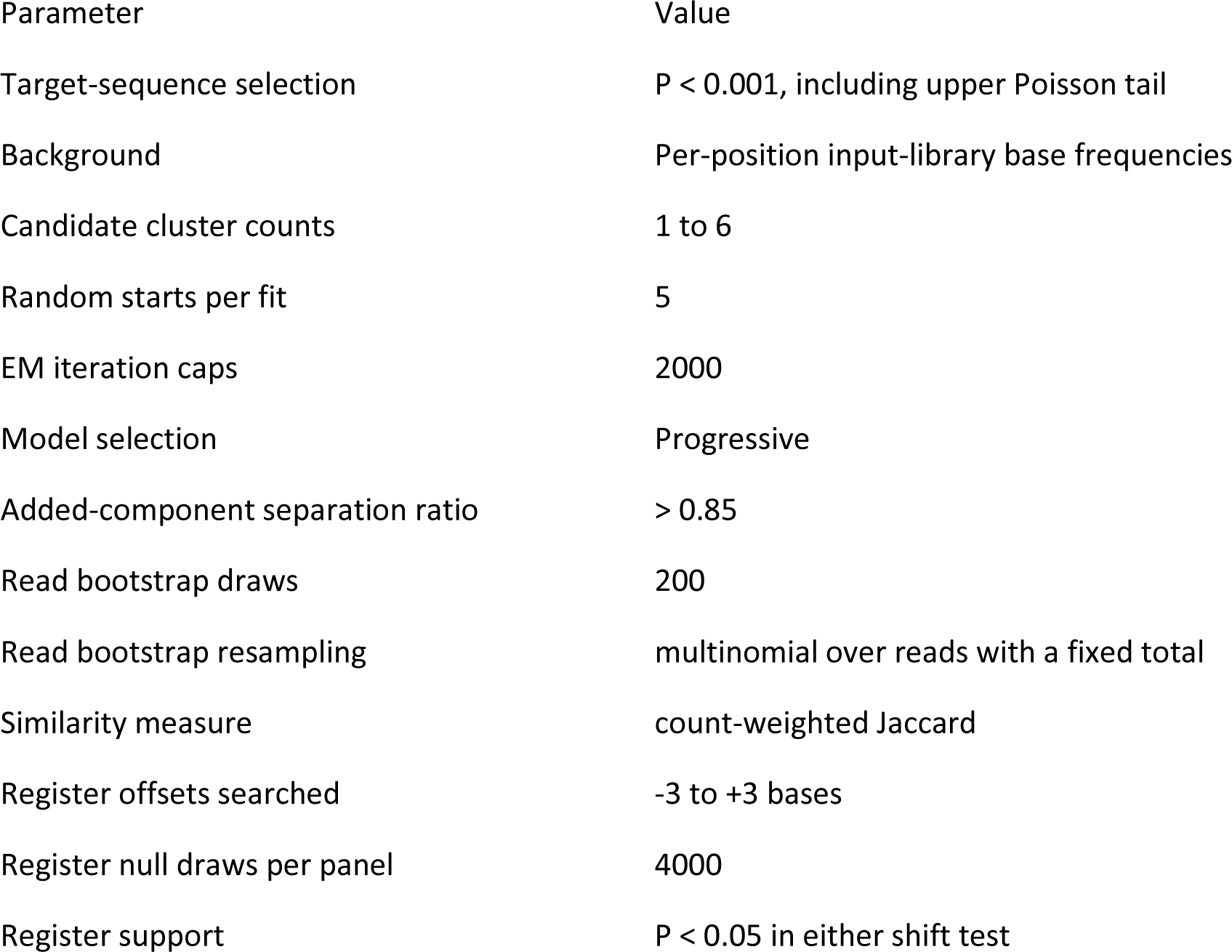

#### Motif database and target recoding

##### Construction of the motif database

Reads from every IEP and intron library experiment were processed as described under *Read processing*, giving for each barcode a count for every distinct sequence recovered in the randomized window. Counts were corrected for the composition of the matched input library, and sequences falling below a per-sample count threshold, set from the shape of the count distribution for that sample, were treated as background and discarded.

For each variant the surviving sequences, weighted by count, were used to derive a position weight matrix across the randomized window, from which a consensus motif was read off in degenerate (IUPAC) notation. The consensus was then used to distinguish positions that the variant genuinely specifies from positions at which it has no base preference: a uniform background was subtracted from the matrix, and each position was classified according to the extent to which its base distribution departed from uniform, those remaining close to uniform being masked as unconstrained. The recovered sequences were re-counted considering only the specified positions, so that reads differing solely at positions the variant does not discriminate were aggregated. This gave, for each variant, a motif table: a set of distinct recognised sequences over the specified positions, each with a summed read count, and each additionally expressed as a fraction of the count of that variant’s most abundant entry.

Activity was the total number of retrohoming reads recovered for the variant relative to the number recovered for the reference IEP, assayed against the same target library in the same experiment. Intron libraries were scored separately, as the count of each IBS2–EBS2 pair relative to that of the wild-type pair in the same sample. The motif tables, consensus motifs and activities were stored in a SQLite database indexed by the protein or intron region characterised, referred to below as the motif database.

##### Target-site selection

Candidate insertion sites were identified with in-house software by scanning a query sequence at single-base steps on both strands. At each step the target footprint was defined about the notional splice site at that position and partitioned into the sub-regions contacted by the 5′ protein regions, by EBS2, by EBS1 and by the 3′ protein region, positions being numbered relative to the splice site such that the base immediately 3′ of it is +1 and the base immediately 5′ of it is −1, there being no position 0. A site was accepted only where the motif database returned, together, a 5′-region variant and a 3′-region variant whose recorded motif or consensus motif matched the target at the corresponding sub-region, and an intron EBS2 characterised as supporting integration with the target’s IBS2; matching was exact under IUPAC, with no mismatch tolerance. Accepted sites were ranked by the product of the corresponding activity terms, each capped at 2.5-fold over the reference, so that combinations expected to give more efficient integration ranked above those expected to give less.

##### Design of recoded IEPs and introns

For each selected combination, the variant 5′ and 3′ protein regions were grafted onto the backbone IEP in place of the corresponding backbone regions, at fixed junctions, giving a single non-naturally occurring IEP that carries the 5′ specificity of one donor and the 3′ specificity of the other. The coding sequence was generated by aligning the chimera to the backbone protein with MUSCLE (v5.2) and retaining the backbone codon wherever the residue was unchanged, so that each construct differs from the validated backbone at the minimum number of bases, and was then checked and, where necessary, adjusted by synonymous substitution alone to remove features expected to impair expression: cryptic splice sites, out-of-frame internal initiation codons, a defined set of restriction and polyadenylation motifs, and local GC content outside 0.45 to 0.65, with the original sequence being retained where no synonymous adjustment was available.

The intron template was recoded by substituting the target segments flanking the insertion site, together with EBS1 and EBS2. EBS1 is fully determined by the chosen site, being the reverse complement of the target bases that pair IBS1. EBS2 is not: rather than taking the Watson–Crick complement of IBS2, the highest-activity partner of the target’s IBS2 recorded in the motif database was used.

### 4. ADDITIONAL RESOURCES

Not applicable.

### 5. KEY RESOURCES TABLE

| REAGENT or RESOURCE | SOURCE | IDENTIFIER |
|---|---|---|
| Bacterial and virus strains |  |  |
| *E. coli* BL21(DE3) | NEB | Cat#C2527H |
| 5-alpha competent cells | NEB | Cat#C2987H |
| Chemicals, peptides, and recombinant proteins |  |  |
| See Table S4 Protein sequences used in this study | N/A | N/A |
| Q5 High-Fidelity 2X Master Mix | New England Biolabs | Cat#M0492L |
| NEBuilder HiFi DNA Assembly Master Mix | New England Biolabs | Cat# E2621L |
| ATTO-540Q-ddATP | Jena Bioscience | Cat#NU-1612-540Q |
| Terminal (deoxynucleotidyl) transferase | New England Biolabs | Cat#M0315L |
| RNase inhibitor | This paper | N/A |
| RNase A | New England Biolabs | Cat#T3018-2 |
| Blunt/TA Ligase Master Mix | New England Biolabs | Cat#M0367L |
| Native Barcoding Kit 24 V14 | Oxford Nanopore Technologies | Cat#SQK-NBD114.24 |
| Native Barcoding Kit 96 V14 | Oxford Nanopore Technologies | Cat#SQK-NBD114.96 |
| MinION/GridION Flow Cell (DNA) | Oxford Nanopore Technologies | Cat#FLO-MIN114 |
| NEBNext dA-Tailing Module | New England Biolabs | Cat#E6053S |
| NEBNext Ultra II End Repair/dA-Tailing Module | New England Biolabs | Cat#E7546S |
| Deoxynucleotide (dNTP) Solution Set | New England Biolabs | Cat#N0446S;N0447 |
| Lipofectamine 3000 Transfection Reagent | Thermo Fisher Scientific | Cat# L3000008 |
| Opti-MEM I Reduced Serum Medium | Thermo Fisher Scientific | Cat# 31985-070 |
| DMEM, high glucose | Thermo Fisher Scientific (Gibco) | Cat# 11965-092 |
| GlutaMAX Supplement | Thermo Fisher Scientific | Cat# 35050-038 |
| Fetal bovine serum, heat-inactivated | Thermo Fisher Scientific (Gibco) | Cat# A5256801 |
| Penicillin-streptomycin | Thermo Fisher Scientific (Gibco) | Cat# 15140-122 |
| TrypLE | Thermo Fisher Scientific | Cat# 12604-021 |
| SafeView nucleic acid stain | NBS Biologicals | Cat# NB5-SV1-C |
| GeneRuler 1 kb Plus DNA Ladder | Thermo Fisher Scientific | Cat# SM1331 |
| LB agar | Melford | Cat# L24021 |
| 3x LB medium | Melford | Cat# L24041 |
| LB medium | Formedium | Cat# LMM0125 |
| TB | Melford | Cat# T15000 |
| Carbenicillin | Melford | Cat# C46000 |
| Kanamycin | Melford | Cat# K22000 |
| IPTG | Melford | Cat# I56000 |
| SOC | NEB | Cat# B9020S |
| Lysozyme | Thermo Fisher Scientific | Cat# 89833 |
| EDTA-free Protease Inhibitor Cocktail | Pierce | Cat# A32965 |
| Ni-Agarose EXTRACHEL resin | ABT | Cat# 6RREXCH-500 |
| Cytiva™ Protein Select™ resin | Cytiva™ | Cat# 17542101 |
| Bradford Plus Protein Assay | Pierce™ | Cat# 23238 |
| Quick Coomassie Stain | New Biotech | Cat# NB-45-00078-1L |
| Invitrogen™ Novex™ Tris-Glycine Mini Protein Gels, 14% | Invitrogen | Cat# XP00145Box |
| Tris-Glycine running buffer | Invitrogen | Cat# LC26755 |
| Glyoxal, 40% w/w aq. soln. | Thermo Fisher Scientific | Cat# A16144 |
| RNA Storage Solution (1 mM sodium citrate pH 6.5) | Thermo Fisher Scientific (Invitrogen) | Cat# AM7001 |
| MassRuler DNA Loading Dye (6X) | Thermo Fisher Scientific | Cat# R0621 |
| Novex TBE-Urea 6% PAGE gel | Thermo Fisher Scientific (Invitrogen) | Cat# EC68652BOX |
| Critical commercial assays |  |  |
| EasyPure PCR Purification Kit | TransGen Biotech | Cat#EP101-02 |
| Qubit dsDNA HS Assay Kit | Thermo Fisher Scientific | Cat#Q32854 |
| Qubit Assay Tubes | Thermo Fisher Scientific | Cat#Q32856 |
| RNAClean XP beads | Beckman Coulter | Cat#A63987 |
| EasyPure Plasmid MiniPrep Kit | TransGen Biotech | Cat# EM101-02 |
| DreamTaq Hot Start Green PCR Master Mix (2X) | Thermo Fisher Scientific | Cat# K9022 |
| TURBO DNase | Thermo Fisher Scientific | Cat# AM2238 |
| Luna Universal Probe qPCR master Mix | NEB | Cat# M3004 |
| QuickExtract DNA Extraction Solution | LGC | Cat#QE09050 |
| Experimental models: Cell lines |  |  |
| Human: HEK293 | ATCC | Cat# CRL-1573 |
| Oligonucleotides |  |  |
| See Table S2 Sequences of oligonucleotides and DNA retrohoming target amplicons used in this study | N/A | N/A |
| Recombinant DNA |  |  |
| See Table S1 Plasmids used in this study | N/A | N/A |
| Software and algorithms |  |  |
| MinKNOW v25.09.16 | Oxford Nanopore Technologies | N/A |
| BLAST | See reference (7) | https://blast.ncbi.nlm.nih.gov/; version TBC |
| Custom mixture-model clustering, bootstrap and register-shift code | This paper | N/A |
| Warp | See reference (2) | https://github.com/warpem/warp |
| cryoSPARC v5.0.2 | See reference (3); Structura Biotechnology | https://cryosparc.com/ |
| Coot 0.9.8.95 | See reference (4) | https://www2.mrc-lmb.cam.ac.uk/personal/pemsley/coot/ |
| Phenix 1.21 | See reference (5) | https://phenix-online.org/ |
| EPU, with aberration-free image shift (AFIS) | Thermo Fisher Scientific | N/A |
| Other |  |  |
| Table S5 RNA sequences used in this study | N/A | N/A |
| Chemistry technical document, v1 rev. AV | Oxford Nanopore Technologies | CHTD_500_v1_revAV_24July2026, https://nanoporetech.com/document/chemistry-technical-document |
| Protocol, Ligation sequencing amplicons - Native Barcoding Kit V14 (SQK-NBD114.24) | Oxford Nanopore Technologies | https://nanoporetech.com/document/ligation-sequencing-amplicons-native-barcoding-v14-sqk-nbd114-24 |
| MinION Mk1B sequencing device | Oxford Nanopore Technologies | Cat#MIN-101B |
| MinION Mk1D sequencing device | Oxford Nanopore Technologies | Cat#MIN-101D |
| Qubit 2.0 Fluorometer | Thermo Fisher Scientific | Cat#Q32866 |
| DeNovix DS-11 spectrophotometer | DeNovix | N/A |
| Implen NP80 NanoPhotometer | Implen | N/A |
| T8-ISO, Isothermal Amplification Instrument | Axxin | N/A |
| ChemiDoc Imaging System | Bio-Rad | N/A |
| BioLite 24-well multidish, cell culture treated | Thermo Fisher Scientific | Cat# 130186 |
| Disposable Columns 2 mL | Thermo Scientific | Cat# 29920 |
| Slide-A-Lyzer™ MINI Dialysis Devices, 10K MWCO | Thermo Scientific | Cat# 69572 |
| 37 °C shaking incubator | SciQuip | Incu-shake TL6-5R |
| 20 °C shaking incubator | Lab Companion | IS-971R |
| Spectrophotometer | BioChrom | NovaSpec Plus |
| Sonicator | FisherBrand | FB120 |
| Sonicator tip (1/8 inch) | Bandelin | 422-A |
| Titan Krios transmission electron microscope | Thermo Fisher Scientific | Model#G3 |
| Selectris X energy filter | Thermo Fisher Scientific | N/A |
| Falcon 4i direct electron detector | Thermo Fisher Scientific | N/A |
| Vitrobot | Thermo Fisher Scientific | Model#MarkIV |
| UltrAuFoil R1.2/1.3 Au 300-mesh grids | Agar Scientific | Cat# AGS187-3 |

