## Supplementary information for "Structural basis for reprogramming group II intron target specificity"

### Supplementary materials

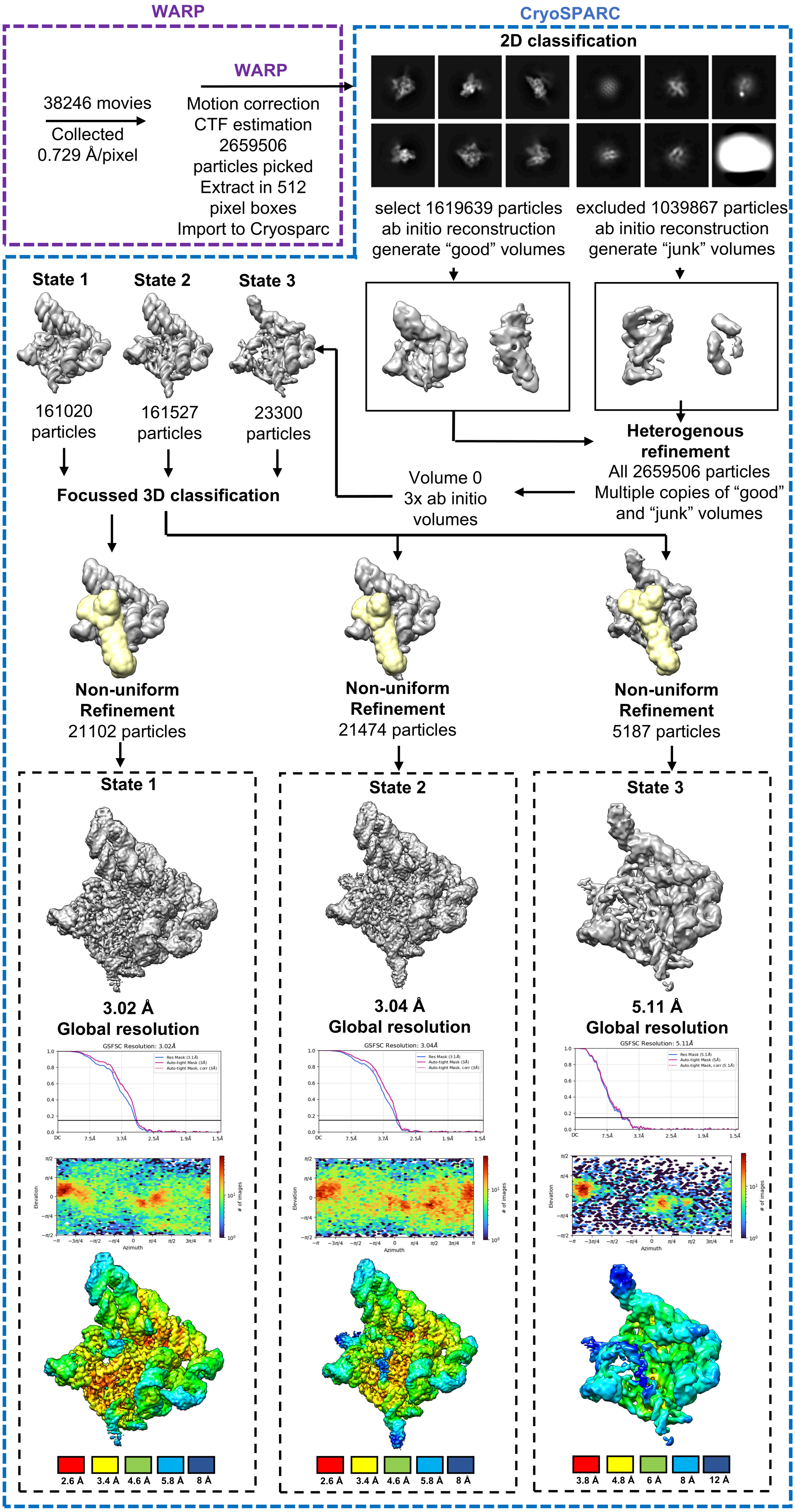


OK

**Figure S1. Cryo-EM data processing workflow for the Voyager-1 complexes.** Single-particle processing from data collection to the final maps for states 1, 2 and 3; full details are given in STAR Methods. Steps carried out in WARP are boxed in purple and those in cryoSPARC in blue; particle numbers are given at each stage. Yellow marks the DNA and HNH regions on which the focussed 3D classification was centred.
For each state the boxed panel shows, from the top: the final map; the gold-standard FSC curve, for the resolution mask and for the auto-tight mask with and without correction, with the global resolution quoted above it; the distribution of particle viewing directions, plotted as azimuth against elevation and coloured by the number of images; and the map coloured by local resolution on the scale beneath. The local-resolution scale differs for state 3.

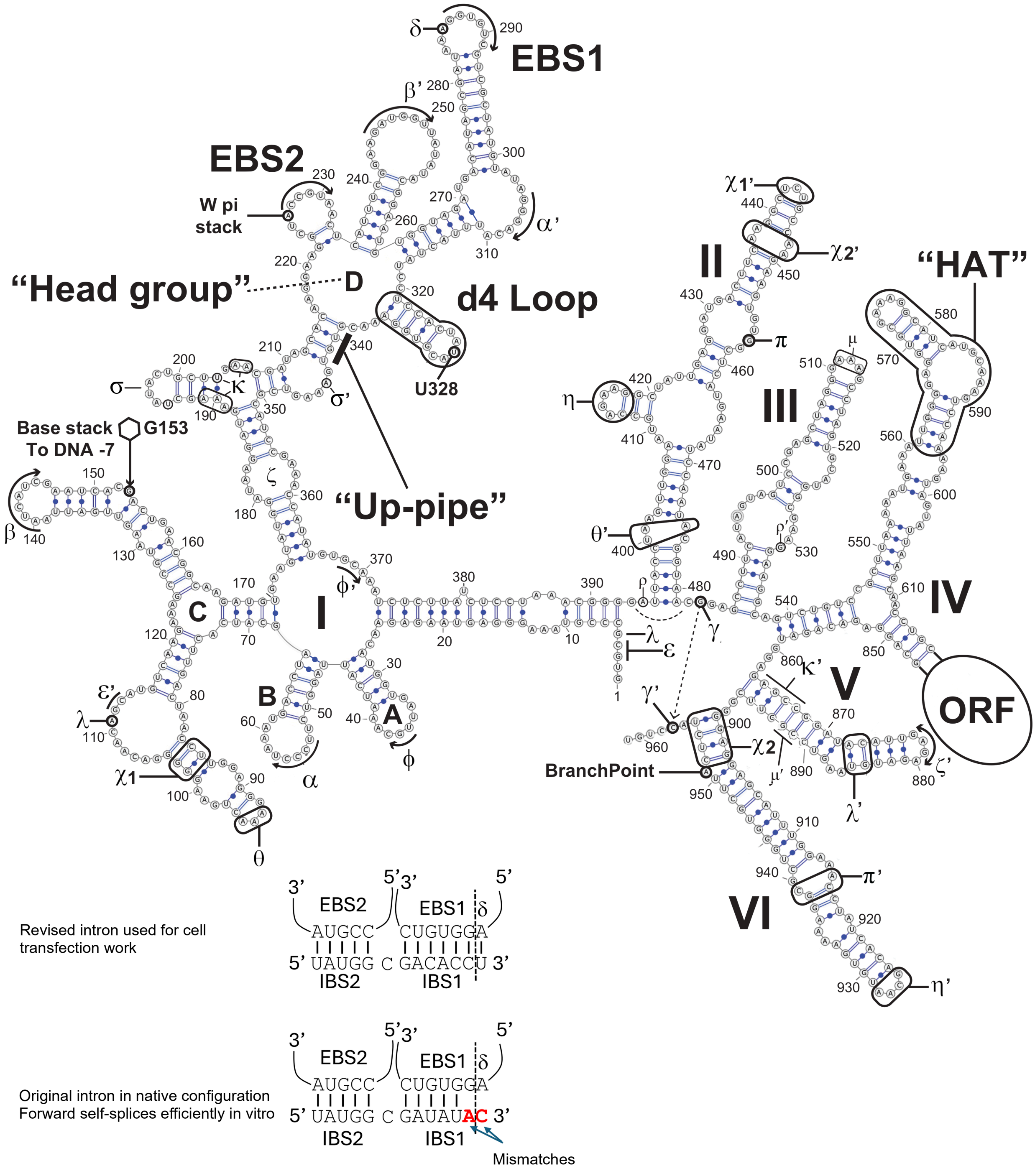


**Figure S2. Secondary structure of the Voyager-1 intron.** Secondary structure of the Voyager-1 (class IIA) intron as present in the construct used for cryo-electron microscopy, drawn in the conventional group II scheme. The six domains are labelled I–VI and the subdomains of domain I are labelled A–D (Ia–Id).

Nucleotides are numbered from the intron 5′ end (G1). Watson–Crick G–C pairs are drawn as a double blue line; A–U pairs and G·U wobbles as a single blue line with a filled blue circle.

The intron-encoded protein (IEP) open reading frame is carried within domain IV. The construct retains only a 230-nucleotide remnant of that reading frame, and the domain IV hairpin to the main ORF segment is drawn collapsed to the five stem base pairs, with the deleted interior represented by the labelled bubble (ORF); the bubble is a placeholder and not drawn to scale. All numbering in this figure, and throughout the paper, referencing intron base positions, interactions and function refers to this 961-nucleotide construct (STAR methods). In cell delivery experiments introns with a further deletion of the ORF were used, but specific base positions are not discussed herein.

Features that engage the DNA target (EBS1,EBS2, delta) as well as some that engage the IEP are annotated directly (d4 loop, HAT). Arrows mark the exon-binding sites EBS2 (227–231) and EBS1 (286–291). The branch-point adenosine (A951, domain VI) is ringed and labelled. U328 marks the tip of the class IIA-specific domain I d4 loop. “W pi stack” marks A226, the base immediately preceding EBS2 on which IEP W418 stacks as the RNA turns to present the EBS2 nucleotides. G153, drawn as a ringed base, stacks onto the DNA base at position −7. The “head group”, the “up-pipe” are informal names used in the text for these regions of domain I, and “HAT” denotes the domain IVa hairpin that forms the principal high-affinity intron–IEP anchor.

Greek symbols denote tertiary interactions, each partner carrying the unprimed and primed form of the same letter (α–α′, β–β′, γ–γ′, δ, ε–ε′, ζ–ζ′, η–η′, θ–θ′, κ–κ′, λ–λ′, μ–μ′, π–π′, ρ–ρ′, σ–σ′ and φ–φ′); heavy outlines enclose the nucleotides that participate in the contact so indicated, except in domain V, where κ′ and μ′ converge on the same C867:G892 base pair from opposite strands and are therefore drawn as brackets lying alongside the strand each one reads. ε at G3/C4 pairs with ε′ at 112/113 at the 5′ end of the intron, and λ at G5 completes a three-way contact with λ at 111 and λ′ at 884/885. π–π′ is the G457:C939 pair joining domain II to domain VI. κ runs from A189–A191 and U203/A205/A206 in domain I to G865, C866 and C867 of domain V and to G893 and U895 opposite them; μ runs from the domain III terminal loop (A511–G514) to G868, C891 and G892. σ–σ′ marks a contact not previously described: A344 inserts into the 194–201 hairpin and makes a reverse-Hoogsteen pair with U194, also contacting A197, C198, U199, G200 and C201; the ring marks U194. ρ–ρ′ is the stack of A394 on G531, not previously reported in a class IIA intron but corresponding to the contact termed p in class IIB introns (lettered ρ here, as p and π are both in use). The interactions φ, μ, ρ, σ, χ1 and χ2 are identified here and are developed in Supplementary Note S3: φ pairs U36:A370, G37:C369 and C38:G368 between the domain Ia turn and the domain I hub; χ1 pairs G103 in domain Ic with U442 in the long-form domain II, a contact absent from Ll.LtrB; and χ2 marks the backbone-mediated ribose zipper between domain II and domain VI. These are presented as observations rather than established interactions.

At the foot of the figure, two rows show the exon-binding sites paired with the intron-binding sites of the flanking exon: intron 3′→5′ above, exon 5′→3′ below, ticks marking pairs and the dashed line the splice site; the single unpaired exon position between IBS2 and IBS1 is −7, the base onto which G153 stacks. In the native configuration — the intron used for the cryo-electron microscopy work — IBS2 pairs EBS2 throughout, but the two positions reading the splice junction do not: G286, the 5′-most nucleotide of EBS1, and δ (A285) face mismatched exon bases, shown in red. That intron still self-splices efficiently in the forward direction in vitro, so the structures reported here are unaffected. For the work in cells the flanking exon was revised so that all twelve positions pair, restoring both the EBS1 and δ interactions; both are shown.

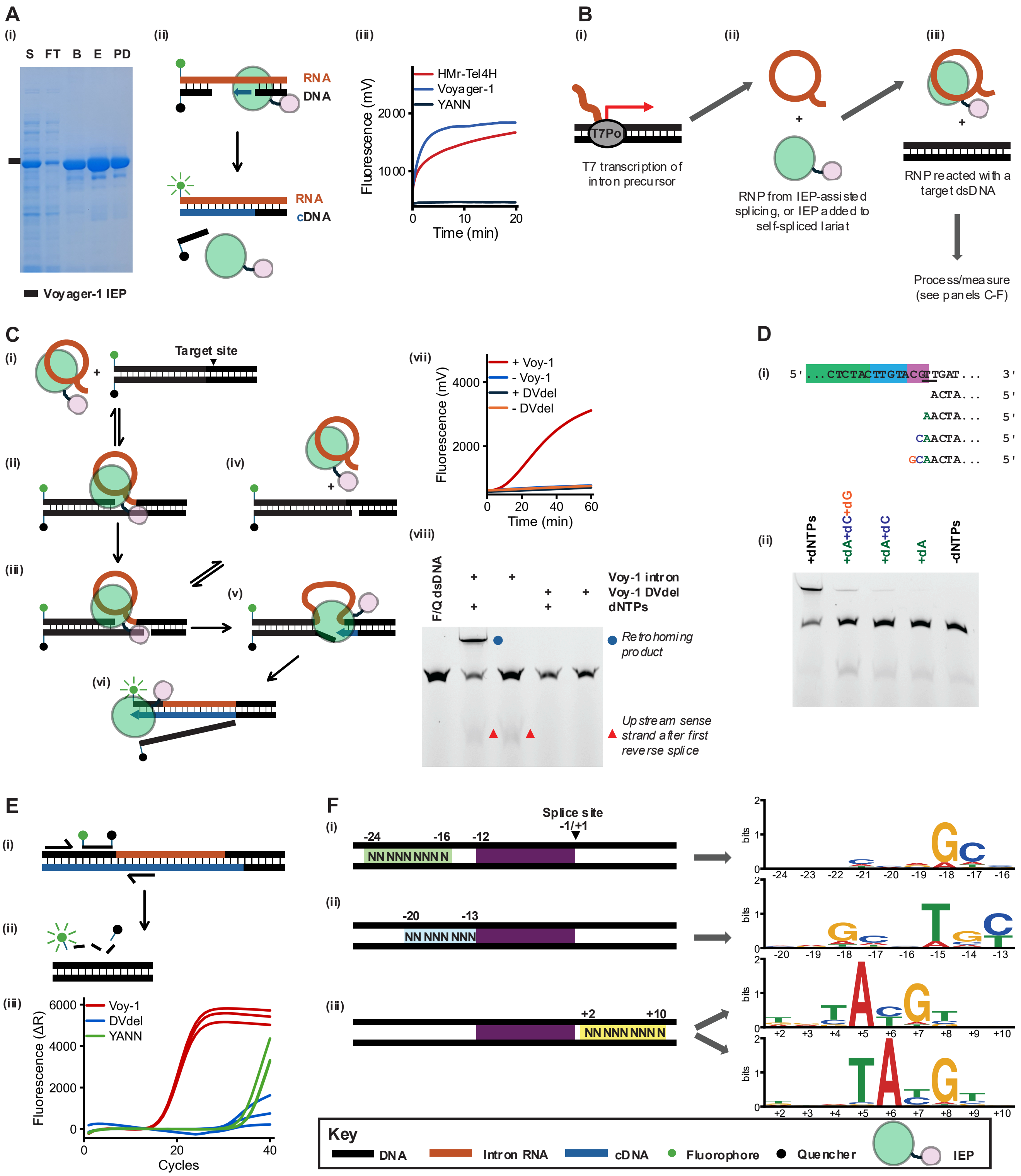


**Figure S3. In vitro reconstitution of Voyager-1 retrohoming: IEP purification, reverse transcription, dsDNA invasion and target-site specificity.** All reactions used purified Voyager-1 protein and T7 polymerase-transcribed intron RNA to assemble ribonucleoprotein (RNP) particles, as well as a synthetic double-stranded DNA (dsDNA) target. Element colours are given in the Key at the foot of the figure: black, DNA; orange, intron RNA; blue, cDNA; green circle, fluorophore; black circle, quencher; green and pink shape, the intron-encoded protein (IEP), the pink lobe denoting the endonuclease domain.

(A) Purification and reverse-transcriptase activity of the Voyager-1 IEP. (i) Coomassie-stained SDS-PAGE across the purification: S, soluble lysate; FT, flow-through; B, bound (on-resin); E, eluate; PD, post-dialysis. The bar marks the Voyager-1 IEP. (ii) Schematic of the fluorescence reverse-transcription assay. A fluorophore-labelled DNA primer and a quencher-labelled downstream DNA oligonucleotide are annealed to an RNA template; primer extension by the IEP (arrow) displaces the quencher-bearing strand, separating quencher from fluorophore and yielding cDNA with a concomitant rise in fluorescence. (iii) Reverse-transcription time courses (fluorescence, mV) for the Voyager-1 IEP, the HMr-Tel4H IEP, and the YANN RT domain variant, which is inactive throughout.

(B) Assembly and assay workflow. (i) T7 RNA polymerase transcription of the intron precursor. (ii) RNP particles generated either by IEP-assisted splicing or by adding purified IEP to self-spliced lariat RNA. (iii) RNPs reacted with a target double-stranded DNA. Products were then processed and measured as shown in panels C to F.

(C) Retrohoming into a fluorophore/quencher-labelled dsDNA target. (i–vi) Model of the reaction. (i) RNP and a dsDNA substrate carrying a fluorophore on the 5′ end of the sense strand and a quencher on the 3′ end of the antisense strand; the target site is arrowed. (ii) Reversible recognition and reverse-splicing into the target. (iii) First reverse-splicing into the target. (iii) Cleavage of the antisense strand by the IEP endonuclease domain generates the 3′-OH primer that the reverse transcriptase engages. (iv) Reversible splicing out of the intron can release the RNP from the endonuclease cleavage product. (v) Extension of the primer by RT, initially through DNA-templated DNA synthesis, traps the reverse-spliced product. (vi) Reverse transcription produces cDNA and ultimately displaces the quencher-bearing strand, relieving quenching and generating fluorescence. (vii) Fluorescence time course (mV) for wild-type Voy-1 RNP and for intron domain V deletion (DVdel) RNP, ± dNTPs; only wild-type RNP supplied with dNTPs gives a signal. (viii) Denaturing gel electrophoresis of the same reactions from (vii), after column-purification and glyoxal-treatment. Lane 1, fluorophore/quencher dsDNA alone (F/Q dsDNA); lanes 2 and 3, Voy-1 intron RNP ± dNTPs; lanes 4 and 5, Voy-1 DVdel RNP ± dNTPs. The full retrohoming product (blue circle) requires both an intact intron and dNTPs, whereas the faster-migrating upstream sense strand released by the first reverse-splicing step (red triangles) forms with the intact intron irrespective of dNTPs, and not with DVdel.

(D) Reverse transcription initiates in register at the target-site junction. (i) Intron RNA (green) retrohomed into target DNA (blue; with templating positions +6 to +8 vs the splice site blocked in pink); nascent cDNA products are shown beneath. Supplying restricted dNTP subsets stalls synthesis at the positions predicted by the template: +dA adds a single A, +dA+dC adds CA, and +dA+dC+dG adds GCA. (ii) Denaturing gel electrophoresis of column-purified and glyoxal-treated reactions that had been supplied with the complete dNTP set, with the indicated subsets, or with no dNTPs. Only the complete set yields the fully extended retrohoming product.

(E) Junction-specific qPCR assay for retrohoming product. (i) A forward primer in the target DNA and a reverse primer within the intron cDNA flank the insertion junction. (ii) Hydrolysis of a dual-labelled hydrolysis probe during amplification separates fluorophore from quencher. (iii) Amplification curves (ΔR against cycle number, three technical replicates per condition) for wild-type Voy-1, DVdel and YANN RNPs. Only wild-type gives specific amplification; DVdel and YANN show only late, non-specific signal.

(F) Selection of target-site preferences from randomised libraries. Three double-stranded target libraries were built on a fixed core for RNA recognition (positions −12 to +110, purple), randomising (i) positions −24 to −16 (green, N9), (ii) positions −20 to −13 (blue, N8) and (iii) positions +2 to +10 (yellow, N9), numbered relative to the insertion site (−1/+1, arrowhead). Sequence logos (bits) summarise the sequences recovered after reaction with Voyager-1 RNPs. The upstream libraries select G at −18 with C at −17 (i), and T at −15 with C at −13 (ii). The downstream library (iii) yields two populations carrying the related YAYGY motif offset by one nucleotide: YAYGY at +4 to +8 (upper logo) and TAYGY at +5 to +9 (lower logo).

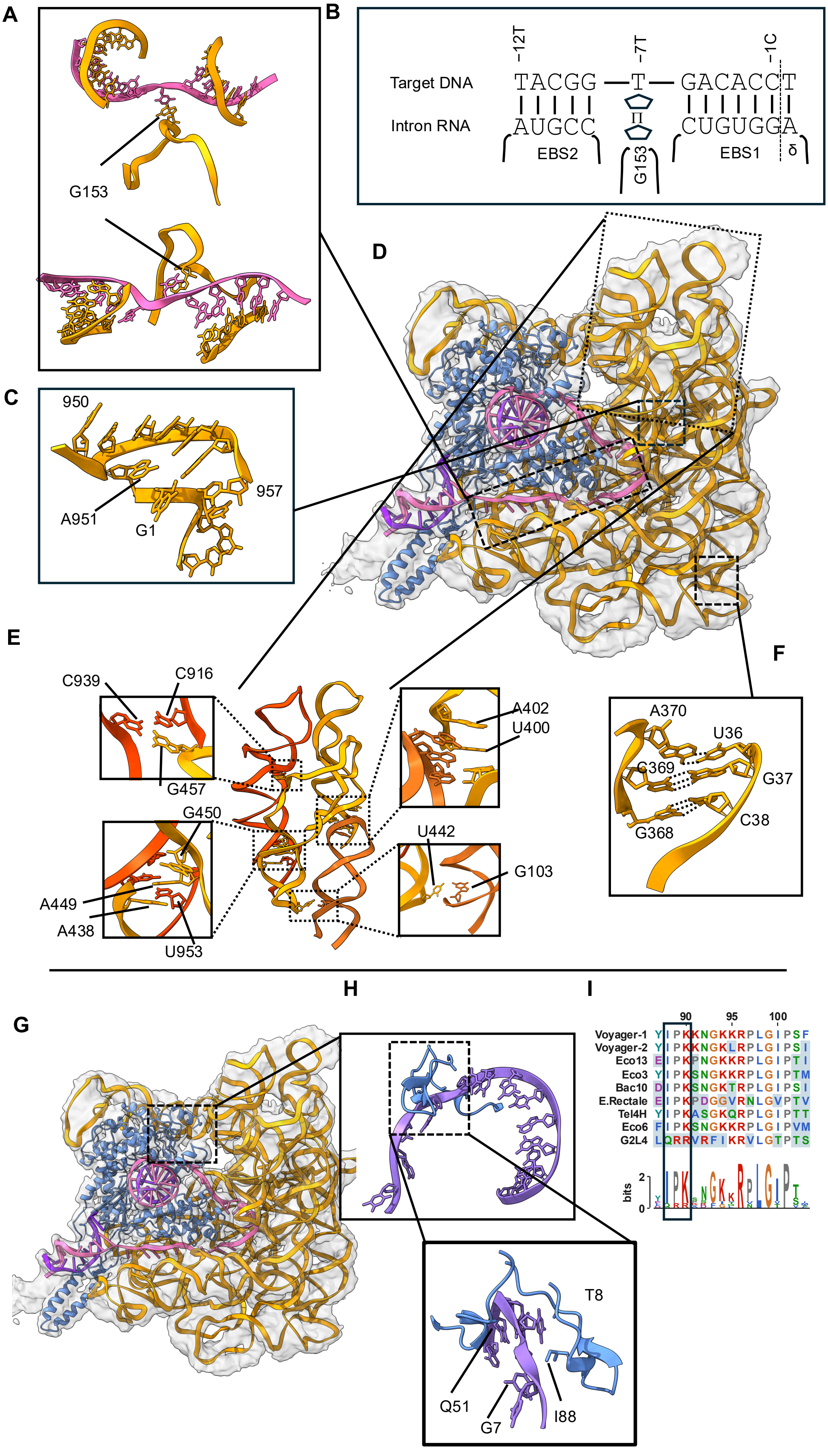


**Figure S4. Voyager-1 cryo-EM model novel RNA contacts A-F, and Voyager-2 contacts spliced ssDNA entering polymerase.** Throughout, intron RNA is orange/gold, the IEP is blue and target DNA is pink in (A) and (D) and purple in (G) and (H). Transparent grey surfaces in (D) and (G) are the cryo-EM density; black boxes and leader lines locate each inset within the overall view.

(A) Two orthogonal views of intron nucleotide G153, where the intron backbone twists to present the base for pi stacking with the -7 sense target DNA base. Upper, view across the duplex axis; lower, view along it. Bases are drawn as sticks.

(B) Pairing scheme for the same region, target DNA sense strand above (5′ to 3′, positions −12, −7 and −1 marked) and intron RNA below. Vertical ticks are Watson–Crick pairs. EBS2 pairs the target over −12 to −8 and EBS1 over −6 to −1; the intervening −7 T is not read by an EBS nucleotide but is accommodated by pi stacking to intron G153 (pentagon), which occupies that column from the intron side. δ marks the intron nucleotide pairing the −1 C.

(C) The domain VI hairpin and the intron 5′ end, nucleotides 950–957, with the branch-point adenosine A951 and intron G1 labelled.

(D) Overall view of the complex, model in density, oriented to show where the insets of (A), (C), (E) and (F) sit. Boxes and solid leader lines mark each.

(E) Long-range tertiary contacts between the two helical stacks boxed in (D), drawn with the two stacks in contrasting shades. Dotted boxes on the central view locate four insets, each naming the nucleotides involved.

(F) A three-base-pair long-range duplex between the 36–38 and 368–370 segments — G368:C38, C369:G37 and A370:U36 — with hydrogen bonds as dotted lines.

(G) The Voyager-2 complex macroscopic model with the box marking the DNA–IEP interface enlarged in (H)(high resolution model for entire complex not presented herein).

(H) The interface boxed in (G): the target DNA (purple) packed against a β-hairpin of the IEP (blue). The lower inset enlarges the contact itself. The DNA nucleotides G7 and T8 , and protein residues Q51 and I88 are indicated, all four side chains and bases drawn as sticks.

(I) Alignment of the IEP segment spanning residues for Voyager-1, Voyager-2, Eco13, Eco3, Bac10, E. rectale, Tel4H, Eco6 and G2L4, residues coloured by identity and numbered above. The sequence logo beneath gives the information content in bits at each column. The box encloses I88–K90, the segment that contributes I88 to the DNA contact in (H), and shows that the isoleucine and the proline–lysine that follow it are invariant across the set.

**Detailed notes on additional intron interactions reported here as part of Figure S4:**

**Other previously unreported intron-intron and intron-protein interactions**

Over time studies have contributed previously unreported RNA, or RNA-protein, interactions. We noted several new possible interactions worthy of attention based on our structures which may further a more complete picture of GII intron architecture.

**A proposed novel intron clasp.** We inferred existence of a previously unreported Watson–Crick interaction between the turn region of intron domain Ia and the opposing side of the domain I hub, hereafter the (GIIA) phi (φ) interaction (Figure S2, Figure 3F).

φ is likely to be a discrete three-base-pair duplex. U36 pairs A370, G37 pairs C369 and C38 pairs G368, in antiparallel register. The proposed register terminating sharply on both sides .

The interaction does sit in a region of below-average order however the pairing geometry appears nonetheless unambiguous, and we note the same overall close approach of equivalent RNA segments in deposited LtrA structures (for example 8H2H), albeit no specific interaction had been modelled or tested before

In support of this potential interaction, deeper phylogenetic and homology analysis of class IIA introns in our database provides further support that the proposed interaction would likely be widespread as appropriate 3 residue, or greater, pairings are generally evident. This interaction would neatly explain the previously undefined function of the DIa1 loop in supporting structural integrity to the folded intron in vivo.

**A new previously reported novel non-Watson–Crick domain II/domain I interactions to add.**

Voyager-1 intron maintains the previously described interaction between domain II and domain I, the q- q ‘ interaction (theta-theta’) (Michel et al., 1989; Costa et al., 2000; Toor et al., 2008). In addition, we designate as chi 1 (χ1) another interaction between the distal end of Voyager-1 domain II and domain IC, adjacent to but distinct from (θ) (Figure S2, Figure 3E). χ1 resolves as a single hydrogen bond in Voyager-1 between U442 and G103, at below 3 Å, with a second, longer possible contact to G104. This novel interaction is not present in LtrB. Voyager-1 contains an extended domain II sequence in comparison; thus we categorize LtrB as having a short-form domain II, and the Voyager-1 intron as having a long-form domain II. Analysis of a larger set of class IIA introns indicates that shorter and longer forms of domain II appear roughly evenly distributed in nature. Conservation of the nucleotides at the end of the long-form stem, i.e. including U442 and adjacent residues, are not however well-conserved exhibiting variations in the length and sequence of this turn. Thus, the criticality of this interaction may vary with modest total contribution to domain interactions. Consistent with this we tested mutations in this residue and observed a modest, but detectable, decrease in retrohoming efficiency.

**The domain II extension packs against domain VI.** The Voyager-1 intron maintains two previously described interfaces between domain II and domain VI, the h- h ' (eta-eta’) and the p- p ' interactions (Michel et al., 1989; Costa et al., 2000). The p- p ' interaction is base-mediated and sequence-dependent, and the h- h ' interaction involves a classical tetraloop (G925-A928) interaction with a receptor helix (C412-C420). We note here a new previously unreported interfaces between domain II and domain VI in an extended RNA segment.

Domain II appears to make a second set of contacts with domain VI (Figure 3E). It involves residues 438, 439, 449, 450 and 451 pressed against domain VI residues 901–904 and 952–954, dominated by four putative and reciprocal 2′-hydroxyl pairs — 449:952 (2.83 Å), 439:954 (2.91 Å), 450:903 (2.94 Å) and 438:901 (3.50 Å) — together with 2′-OH to ring-oxygen contacts this is designated a novel interaction χ2. This appears to be a ribose zipper and is therefore mediated almost entirely by backbone chemistry rather than by base identity. We have thus not been able to readily explore this further in detail by base substitution but propose it adds an additional stabilisation interface optionally used in nature in contribution to the earlier described interfaces.

Mechanistically domain II anchors and stabilise domain VI in the position necessary to catalyse forward splice 2 and/or reverse splice 1, as described previously (Chan et al., 2012; Marcia and Pyle, 2012; Robart et al., 2014). It appears that Voyager-1 has more extended interactions compared to LtrB. As we shall see below (state3, domain VI down particles) the domain II extension becomes disordered in some states and we propose probably serves as a steric gate becoming highly mobile. The extended form of domain II is clearly not obligatory for GIIA intron function but likely an adaptation to further stabilise domain VI-up intermediates in certain physiological environments.

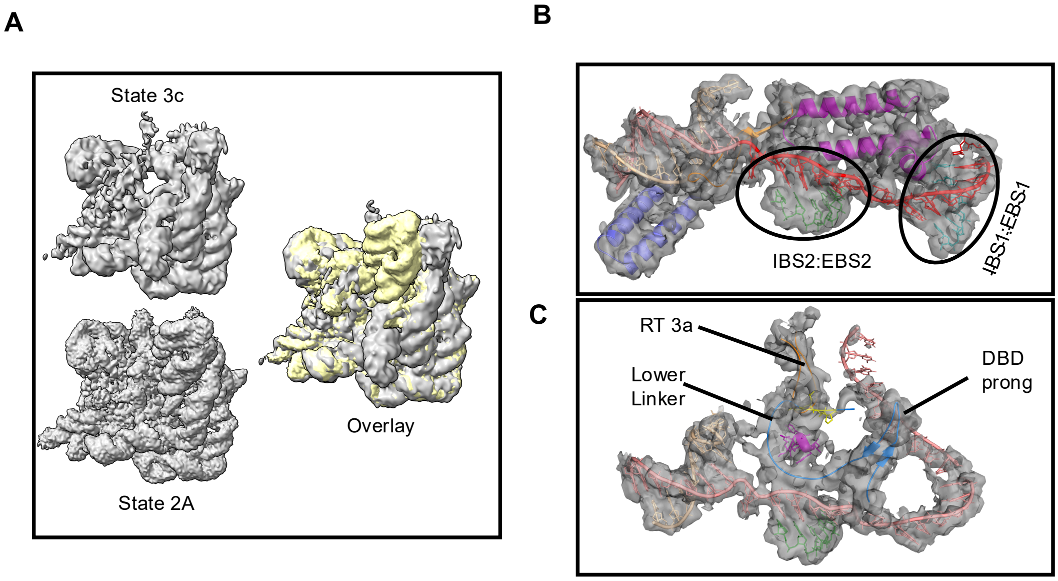


**Figure S5. Cryo-EM densities supporting a limited State 3c reconstruction.**

(A) Low-pass surfaces of the State 3c (upper) and State 2A (lower) maps in the same orientation, with the two superimposed at right (overlay; State 3c in yellow, State 2A in grey) to show that the two states share a common core and differ in the peripheral lobes.

(B) State 2 model mapped onto the electron density from state 3 for the two exon-binding duplexes. Circles mark the IBS2:EBS2 and IBS1:EBS1 pairs, with target DNA in red, intron RNA in pale colours and the IEP helices in magenta and blue. Overlap of density with the state 2 model suggests state 3c retains similar interactions.

(C) State 2 model mapped onto the electron density from state 3 in the region of the RT 3a insert, the lower linker and the DBD prong, each arrowed. Overlap of density with the state 2 model suggests state 3c retains similar interactions.

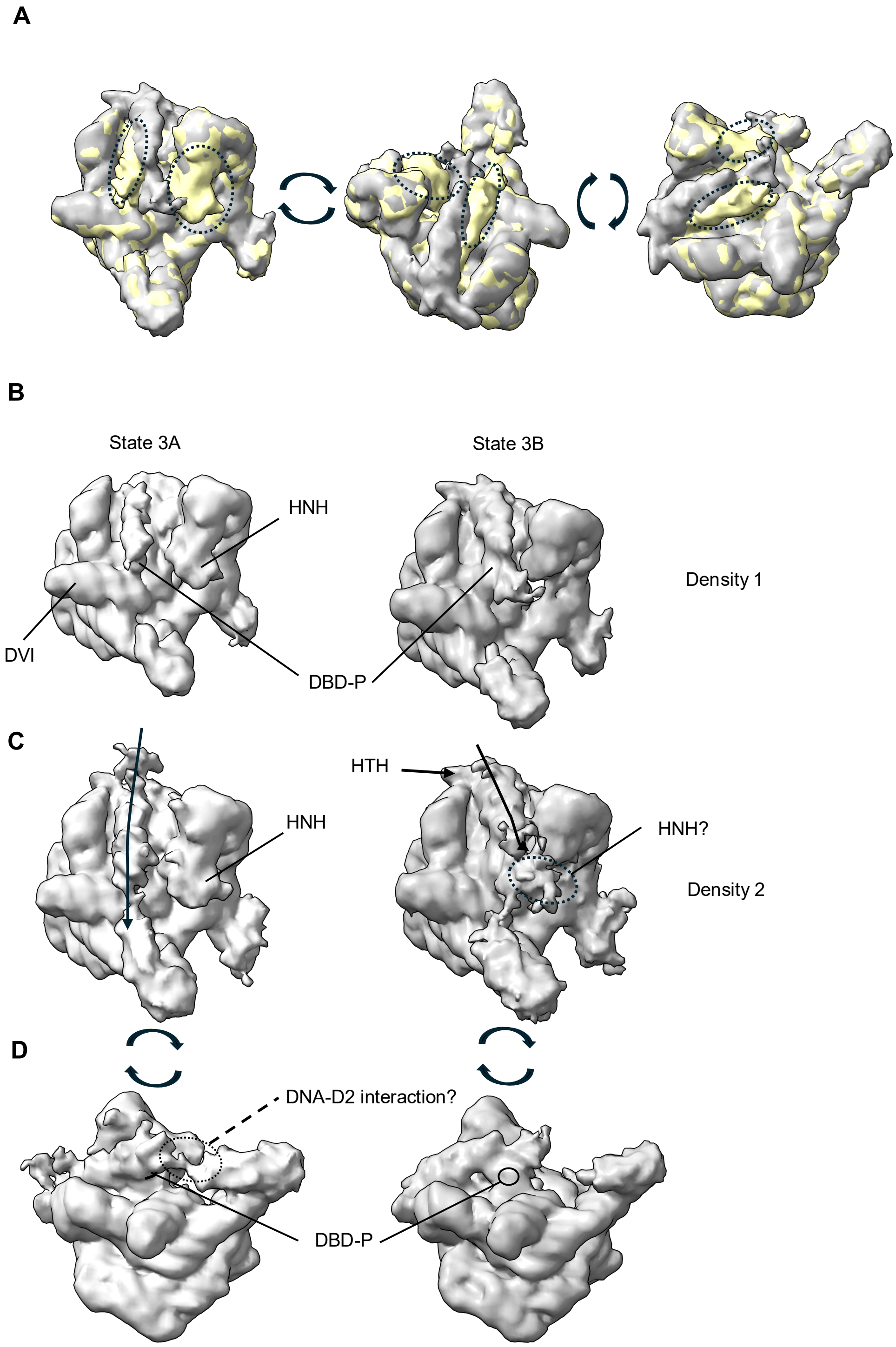


**Figure S6. Low resolution maps dubbed states 3A and 3B recovered from frustrated IBS1:EBS1 complexes differ in the position of the HNH domain and the position of suspected DNA strand.**

(A) Overlay of the State 3A and State 3B electron density maps (State 3A yellow, State 3B grey) in three orientations related by the rotations arrowed between them. Dotted outlines ring the two regions where the two maps disagree; the rest of the envelope is common to both.

(B) The first of those two regions, shown separately for State 3A (left) and State 3B (right) and labelled Density 1. In State 3A the lobe is resolved and can be assigned to the HNH domain, with domain VI (DVI) and the DBD prong (DBD-P) adjacent; in State 3B the same lobe is absent.

(C) The second region, labelled Density 2, in the same two states. The arrow follows the path from the HTH down into the body of the particle. In State 3B an unassigned density appears at the dotted outline in a position consistent with, but not sufficient to establish, a differently placed HNH domain — hence "HNH?".

(D) The same two maps rotated as arrowed, viewed onto the DBD prong. The dotted outline in State 3A encloses prong density that in State 3B is reduced to the small feature circled. The dashed leader marks density beside the prong in State 3A that may represent contact between the target DNA and the domain II extension; it carries a query because neither the assignment nor the contact is established at this resolution.

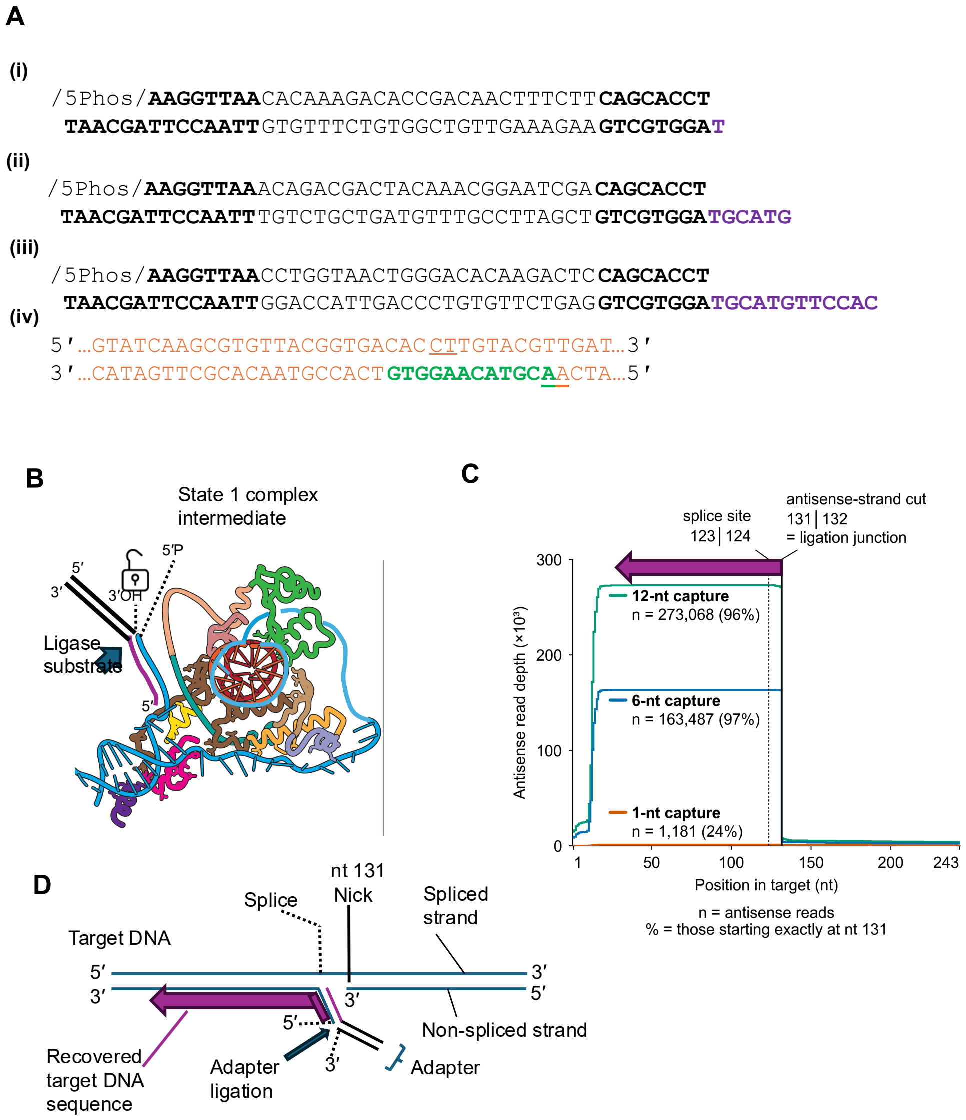


**Figure S7. Sequencing of the antisense-strand cut: adapter design, capture-overhang length and per-position read depth.**

(A) Sequences of the dsDNA sequencing adapters, each with its flanking segments and a capture overhang of (i) 1 nt, (ii) 6 nt or (iii) 12 nt (overhang in purple; the 5′-phosphate required for ligation is marked /5Phos/). (iv) The dsDNA retrohoming target, with triangles denoting the splice site (top strand) and the endonuclease nicking site (bottom strand); underlined bases flank the cleaved phosphates. The target is 243 bp long but only positions −25 to +12 relative to the splice site are shown. The preferred endonuclease nick site for Voyager-1 is +8/+9, which is 131/132 nt from the upstream end of the molecule.

(B) Schematic of the State 1 complex intermediate, showing the dsDNA adapter with its 12-nt capture overhang annealed to the 21-nt single-stranded region that reverse splicing generates in the absence of dNTPs, and the resulting 5′-phosphate/3′-OH pair presented to the ligase.

(C) Per-position read depth on the antisense strand of the 243-bp target after cleavage by the ribonucleoprotein and ligation of an adapter carrying a 1-nt, 6-nt or 12-nt capture overhang. The RNP cuts the antisense strand between nt 131 and nt 132 and leaves a 5′-phosphate that is ligated to the adapter, so nt 131 is both the ligation junction and the expected boundary of every productive read; a clean cut-and-ligate product carries no signal 3′ of it. The dotted line marks the splice site at 123|124. n is the number of antisense reads recovered and the percentage is the fraction of those reads starting exactly at nt 131: 1,181 (24%) for the 1-nt overhang, 163,487 (97%) for 6 nt and 273,068 (96%) for 12 nt. Lengthening the overhang beyond 6 nt therefore raises yield without improving precision.

(D) Where the recovered sequence sits on the target. The spliced and non-spliced strands are marked, together with the splice site and the nick at nt 131; the adapter ligates to the 5′-phosphate left at the nick and the purple arrow shows the stretch of target DNA read back from it.

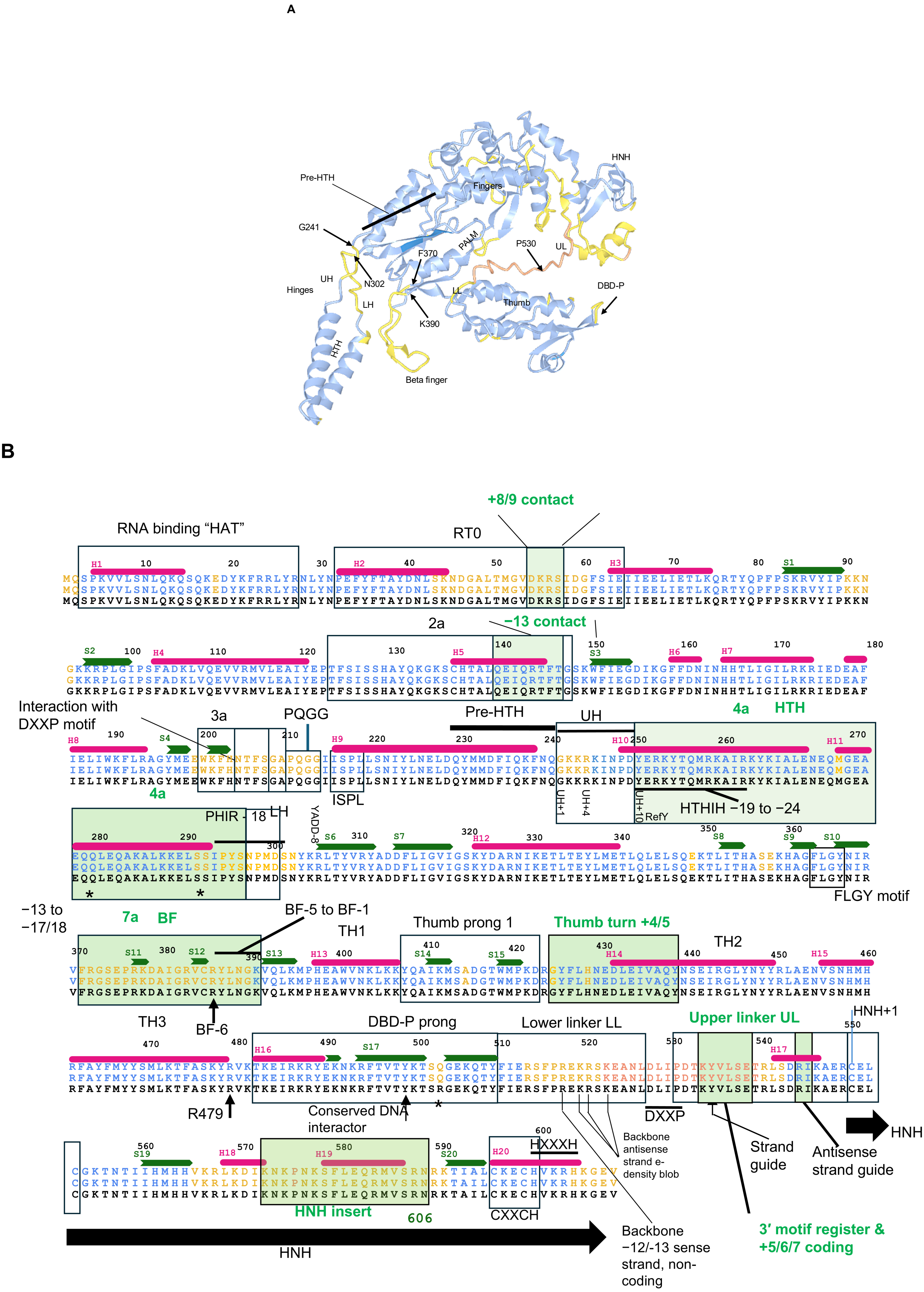


**Figure S8. Domain architecture of the Voyager-1 IEP and the sequence features that define it.**

(A) Ribbon model of the Voyager-1 IEP coloured by confidence of assignment — blue where the chain is modelled, yellow for the loops and inserts discussed in the text, salmon for the lower linker. Domains and subelements are labelled: the helix-turn-helix (HTH) with its upper and lower hinges (UH, LH) and the pre-HTH helix; the fingers and palm of the reverse-transcriptase core; the beta finger; the thumb; the lower and upper linkers (LL, UL); the DBD prong (DBD-P); and the HNH endonuclease domain. Boundary residues are arrowed: G241 and N302 at the hinges, F370 and K390 at the ends of the beta finger, and P530 in the upper linker.

(B) The Voyager-1 IEP sequence aligned to its closest database relative (UniProt A0A7Z8RW25_A; BLAST E = 0.0), with the consensus beneath in black. Residues are coloured by conservation. Secondary structure observed in the model is drawn above the sequence — magenta bars for helices H1–H20, green arrows for strands S1–S20 — and numbered. Boxes name the conserved blocks of the reverse-transcriptase and endonuclease families (RT0, 2a, 3a, PQGG, 4a, YADD, FLGY, 7a, DXXP, CXXCH, HXXXH) and the structural elements of (A). Green shading marks the segments that contact the DNA target, labelled with the target position each reads: the +8/9 contact in RT0, the −13 contact in 2a, HTHIH reading −19 to −24, the beta finger reading −13 to −17/18, the thumb turn at +4/5, and the upper linker carrying the 3′ motif register and the +5/6/7 code. Arrows and brackets below the sequence name individual residues of interest (R479, the conserved DNA interactor in the DBD prong) and the backbone contacts made to the sense and antisense strands. The black arrow at the foot marks the extent of the HNH domain, ending at residue 606.

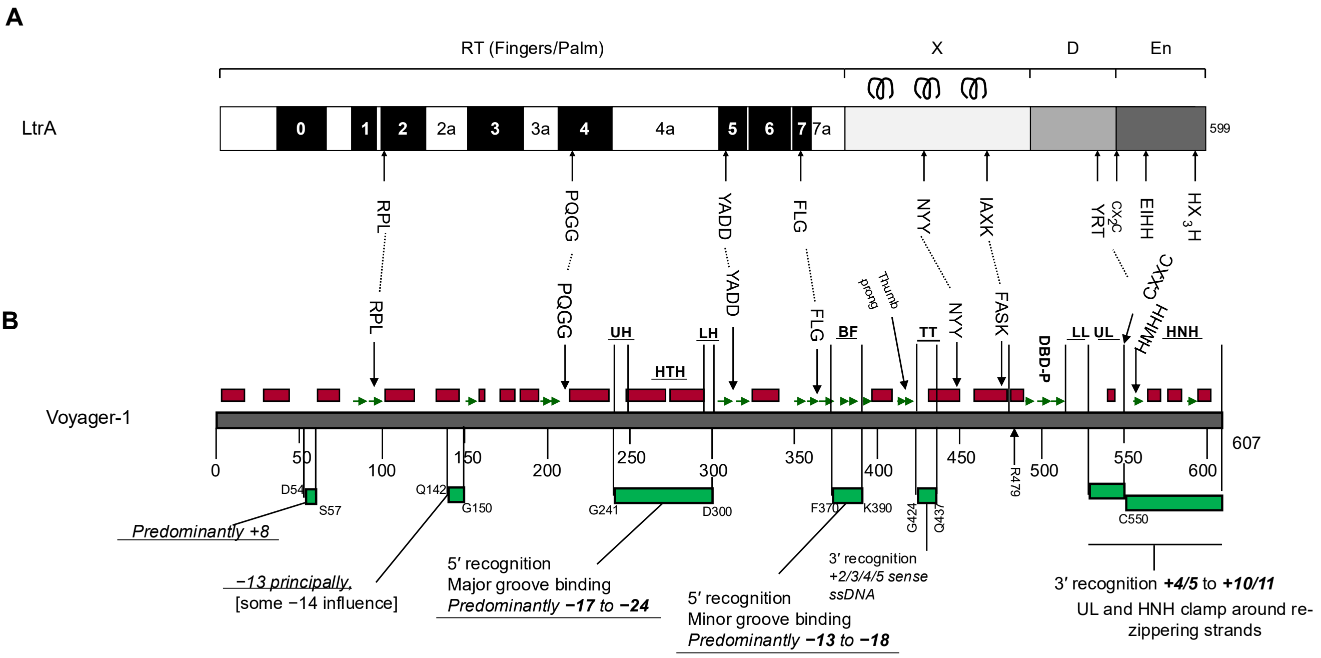


**Figure S9. The Voyager-1 IEP mapped onto the canonical group IIA intron-encoded protein.**

(A) The LtrA domain map drawn to scale: the reverse-transcriptase region (fingers and palm) carrying the conserved blocks 0–7 with the inserts 2a, 3a, 4a and 7a between them; domain X, whose three helices are drawn above it; the DNA-binding domain D; and the endonuclease domain En. The scale ends at residue 599. Arrows below name the sequence motifs used to anchor the alignment: RPL, PQGG, YADD, FLG, NYY, IAXK, YRT, $CX_2C$, EIHH and $HX_3H$.

(B) The Voyager-1 IEP on the same scale, numbered 0–607. Red blocks above the bar are the helices and green arrows the strands observed in the cryo-EM model; the labels above them mark the elements named in Figure S8 (UH, HTH, LH, FLG, BF, thumb prong, TT, NYY, FASK, DBD-P, LL, UL, HMHH, CXXC, HNH), and the arrows carry each LtrA motif down onto its Voyager-1 equivalent. Green blocks below the bar are the segments that read the DNA target, with their boundary residues given (D54–S57, Q142–G150, G241–D300, F370–K390, G424–Q437, R479, C550 to the C terminus). The annotations beneath state which part of the target each reads: the RT0 segment reads predominantly +8; the 2a segment reads principally −13 with some influence at −14; the HTH binds the major groove, predominantly −17 to −24; the beta finger binds the minor groove, predominantly −13 to −18; the thumb reads +2/3/4/5 of the sense strand as single-stranded DNA; and the upper linker and HNH domain together clamp the re-zippering strands over +4/5 to +10/11.

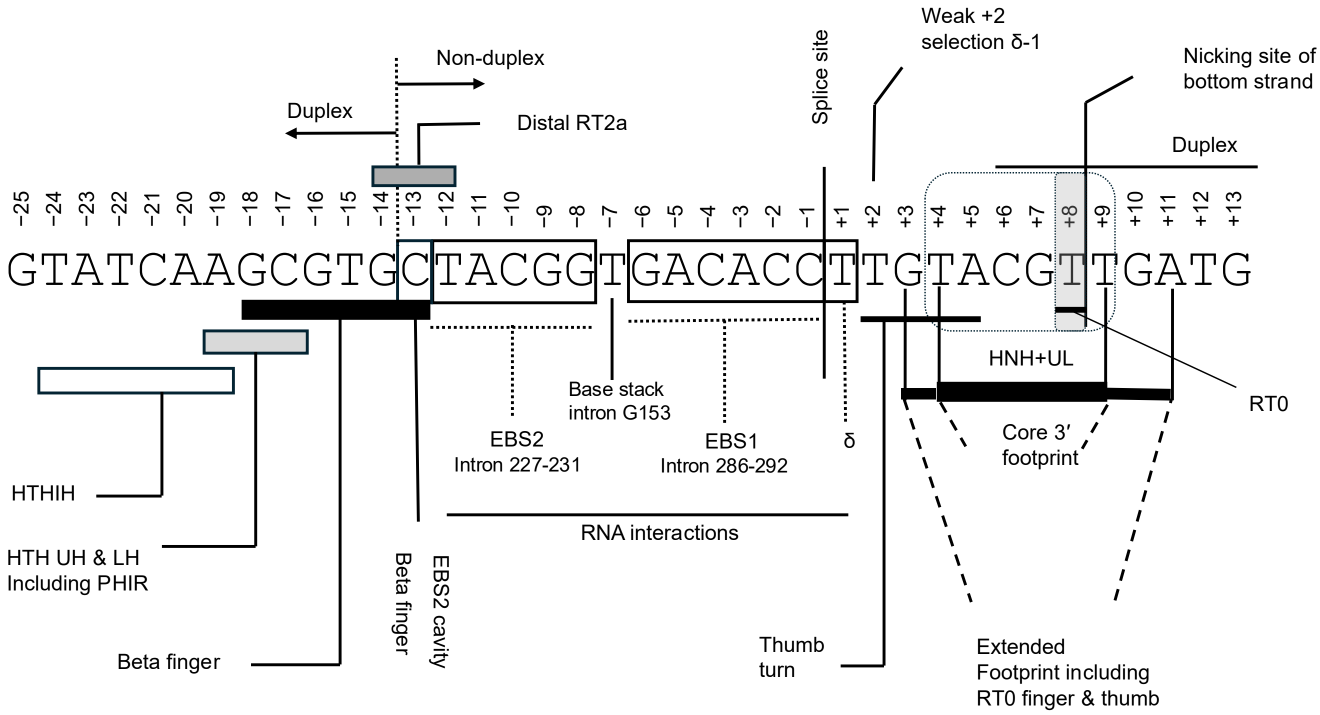


**Figure S10. Composite interaction map of the Voyager-1 target site.**

The target sense strand is written 5′ to 3′ and numbered −25 to +13 relative to the insertion site, with no position 0. Bars beneath the sequence give the footprint of each element: open bars for the HTH, grey bars for the HTH upper and lower hinges including PHIR, and solid bars for the beta finger and for the core 3′ footprint made by the HNH domain and upper linker. The grey block above the sequence is the distal RT2a contact at −13. Boxes on the sequence itself enclose the two RNA-pairing blocks, EBS2 (intron 227–231) over −12 to −8 and EBS1 (intron 286–292) over −6 to −1, with the δ pairing at −1 marked; the dotted leader between them marks the −7 position occupied by the base stack of intron G153. Vertical arrows and brackets mark the splice site between −1 and +1, the weak δ–1 selection at +2, the thumb turn at +3, and the nicking site on the bottom strand at +8/+9 (shaded). The horizontal arrows at the top give the state of the duplex: paired to the left of −14 and to the right of +9, unpaired between. Dashed lines at the right enclose the extended footprint that includes the RT0 finger and the thumb.

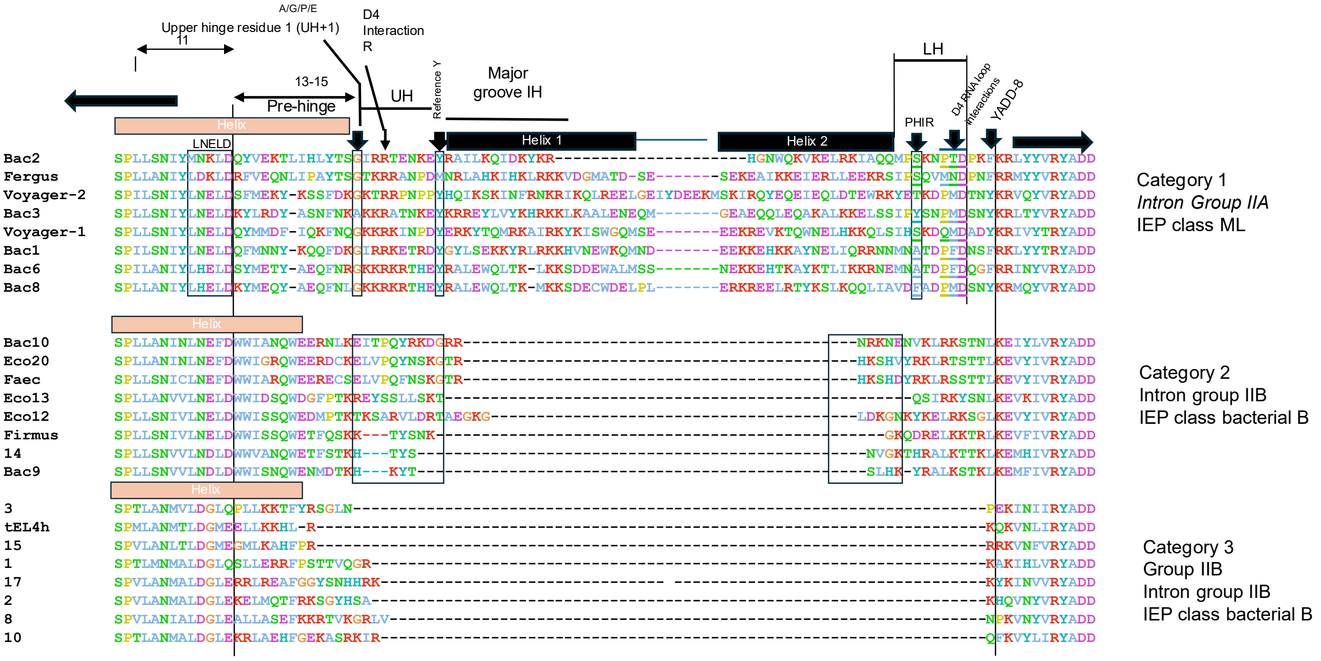


**Figure S11. The domain IVa insert separates class A intron-encoded proteins from class B.**

Alignment of the region spanning the pre-hinge, upper hinge, HTH and lower hinge for twenty-three intron-encoded proteins, residues coloured by chemistry and gaps shown as dashes. The proteins are grouped into three categories, named at the right: Category 1, group IIA introns with class ML intron-encoded proteins (Bac2, Fergus, Voyager-2, Bac3, Voyager-1, Bac1, Bac6, Bac8); Category 2, group IIB introns with class bacterial B proteins (Bac10, Eco20, Faec, Eco13, Eco12, Firmus, 14, Bac9); and Category 3, a second set of group IIB introns with class bacterial B proteins (3, tEL4h, 15, 1, 17, 2, 8, 10). Bars above the alignment mark the observed secondary structure — the salmon helix of the pre-hinge, and the two black helices of the major-groove-binding HTH (Helix 1 and Helix 2). Vertical arrows and boxes name the positions used to define the register: the first residue of the upper hinge (UH+1, which is A, G, P or E), the arginine that interacts with domain IV, the reference tyrosine, the PHIR position in the lower hinge, the residues that contact the domain IV RNA loop, and the YADD motif that closes the window at the right. Horizontal measures at the top left give the spacing between these anchors (11 residues; 13–15 residues). Only Category 1 carries the extended insert that supplies Helix 2 and the major-groove contact; in Categories 2 and 3 the corresponding segment is deleted, so the HTH of a class A protein has no equivalent in a class B protein.

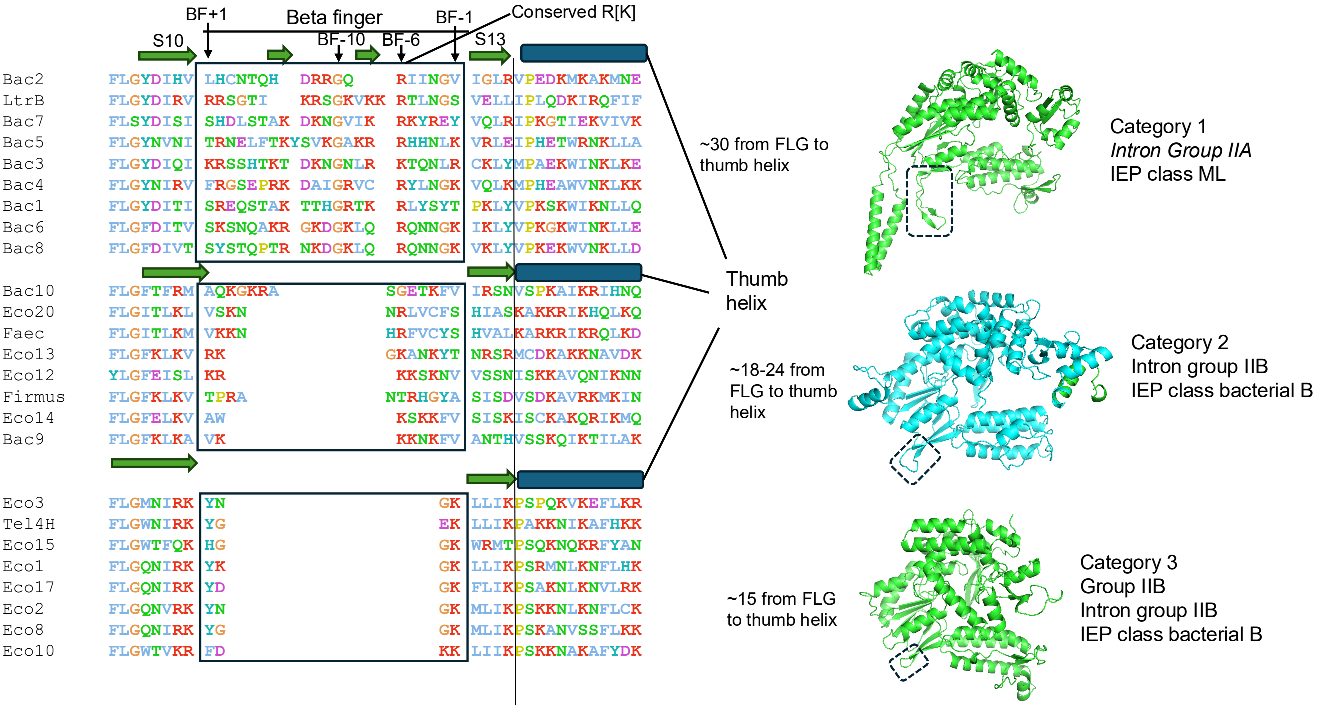


**Figure S12. The beta finger and thumb helix across group II intron-encoded proteins.** Sequences from class IIA and class IIB intron-encoded proteins (IEPs) are aligned on the FLG motif at the left and on the conserved R[K] and the thumb helix at the right, with the S10 and S13 strands arrowed and the beta-finger limits marked BF+1, BF−10, BF−6 and BF−1. Residues are coloured by identity and dashes are alignment gaps. Voyager-1 appears under its laboratory name Bac4, and other Gene Weaver isolates under their Bac, Eco, Fergus, Xeno and Tel4h designations; remaining rows are database entries, given either by assembly accession or by the internal clade identifier under which they were recovered. Category 1 is group IIA with the maturase-like (ML) IEP class and categories 2 and 3 are group IIB with the bacterial class B IEP. The three categories differ in the length of the finger: approximately 30 residues separate FLG from the thumb helix in category 1, 18 to 24 in category 2 and 15 in category 3, so the elaborate class IIA finger is progressively reduced in the class IIB proteins. Ribbon models at the right show the corresponding region, boxed, in a representative of each category.

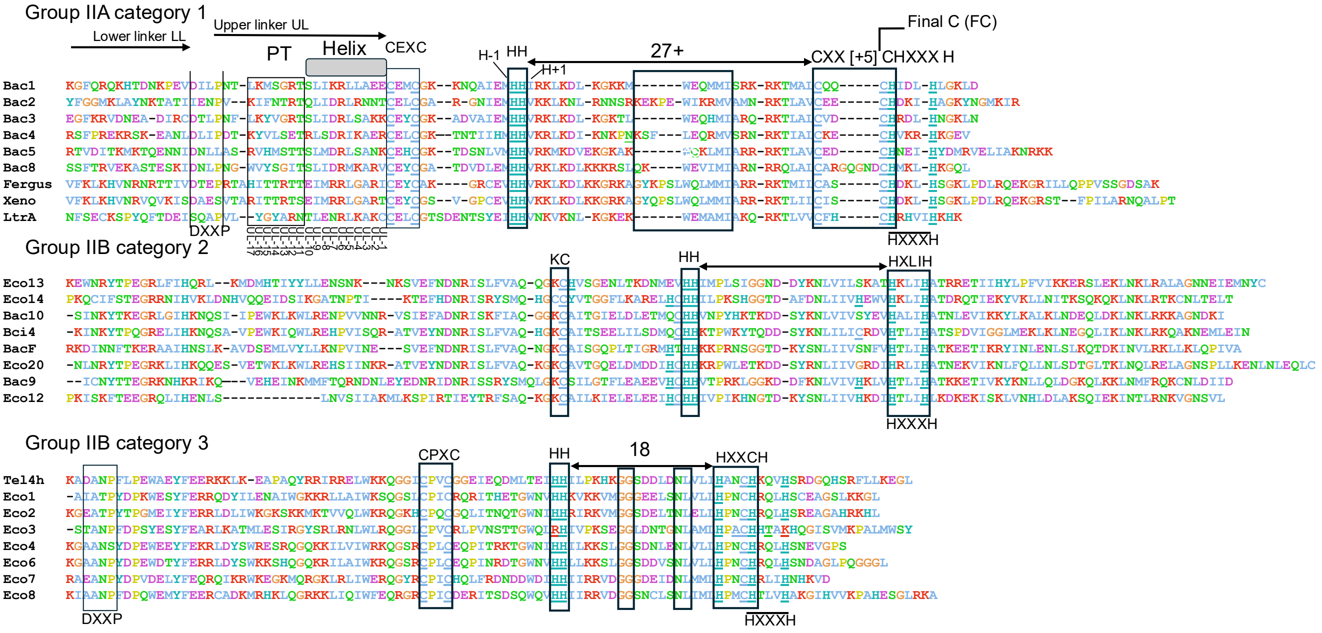


**Figure S13. The HNH domain and its linkers across group II intron-encoded proteins.** Group IIA category 1 above, then the two group IIB categories. Residues are coloured by identity and dashes are alignment gaps. Voyager-1 appears under its laboratory name Bac4, and other Gene Weaver isolates under their Bac, Eco, Fergus, Xeno and Tel4h designations; remaining rows are database entries, given either by assembly accession or by the internal clade identifier under which they were recovered. Category 1 is group IIA with the maturase-like (ML) IEP class and categories 2 and 3 are group IIB with the bacterial class B IEP. Annotations mark the lower and upper linkers, the pass-through segment (PT) and the helix within it, the CEXC motif that opens the domain, the H−1, HH and H+1 positions, and the cysteine and histidine positions boxed at the right. The spacing between the HH pair and the terminal ligands separates the categories: 27 or more residues in group IIA category 1, which ends in a CXX[+5]CHXXXH arrangement and a final cysteine (FC), against the HXLIH of category 2 and the 18-residue spacing and HXXCH of category 3. The DXXP motif at the start of the upper linker, and the CPXC and KC positions, are marked where present.

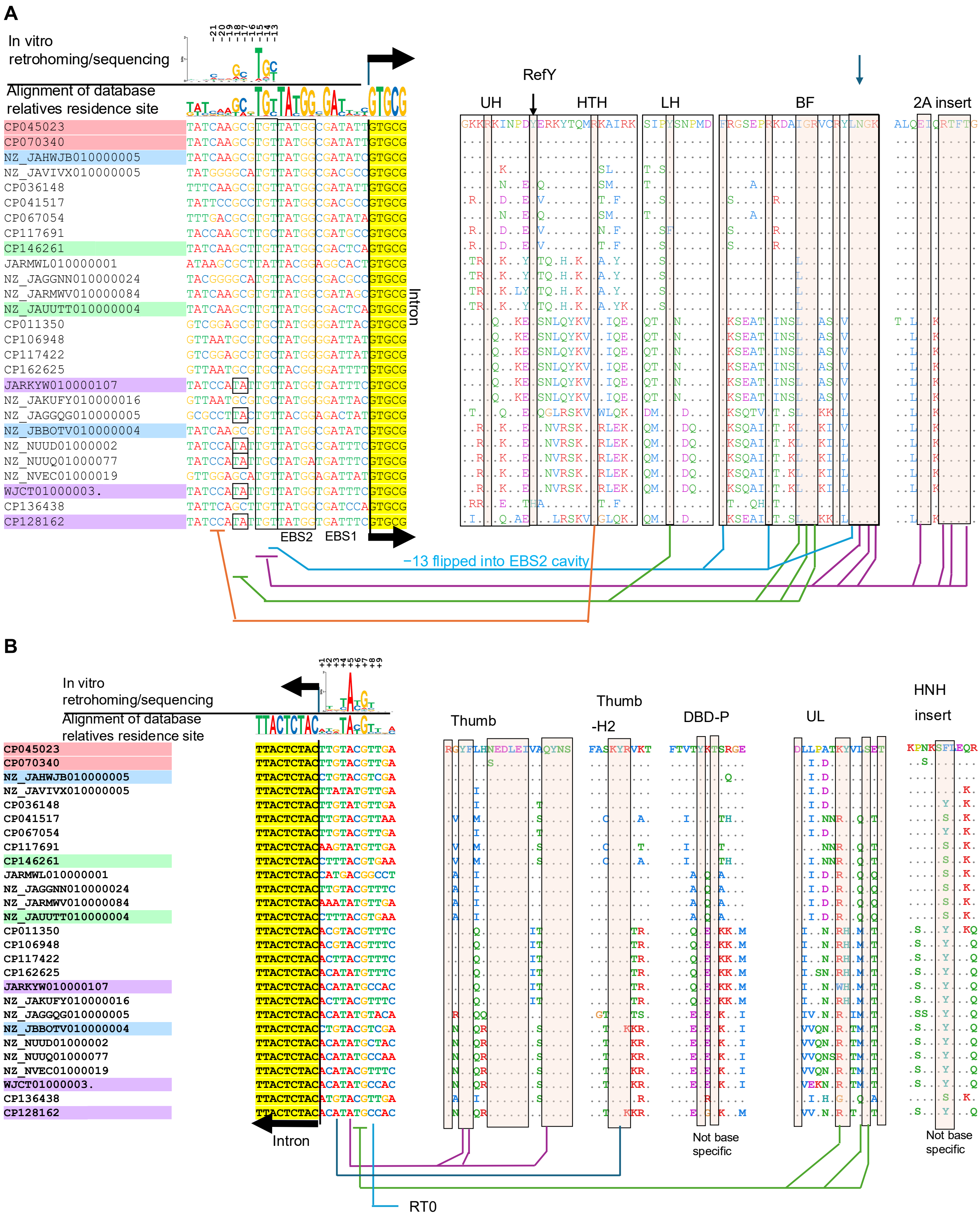


**Figure S14. Coding contacts in Bac4 relatives: conserved residues in contact regions suggest those most critical for coding.**

Both panels show the same twenty-seven database relatives of Bac4, listed by accession at the left, with coloured highlighting grouping the accessions that share a residence-site sequence. At the left of each panel the residence site recovered for each relative is aligned, above it the sequence logo of those aligned residence sites, and above that the sequence logo obtained from in vitro retrohoming and sequencing of Voyager-1, and the black arrow marks the intron and the direction of reading. On the right protein segments of interest that approach that part of the target are aligned, the top row being Bac4 as reference, a dot meaning identical to the reference and a coloured letter a residue that differs. Coloured lines running beneath the figure connect each protein column to the target position it reads. Both logos give information content in bits on a 0 to 2 scale, with bases coloured A red, C blue, G amber and T green.

(A) The 5′ side. Target positions covering EBS2 and EBS1 are aligned at the left, and the protein segments at the right are the upper hinge (UH), the HTH, the lower hinge (LH), the beta finger (BF) and the 2A insert, with the reference tyrosine (RefY) arrowed. Shaded columns are conserved regions of interest, specifically those that likely contact or approach DNA, or instead may mediate other key interactions driving conservation. The blue annotation marks the −13 base that flips into the EBS2 cavity. Relatives that share a residence-site sequence also share the residues in the shaded columns, and the three sequence groups visible in the target alignment are recovered independently in the protein alignment.

(B) The 3′ side, same twenty-seven relatives and same conventions. The protein segments shown are the thumb, thumb helix 2, the DBD prong, the upper linker (UL) and the HNH insert, with the RT0 contact marked below. The DBD prong and the HNH insert are annotated “not base specific”: they vary between relatives without closely tracking the target sequence, so their variation is not read as coding.

Note that the interactions proposed are not exhaustive but the conservation/interaction map intended only to aid and support other experimental and structural data.

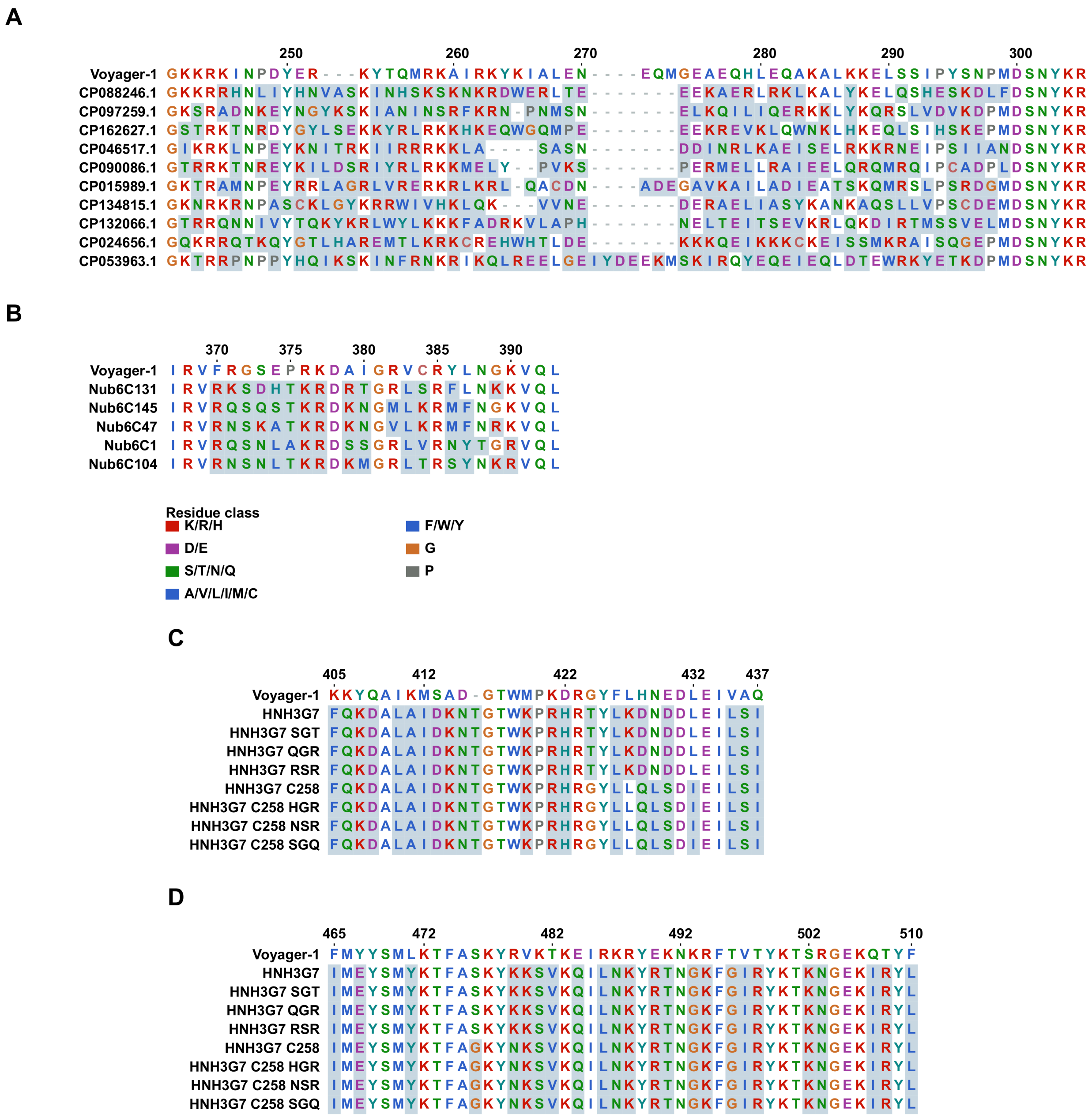
A
Voyager-1
CP088246.1
CP097259.1
CP162627.1
CP046517.1
CP090086.1
CP015989.1
CP134815.1
CP132066.1
CP024656.1
CP053963.1
B
Voyager-1
Nub6C131
Nub6C145
Nub6C47
Nub6C1
Nub6C104
Residue class
K/R/H
D/E
S/T/N/Q
A/V/L/I/M/C
F/W/Y
G
P
C
Voyager-1
HNH3G7
HNH3G7 SGT
HNH3G7 QGR
HNH3G7 RSR
HNH3G7 C258
HNH3G7 C258 HGR
HNH3G7 C258 NSR
HNH3G7 C258 SGQ
D
Voyager-1
HNH3G7
HNH3G7 SGT
HNH3G7 QGR
HNH3G7 RSR
HNH3G7 C258
HNH3G7 C258 HGR
HNH3G7 C258 NSR
HNH3G7 C258 SGQ

E

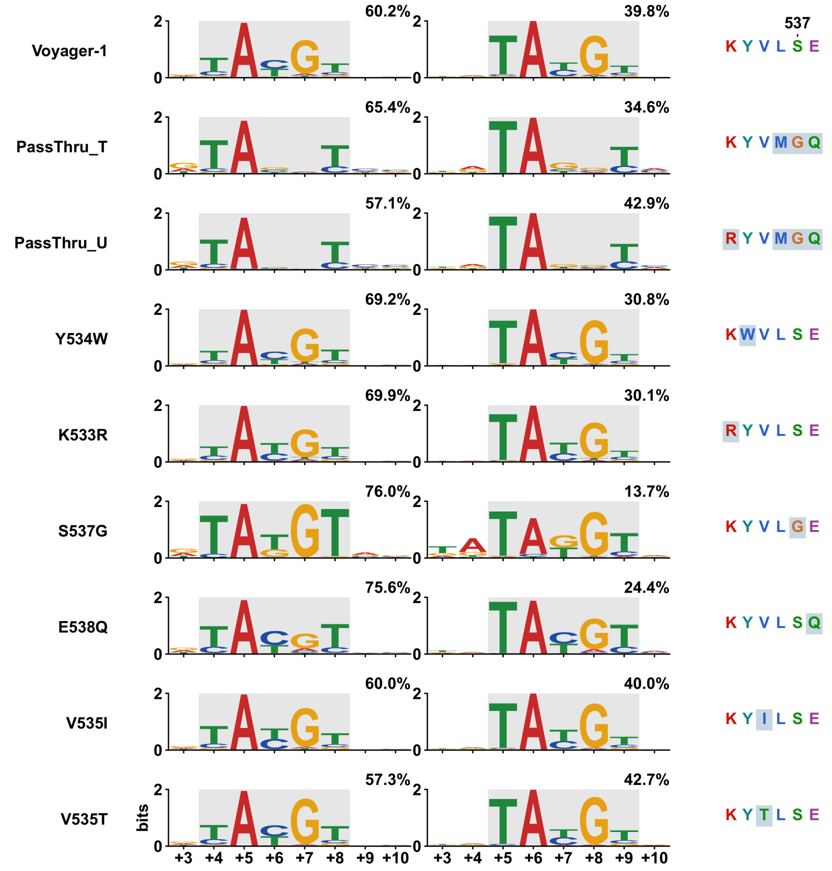


F

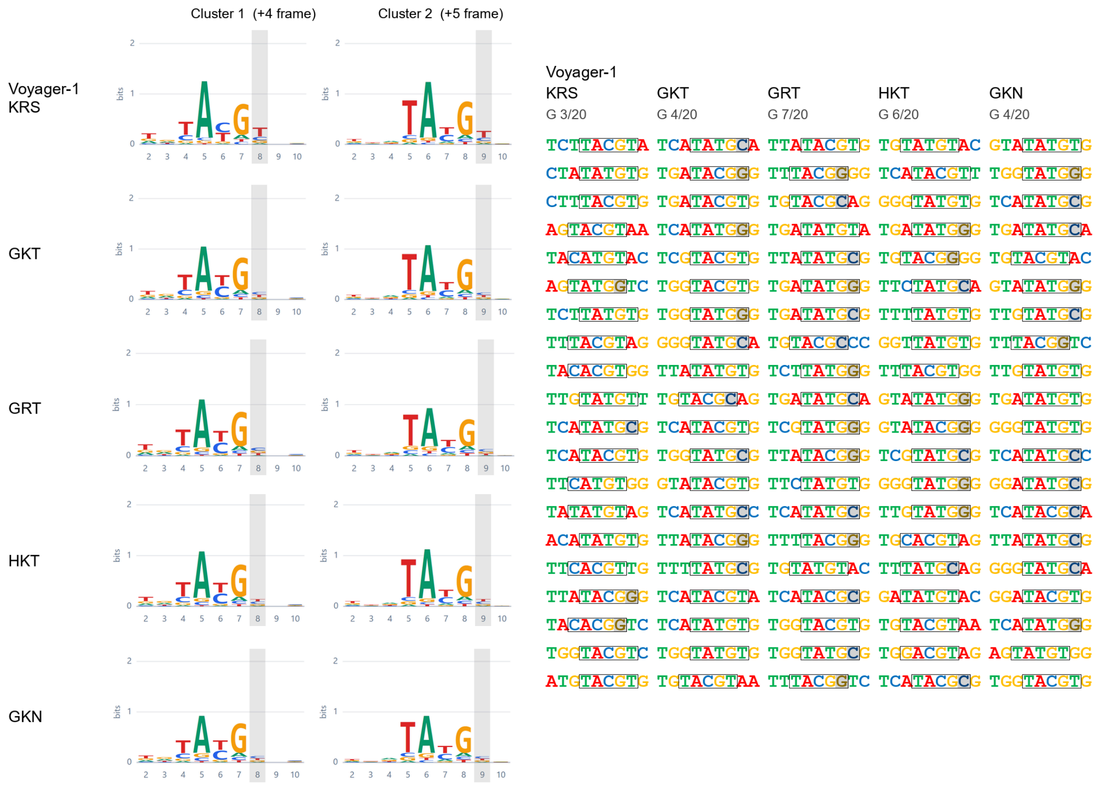


**Figure S15. Sequence context for the reprogramming constructs of Figures 5 and 6.**

(A) The hinge–HTH region. Voyager-1 G241–R305 aligned across the ten natural class IIA donors whose blocks were grafted to give the hybrids in Figure 5D, each labelled by accession.

(B) The beta-finger variable region. Voyager-1 I367–L393 with the five Nub6C donors used in Figure 5E.

(C and D) The two segments in which the Figure 6 constructs differ from Voyager-1: the thumb at 405–437 (C) and the thumb helix 2 and pre-prong region at 465–510 (D). Eight of the eighteen constructs differ in each window; the remainder match Voyager-1 across both. Also varying, too short to draw as a block: 291 (17 constructs, all S291G).

(E) Upper-linker substitutions, with fitted motifs over +3 to +10 and the pass-through linker sequence K533–E538 written at the right of each row. Letter heights are information content in bits and the percentages are fitted mixture weights for the two components drawn; grey marks the motif core supported by the register-shift test. Each row is one construct: its two drawn components, then the linker residues behind them. A construct's two components share a labelled scale whose limit varies between constructs. Products are those significant at $P < 0.001$ under the site-dependent Poisson model, each against the matched input library of its own batch; the number of components is the paper's pairwise selection, the paired bootstrap vote included. The grey core is tested at $P < 0.05$ against 4,000 matched decoys. The two drawn components of every construct, Voyager-1 included, lie one base apart, so the grey steps right between the columns. S537G fits a further component carrying 10.2% of its products, at 1.5 bits over this window against 8.5 and 7.6 for the two drawn, and no single position above 0.5 bits. A component that flat is the products the specific components do not claim rather than a register of its own, so it is named here rather than drawn; the matrix holds back any component below 3 bits. The rows are not one assay: they come from 25/2, 25/4, 31/14, 31/15, 31/16, with Voyager-1 drawn from 31/16. Rows are cut to the window those assays share, and each was measured against its own batch's parent. Also varying, too short to draw as a block: 291 (8 constructs, all S291G).

(F) RT0 substitutions at the +8/+9 contact. Integration-sequencing logos, two clusters per construct, and the twenty most frequently recovered target sequences for Voyager-1 (K55, R56, S57; KRS) and for four substitutions of that triplet; D54 was not altered. Boxes mark the recovered YAYGY motif in each sequence and follow its register, which is distributed between the +4 and +5 frames. At the final motif position G rises from 3 of 20 sequences in Voyager-1 to between 4 and 7 of 20 in the substitutions, and A is absent from all five constructs. Bits axes differ between constructs, so letter heights are not comparable between rows. The grey band marks that final position in each logo, +8 in the Cluster 1 (+4 frame) logos and +9 in Cluster 2 (+5 frame); in the sequence lists a base other than T there is set on a grey ground in a darker tone, and the count of G at that position is given above each list. Information content at this column falls as the preference relaxes, so the shift toward G reads more clearly in the sequence lists than in the logo letter heights.

Throughout (A–E), colour denotes residue class, tint marks a difference from Voyager-1 and a dash marks a gap. All positions are Voyager-1 (Bac4) residue numbers and exclude the purification tag. Every reference column within a displayed block is kept; an insertion column no construct in the panel fills is dropped.

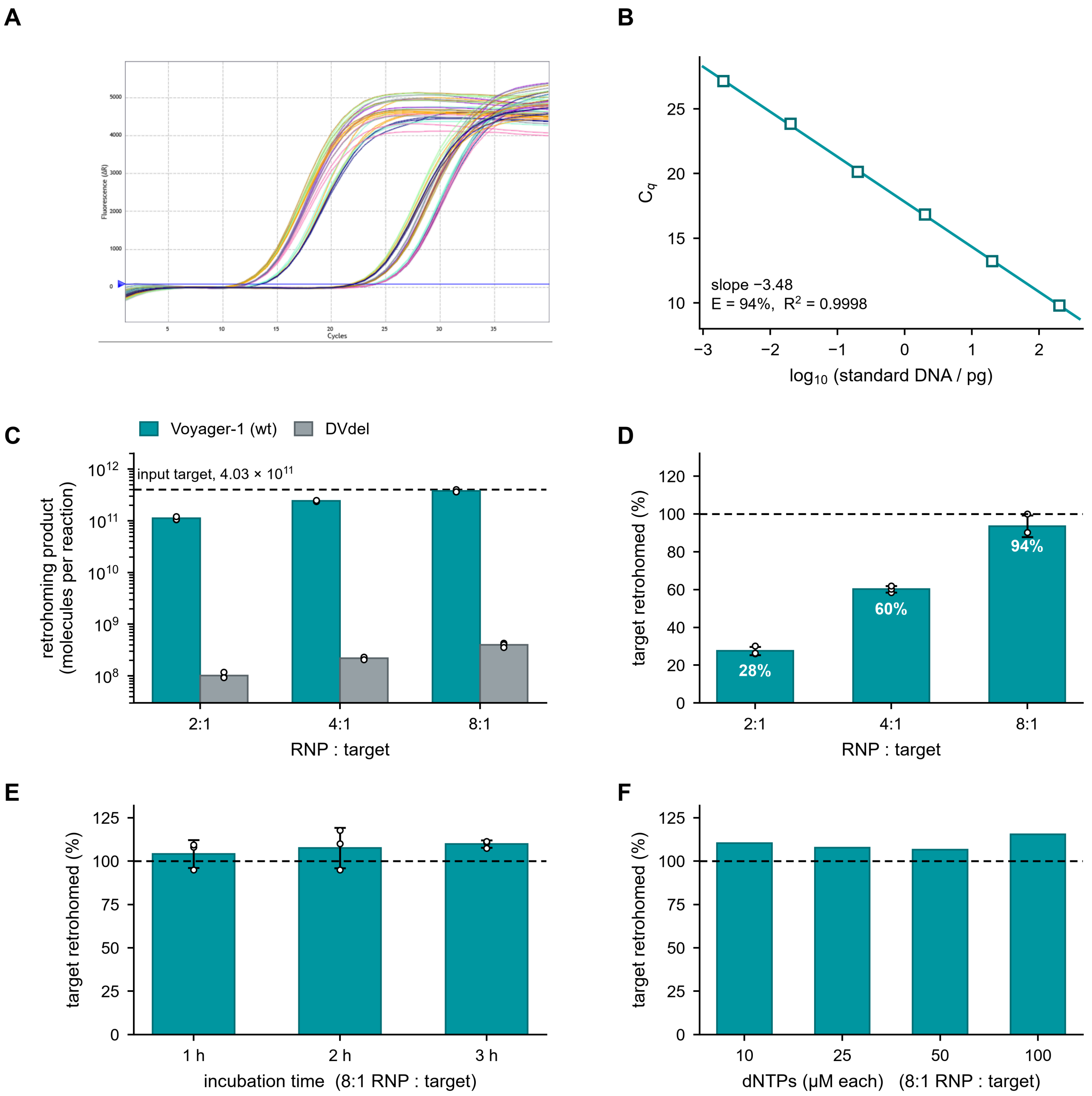


**G**

Conversion of $C_q$ to molecules of retrohoming product

1. $C_q$ → pg of standard-equivalent DNA, from the curve in (B).
2. pg → molecules, using $M_r = 6.88 \times 10^5$ g $mol^{-1}$ for the 1,114-bp double-stranded standard.
3. × 100 × 40 μL, to return the 1 μL of a 1:100 dilution assayed to the whole 40 μL reaction.
4. × 2, the single-strand correction (see legend): the standard is duplex and doubles from the first PCR cycle, whereas across the junction the retrohoming product carries only one DNA strand, its complement still being RNA. The first cycle therefore copies it without doubling it, so the product trails an equimolar duplex by one duplication — 1.05 $C_q$ at the measured E of 94%.
5. ÷ 4.03 × $10^{11}$ molecules of input target (0.67 pmol) to express as a percentage.

**Figure S16. Voyager-1 saturates its DNA target: quantification of retrohoming by junction-specific qPCR.** Retrohoming reactions contained 0.67 pmol (4.03 × $10^{11}$ molecules) of the 243-bp double-stranded DNA target together with Voyager-1 intron RNA and intron-encoded protein (IEP) at 1.34, 2.68 or 5.34 pmol, giving ribonucleoprotein (RNP)-to-target ratios of 2:1, 4:1 and 8:1, in a final volume of 40 μL at 37 °C. Product was measured with the junction-specific hydrolysis-probe assay of Figure S3E, in which a forward primer in the flanking target DNA and a reverse primer within the intron cDNA bracket the insertion junction. One microlitre of a 1:100 dilution of each reaction was assayed in 20 μL, in three technical replicates, against a standard curve of a synthetic 1,114-bp double-stranded DNA fragment carrying the same junction. DVdel is the domain V deletion RNP, which cannot splice and serves as the negative control. All quantities are corrected for the single-stranded character of the retrohoming product, as set out in (G) and below.

(A) Amplification curves. Fluorescence (ΔR) against cycle number for all reactions and standards on the plate.

(B) Standard curve. $C_q$ against $\log_{10}$ of the mass of double-stranded standard DNA per reaction, over six ten-fold steps from 200 pg to 0.002 pg. The fitted slope of −3.48 corresponds to an amplification efficiency of 94% ($R^2$ = 0.9998); sample $C_q$ values fall within the range of the curve.

(C) Retrohoming product recovered. Molecules of junction product per 40 µL reaction at each RNP-to-target ratio, on a logarithmic scale, for wild-type Voyager-1 and for DVdel. Bars are the mean of three technical replicates and open circles the individual replicates. The dashed line marks the $4.03 \times 10^{11}$ molecules of target supplied. Wild-type product exceeds the DVdel background by approximately three orders of magnitude at every ratio (950- to 1,110-fold).

(D) Proportion of the target retrohomed. The wild-type data of (C) expressed as a percentage of the input target: 28%, 60% and 94% at 2:1, 4:1 and 8:1. Error bars are the standard deviation of three technical replicates and the dashed line marks the input target. Product scales with the RNP supplied until, at 8:1, essentially all of the available target has been converted.

(E) Time course. Target retrohomed after 1, 2 and 3 h at 37 °C at an 8:1 RNP-to-target ratio. The target is already saturated at the first time point, so the reaction reaches completion within 1 h and nothing further is gained by extending it.

(F) dNTP titration. Target retrohomed at an 8:1 RNP-to-target ratio with each dNTP supplied at 10, 25, 50 or 100 µM for 1 h. The target is saturated throughout, so reverse transcription is not limited by dNTP availability anywhere in this range; 10 µM of each dNTP is already sufficient.

(G) Derivation. The chain of factors converting a $C_q$ value into molecules of retrohoming product, and thence into a percentage of the input target.

*The single-strand correction.* The standard is supplied as double-stranded DNA, so both of its strands template amplification from the first PCR cycle and it enters geometric amplification immediately. The retrohoming product does not. Across the region the assay interrogates, reverse transcription has generated only one DNA strand; the complementary strand is still intron RNA over most of that span. The first cycle therefore converts the single DNA strand into a duplex but does not duplicate it, and the product enters geometric amplification one duplication behind an equimolar duplex. Quantities read directly off the standard curve consequently understate the true number of product molecules by a factor of two, and every value reported here has been multiplied by two accordingly. The deficit is a factor of two in amount irrespective of the amplification efficiency, because the event lost is one genuine duplication of the template; expressed as a threshold shift at the measured efficiency of 94%, it corresponds to 1.05 cycles of $C_q$. The correction is an upper bound, in that any product whose RNA strand had been displaced or degraded before the assay would already be duplex and would need no correction.

*Saturation, and values above 100%.* In (E) and (F) the quantification returns 104–115% of the input target. No reaction can convert more target than was supplied, and these values are not evidence that it did: they are the saturated condition reported through a chain of measurements that together carry some 10–15% systematic tolerance, contributed by the molecular weight assumed for the standard, the dilution and pipetting steps, and the upper-bound character of the single-strand correction above. Read simply, the experiment shows that at an 8:1 RNP-to-target ratio the target is saturated — which is the ratio used for the time course in (E) and the dNTP titration in (F), and the reason neither shows a trend. The titration in (D) is the panel that carries the quantitative claim, and its 94% at 8:1 should be read with the same tolerance as approaching complete conversion rather than as a precise figure.

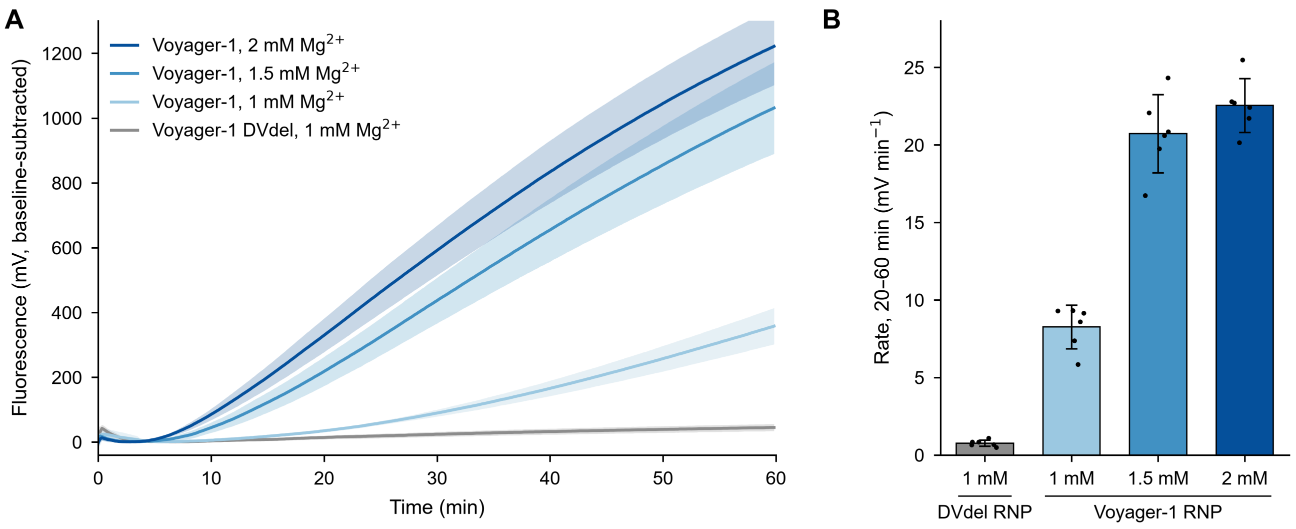


**Figure S17. Voyager-1 retrohomes in vitro at low millimolar magnesium.** Retrohoming was followed with the fluorophore/quencher double-stranded DNA assay of Figure S3C, in which reverse transcription across the insertion junction displaces the quencher-bearing strand and relieves quenching. Voyager-1 ribonucleoprotein (RNP) particles were reacted with the labelled double-stranded DNA target at 37 °C in 1, 1.5 or 2 mM $Mg^{2+}$; the domain V deletion RNP (DVdel), which cannot splice, was assayed in parallel at 1 mM $Mg^{2+}$ as the negative control. Three independent runs were performed, each with duplicate tubes per condition, and all six tubes per condition are pooled throughout.

(A) Fluorescence time courses. Line, mean of the six tubes; shading, standard deviation. Each tube is baseline-subtracted against its own minimum between 1 and 10 min. Signal rises above the DVdel control at every magnesium concentration tested, 1 mM included; at 60 min the signal is 358 ± 56, 1,031 ± 142 and 1,222 ± 119 mV at 1, 1.5 and 2 mM $Mg^{2+}$, against 45 ± 11 mV for DVdel.

(B) Initial rates. Slope of a linear fit over 20–60 min. Bars, mean of the six tubes; error bars, standard deviation; dots, individual tubes. Rates are 0.77 ± 0.20 mV $min^{-1}$ for DVdel and 8.3 ± 1.4, 20.7 ± 2.5 and 22.6 ± 1.7 mV $min^{-1}$ at 1, 1.5 and 2 mM $Mg^{2+}$, that is 11-, 27- and 29-fold over the control. The reaction is close to saturation by 1.5 mM, at 92% of the 2 mM rate, and retains 37% of that rate at 1 mM.

Retrohoming by the holoenzyme therefore does not require the high magnesium of protein-free ribozyme assays. Concentrations are total Mg2+ added; the 25 µM of each dNTP present chelates of the order of 0.1 mM, so free $Mg^{2+}$ is a little below the nominal value. Activity at 1 mM falls within the 0.5–1.2 mM range reported for free $Mg^{2+}$ in mammalian cells, indicating that the magnesium requirement of retrohoming need not be high.

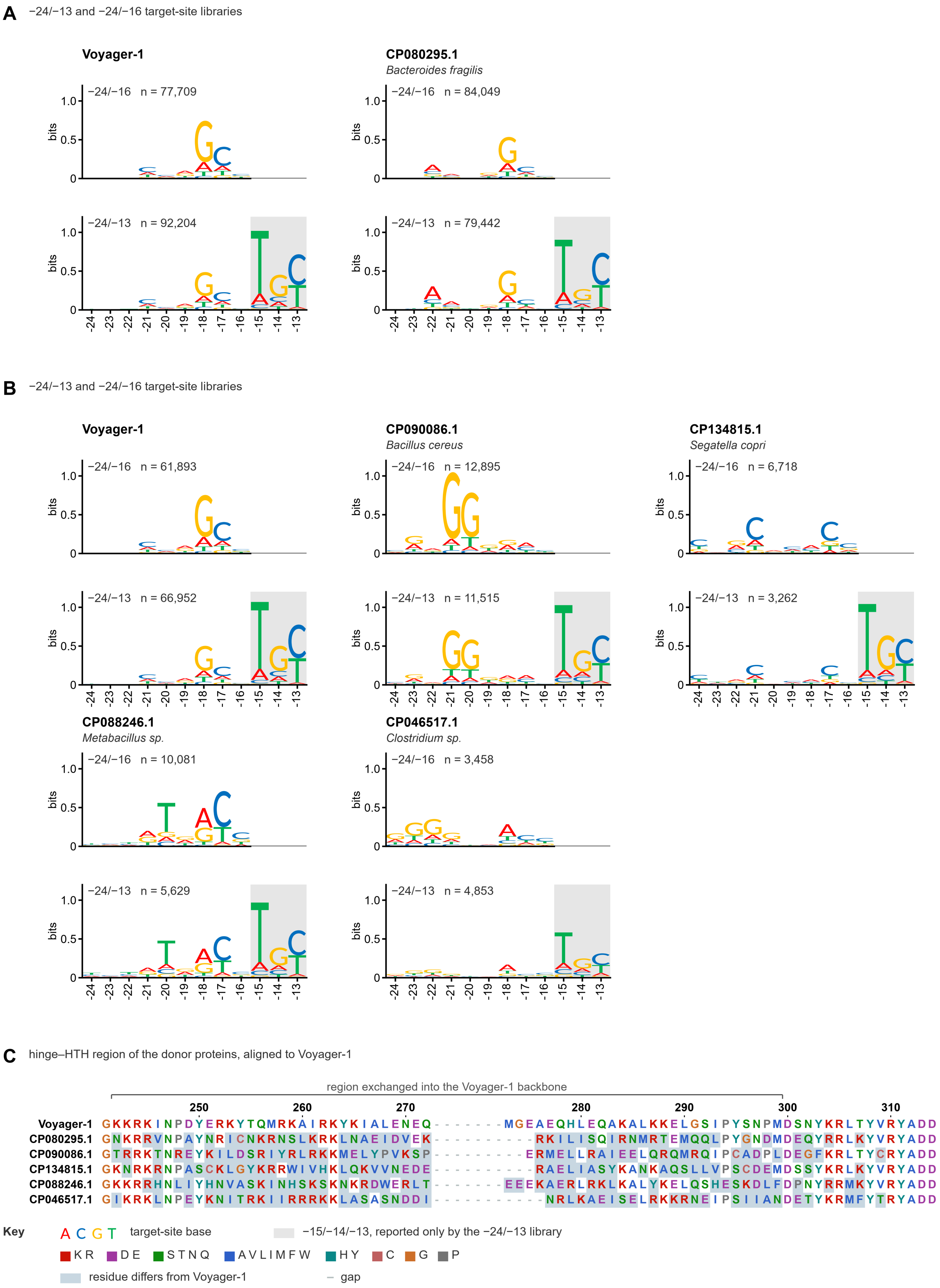


**Figure S18. Exchanging the hinge–HTH region changes recognition of −24 to −16 but leaves −15/−14/−13 unaltered.** Voyager-1 derivatives in which the hinge–HTH region has been replaced by the corresponding region of the intron-encoded protein from the genome indicated. Constructs are named by the donor's genome accession. For each, the target-site preference measured on the restricted −24/−16 library is shown above the preference measured on the full −24/−13 library; both traces share one position pitch, so a given position lies in the same column throughout the figure. Sequence logos give information content in bits ($R_i = 2 - H_i$, no small-sample correction); n is the number of distinct target sequences recovered. **The bit axis runs 0 to 1.2, not the conventional 0 to 2**: the largest stack in the figure is 1.07 bits, and the shorter axis is used throughout so differences between constructs are visible. The scale is identical in every trace. Recovery depth differs between libraries and batches, so each panel carries its own Voyager-1 reference and comparisons are made within a panel.

**(A) Voyager-1 and the *Bacteroides fragilis* CP080295.1 donor.** The G at −18 and C at −17 selected by Voyager-1 are weakened, a preference appears at −22, and −15/−14/−13 reads T·G·C in both proteins.

**(B) Voyager-1 and four further donors.** CP090086.1 selects G at −21 and −20 in both libraries (1.05 and 0.79 bits at −24/−16; 0.70 and 0.59 at −24/−13; consensus CGAGGG over −24 to −19 in each). CP088246.1 reproduces a three-position signature in both: T at −20 (0.56 and 0.48 bits), A at −18 (0.49 and 0.37) and C at −17 (0.71 and 0.54). CP046517.1 gives consensus GGGGTAACC in both, with G at −23 and −22 resolved in the restricted library. CP134815.1 shifts selection to C at −21 and −17; its −24/−16 measurement comes from a separate run of the restricted library, so its two traces are not matched for depth. In every case the lower trace reads −15/−14/−13 as T·G·C.

**(C) The exchanged regions.** The hinge–HTH region of each donor aligned to Voyager-1 from G241 to the RYADD motif and numbered by Voyager-1 residue. Rows are registered on the initial G, the reference tyrosine at 250 and the PMD-like feature at 298–300, with the length difference between donors taken up by one gap run after residue 272; CP134815.1 carries S at the reference tyrosine position. Colour denotes residue class, tint marks a residue that differs from Voyager-1 and a dash marks a gap. The bracket spans the region transferred into the Voyager-1 backbone, the union of the five exchanges (53–56 residues per construct, within positions 242–299). Over that span the donors are 20–29% identical to Voyager-1 and only R244, N247 and K263 are shared by all six proteins, yet all five constructs read −15/−14/−13 as T·G·C.

**Read depth.** Randomising twelve positions rather than nine raises library complexity 64-fold without a matching rise in sequences recovered, so information content scales with n and the two libraries are not on a common absolute scale — Voyager-1 reads −18 at 0.75 bits on the restricted library and 0.50 on the full one. Each construct is read against the Voyager-1 reference within its own library.

#### Voyager-1

27/30 exact

frame 2:27  3:3

| | |
|---|---|
| GTACGCAAC | 34 |
| GCACGTCCC | 32 |
| TCATGTAAC | 31 |
| ACATGTCCA | 30 |
| TTACGTCTT | 30 |
| ACACGCACC | 30 |
| GTACGGTCA | 29 |
| ATACGCACC | 29 |
| ACACGTTCG | 29 |
| ACACGTGCC | 29 |
| TTATGTACA | 28 |
| GGTACGTTC | 28 |
| GGTACGTCC | 28 |
| GTACGGTCC | 28 |
| TTACGTACA | 28 |
| TCACGTAAT | 27 |
| GCACGTACC | 27 |
| ATATGGTCA | 27 |
| ATACGTTCA | 27 |
| CTACGTACC | 27 |
| ATACGCTCT | 27 |
| ACACGTACA | 27 |
| GTATGTGAA | 27 |
| TCATGTACG | 26 |
| TTACGTATC | 26 |
| TCACGTCCG | 26 |
| TATACGCAA | 26 |
| TTACGTACG | 26 |
| GTACGTCCA | 26 |
| GTACGTGCG | 26 |

#### HNH12

21/30 exact

frame 2:12  3:18

| | |
|---|---|
| GTACGCACC | 107 |
| AAACGCCCA | 103 |
| GGTACGCCC | 94 |
| CATACGCCC | 93 |
| TATATGCCC | 90 |
| GTACGCCCA | 85 |
| AAAACGCCC | 84 |
| AATACGCAC | 81 |
| AATAGGCCC | 79 |
| TTACGCCCA | 79 |
| ATACGCACC | 79 |
| TTACGCCTC | 78 |
| AATACGGCC | 77 |
| GTACGGACC | 77 |
| GATACGTCC | 76 |
| TTAACGCCC | 75 |
| AATACGCCC | 75 |
| AGTAGGCCC | 75 |
| CTTACGCCC | 75 |
| GTACGCACT | 74 |
| AAAAGGCCC | 74 |
| TTTAGGCCC | 73 |
| GTACGTACA | 73 |
| GATACGTAC | 72 |
| TCTACGCCC | 72 |
| GTACGCATC | 72 |
| TTACGCTCC | 72 |
| AATACGTCC | 72 |
| GTTACGCCC | 72 |
| GTATGTCCC | 71 |

#### HNH10

16/30 exact

frame 2:12  3:18

| | |
|---|---|
| TATACGCCC | 89 |
| AGTACGCCC | 73 |
| GTACGTCCC | 66 |
| GGTACGCCC | 65 |
| TGTGCACCC | 65 |
| ATACGTCCA | 64 |
| AGTGCACCC | 61 |
| AAACGTCCT | 60 |
| TTACGTCCC | 60 |
| ATACGTCCC | 60 |
| TAAACGCCC | 60 |
| ATTACGCCC | 60 |
| ATACGCCCG | 59 |
| AATAAGCCC | 59 |
| TGTACGCCC | 59 |
| TATGCGCCC | 58 |
| GGTGCACCC | 58 |
| ATACGCACC | 58 |
| TATACACCC | 58 |
| GTACGTACC | 57 |
| AATACGTAC | 57 |
| GTACGCCCG | 57 |
| ATACGGACC | 57 |
| GATACGCAC | 56 |
| TATAAGCCC | 56 |
| AAACGCCCT | 56 |
| AGACGTCCC | 56 |
| AATATCCAC | 55 |
| TGTACGTCC | 55 |
| TATACACCA | 55 |

#### HNH14

15/30 exact

frame 2:5  3:25

| | |
|---|---|
| GGTACGCCC | 83 |
| GGTACACCC | 70 |
| AGTACACCC | 69 |
| TATACGCCC | 66 |
| AATAGGCCC | 64 |
| TATAAGCCC | 64 |
| TTACGTACC | 63 |
| GGTACGTCC | 62 |
| ATTACGCCC | 61 |
| TATACACCC | 61 |
| TGTACGCCC | 60 |
| CATACGCCC | 60 |
| TATAGGCCC | 60 |
| AGTACGCCC | 59 |
| AGTAAGCCC | 59 |
| TTTACGTCC | 58 |
| CTTACGTCC | 58 |
| AATACGCCC | 57 |
| CATACGTCC | 55 |
| GTACGTACC | 55 |
| TATAGCCAC | 55 |
| TGTAAGCCC | 55 |
| TTATCACCC | 54 |
| GGTAGGCCC | 54 |
| GATACGCCC | 54 |
| GTTACACCC | 54 |
| GTACACCTC | 54 |
| GTAAGCCTC | 54 |
| TGTATGCCC | 54 |
| TGTAGGCCC | 54 |

#### HNH15

20/30 exact

frame 2:13  3:17

| | |
|---|---|
| ATACGCCCA | 48 |
| TATAGGCCC | 47 |
| ATACGCCCG | 46 |
| TATACGCCC | 45 |
| ATACGTCCC | 44 |
| TATGCACCC | 42 |
| AATACGCCC | 41 |
| ATACGCACC | 40 |
| AATAGCCCC | 39 |
| GTACGTCCC | 38 |
| TATACACCC | 37 |
| ATACGCCCT | 37 |
| TATAAGCCC | 37 |
| GTACGTCCG | 37 |
| ATACGTCCG | 36 |
| AATAGGCCC | 36 |
| TATACGGCC | 36 |
| AATACGTCC | 34 |
| GTACGCCCA | 34 |
| AATACACCC | 34 |
| TATACGTCC | 33 |
| ATACGTCCA | 32 |
| ATACGTACC | 32 |
| TTACGCCCT | 32 |
| AATATGCCC | 32 |
| AAAACGCCC | 31 |
| TATACGCCA | 30 |
| TATATGCCC | 30 |
| GTACGCCTC | 30 |
| TATAGCCCC | 30 |

#### HNH16

14/30 exact

frame 2:0  3:30

| | |
|---|---|
| GGTACGCAC | 52 |
| TATACGGAC | 50 |
| TATACGTAC | 48 |
| TATAGGCAC | 45 |
| TATAGGTAC | 45 |
| GGTACGTAC | 43 |
| TATAGGCCC | 42 |
| GATACGCCC | 41 |
| TATAAGGAC | 41 |
| TGTAGGCCC | 41 |
| AATACGCAC | 40 |
| TATACGCAC | 39 |
| TGTACGTAC | 38 |
| GATACGCAC | 38 |
| AGTAAGCCC | 38 |
| TATACGCAG | 38 |
| TAAACGCAC | 37 |
| TGTACGGAC | 37 |
| TATAAGGAG | 37 |
| AATACGCAG | 36 |
| CATACGCAC | 36 |
| AAAACGCAC | 35 |
| TGTACGCCC | 35 |
| GGTAAGCAC | 35 |
| GGTAAGGAC | 35 |
| AATAGGTAC | 35 |
| TGTACGCAC | 35 |
| GATACGTAC | 34 |
| GGTACGGAC | 34 |
| GGTAAGGAG | 34 |


#### HNH16

14/30 exact

frame 2:0  3:30

| | |
|---|---|
| GGTACGCAC | 52 |
| TATACGGAC | 50 |
| TATACGTAC | 48 |
| TATAGGCAC | 45 |
| TATAGGTAC | 45 |
| GGTACGTAC | 43 |
| TATAGGCCC | 42 |
| GATACGCCC | 41 |
| TATAAGGAC | 41 |
| TGTAGGCCC | 41 |
| AATACGCAC | 40 |
| TATACGCAC | 39 |
| TGTACGTAC | 38 |
| GATACGCAC | 38 |
| AGTAAGCCC | 38 |
| TATACGCAG | 38 |
| TAAACGCAC | 37 |
| TGTACGGAC | 37 |
| TATAAGGAG | 37 |
| AATACGCAG | 36 |
| CATACGCAC | 36 |
| AAAACGCAC | 35 |
| TGTACGCCC | 35 |
| GGTAAGCAC | 35 |
| GGTAAGGAC | 35 |
| AATAGGTAC | 35 |
| TGTACGCAC | 35 |
| GATACGTAC | 34 |
| GGTACGGAC | 34 |
| GGTAAGGAG | 34 |


#### HNH3

26/30 exact

frame 2:23  3:7

| | |
|---|---|
| AATACGCCC | 66 |
| GTACGTACC | 62 |
| GTACGTCCC | 62 |
| GTACGGACC | 60 |
| GTACGTCCA | 60 |
| ATACGTCCA | 55 |
| GTACGCCCA | 55 |
| GTACGCACC | 55 |
| ATACGCACC | 54 |
| ATACGCCCA | 54 |
| TATACGCCC | 54 |
| TATACGTCC | 53 |
| TTACGTCCC | 52 |
| ATACGCCTC | 50 |
| ATACGTCCC | 50 |
| GTACGCACA | 49 |
| TAAACGCCC | 49 |
| GTACGTAGA | 48 |
| GTACGCCTC | 46 |
| GTACGCCAC | 46 |
| TAAACGTCC | 45 |
| GTACGCACT | 45 |
| ATACGGACC | 45 |
| GTACGTACA | 45 |
| ATACGCACT | 45 |
| ATACGCCCC | 44 |
| GTACGCCCT | 44 |
| TATACGCAC | 44 |
| ATACGCCTT | 44 |
| TGTACGTCC | 44 |

#### HNH1

19/30 exact

frame 2:16  3:14

| | |
|---|---|
| GTACGGACC | 65 |
| GGTACACCC | 63 |
| GTACACCTC | 62 |
| GTACGCACC | 62 |
| GTACGTACC | 61 |
| AGTACGCCC | 59 |
| TGTACGCCC | 58 |
| TATACACCC | 58 |
| TTACGTCCC | 57 |
| GGTACGCCC | 56 |
| TATAGCCAC | 55 |
| TTTACGCCC | 54 |
| GTACGTAGA | 54 |
| GTACATCCC | 53 |
| TATACGCCC | 53 |
| ATACGTCCC | 53 |
| CATACGTCC | 53 |
| ATACGTACA | 52 |
| AAACGTCCC | 52 |
| GTACGTCCC | 52 |
| TATACGTAC | 52 |
| TGTACGTAC | 52 |
| GTACGTGCA | 51 |
| GTAAGTCCC | 51 |
| TATAAGCCC | 50 |
| AGACGTCCC | 50 |
| GTACGTACA | 50 |
| TATATGCCC | 50 |
| ATACGTGCC | 50 |
| AGTACCCCC | 50 |

#### HNH17

20/30 exact

frame 2:15  3:15

| | |
|---|---|
| TGTACGTCC | 54 |
| GTATGCCCA | 54 |
| AAAACGCCC | 51 |
| GATACGTCC | 51 |
| TATACGCCC | 49 |
| AATACGCCC | 49 |
| GTACGCACC | 49 |
| AATACGTCC | 48 |
| TATACGTCC | 48 |
| TAACGCCCT | 48 |
| ATATGCCCA | 47 |
| AAACGCCCC | 47 |
| AAACGCCCT | 46 |
| TGTATGCCC | 45 |
| TATAGCCAC | 45 |
| ATACGCCCA | 44 |
| ATATGTCCC | 44 |
| TGACGTCCC | 43 |
| AAACGTCCC | 43 |
| GATATGCCC | 43 |
| GTACGTCCA | 42 |
| TTTACGCCC | 42 |
| TATACGCAC | 42 |
| AAACGGACC | 41 |
| AATATGCCC | 41 |
| AATACGCCA | 41 |
| TAACGTCCC | 41 |
| TTTATGCCC | 41 |
| GTATGCACC | 41 |
| TAACGCCCC | 41 |

#### HNH9

16/30 exact

frame 2:20  3:10

| | |
|---|---|
| GTACGCAAT | 67 |
| AAACGCACC | 65 |
| AAACGCACT | 63 |
| ATACGCAAC | 63 |
| GTACGCAAC | 61 |
| TATACGCAC | 60 |
| GTACGCACC | 59 |
| TATGCGCCC | 57 |
| AAACGTATC | 57 |
| AAAACGCCC | 54 |
| AAAACGCCA | 53 |
| ACACGCCAT | 53 |
| GTACGGACA | 53 |
| GTACGCCAC | 53 |
| GTACGCATC | 53 |
| TTACGCATC | 53 |
| GTACGCATA | 52 |
| GTACGCACT | 52 |
| TTACGGAGA | 51 |
| TATAGCCAA | 51 |
| GTTACGCCA | 51 |
| GTACGCACA | 51 |
| TAAACGCCC | 50 |
| AAAGCGCCC | 50 |
| CTACGCACC | 50 |
| AATAGCCAG | 49 |
| ATATGCACG | 49 |
| TTACGCACA | 49 |
| TAAGCGCCC | 49 |
| AAACGCACA | 49 |

Bases ACGT   black box, best-scoring YAYGY window (Y = C or T) in that read   G base departing from YAYGY inside the box (shown in a darker shade of its own colour)   frame 2 / 3, motif starting at the 2nd or 3rd position of the 9-mer

```
            CXXC Zn          HH                               C...C Zn

Voyager-1   CELCGKTNTIIHMHHVKRLKDIKNKPNKSFLEQRMVSRNRKTIALCK-----ECHVKRHKGE-V
HNH12       CEWCAVSGVPLEMHHIRKLKDLQGK---KKWEQRMIARRRKTMALCV-----SCHHDLHNGKLD
HNH10       CEYCGASDN-LEMHHVRKLKDLKGK---ELWEKHMIARKRKTLAVCY-----NCHVKIHNGKMD
HNH14       CEYCQATNSLIEIHHVRKLKDLKGK---TFWERLMISRQRKTIALCK-----DCHKKLHHGKLD
HNH15       CEWCGVENTPLEMHHVKKLKNLEGK---KRWEQFMIARNRKTIAMCV-----KCHHDLHNGKLD
HNH16       CDWCGTTDIPLEIHHVRKLKDLKGR---KSWERAMIARNRKTMALCANGHGNECHKKLHAGLLD
HNH3        CEMCGKKNLAIEIHHIRKLKDLKGK---KMWEKMMIARKRKTMALCH-----QCHIDLHSGKLD
HNH1        CEYCQATNSLIEIHHVRKLKDLKGK---TFWERLMISRQRKTIALCK-----DCHKKLHHGKLG
HNH17       CEWCGETKKPVEVHHIKKLKDLEGK---ARWEKVMIARRRKTMVLCV-----PCHHKLHAGKLD
HNH9        CELCGKTDCEIEIHHIRKLKDLKGV---NYWERFMIARNRKTLALCV-----DCHKKLHSGKLN
identical / ≥7 of 10
            C+ C+      + HH ++LK+++++     +E  M++R RKT ++C       CH ++H G+++
            .........|.........|.........|.........|.........|.........|....
                    10        20        30        40        50        60
```

Residue colours (ClustalX categories) K R A C I L M F W V N Q S T G P H Y D E grey, gap

**Figure S19. Exchanging the HNH domain relaxes the YAYGY recognition motif and shifts its register.** (A) The thirty most frequently recovered target sequences for Voyager-1 and for nine derivatives in which the HNH domain alone has been exchanged, from the CXXC motif to the C-terminus, leaving the upper linker as wild type. The ten constructs are shown as two blocks for legibility, with Voyager-1 repeated at the head of each so that both can be read against it. Each column lists the 9-mer recovered from the +3 to +11 randomised library with its read count, ordered by count. Bases are coloured A red, C blue, G gold, T green. The black box on each row marks the best-scoring YAYGY window in that read (Y = C or T), scored over the two frames the motif is known to occupy — starting at the 2nd or the 3rd position of the 9-mer — with the invariant A and G weighted most heavily. A grey-highlighted, bold base departs from YAYGY inside the box, and is drawn in a darker shade of its own colour so that it stays legible on the grey. Above each column is the number of the thirty reads matching YAYGY exactly and the split between the two frames.

**(B) The exchanged domains.** MUSCLE alignment of the segment that differs between the constructs: from the CXXC motif at residue 550 to the C-terminus, 55–61 residues over 64 aligned columns, coloured by ClustalX residue category. Bars mark the N-terminal CXXC, the paired histidines of the HNH catalytic motif, and the second, C-terminal pair of cysteines; the two cysteine pairs ligate the second, DNA-distal zinc and are present in all ten. Below the alignment a letter marks a column identical in all ten sequences and a plus sign a column shared by at least seven. The donors are otherwise highly divergent, so the constructs differ across most of the domain while retaining the catalytic and metal-binding framework. Voyager-1 alone carries a three-residue insertion (PNK) that none of the nine donors has, and HNH16 alone carries a five-residue insertion (ANGHG) between the two C-terminal cysteines — HNH16 being also the construct whose recognition register has shifted wholesale in (A).

**What the figure shows.** Voyager-1 is tight: 27 of its 30 best reads match YAYGY exactly and 27 sit in frame 2, so the boxes form an almost straight column. Every HNH exchange loosens this. The exact-match count falls to 14–26, and the register spreads across both frames or moves wholesale — HNH16 has all thirty reads in frame 3 and not one in frame 2. HNH3 is the most conservative exchange (26/30 exact, 23 in frame 2) and HNH16, HNH14 and HNH10 the least (14, 15 and 16 of 30). Across all 300 reads the departures are not evenly spread: the central A is almost untouched (8 departures) while the three pyrimidine positions and the G account for the rest (33, 37, 27 and 21), so the exchanged domains retain the core A·G contact and relax the pyrimidine preferences around it.

**Constructs.** Voyager-1 is Bac4 fix2 (B1). The nine exchanges are the Bac4_HNH<N>_repaired series, N = 12, 10, 14, 15, 16, 3, 1, 17 and 9 — the family in which only the HNH domain is replaced. Read counts are raw, not normalised to library composition, and differ several-fold between constructs; the figure compares motif composition within a column, never count magnitude between columns.
